# Strain tracking with uncertainty quantification

**DOI:** 10.1101/2023.01.25.525531

**Authors:** Younhun Kim, Colin J. Worby, Sawal Acharya, Lucas R. van Dijk, Daniel Alfonsetti, Zackary Gromko, Philippe Azimzadeh, Karen Dodson, Georg Gerber, Scott Hultgren, Ashlee M. Earl, Bonnie Berger, Travis E. Gibson

**Affiliations:** Department of Mathematics, Massachusetts Institute of Technology, Cambridge, MA, USA; Department of Pathology, Brigham and Women’s Hospital, Boston MA, USA; Infectious Disease and Microbiome Program, Broad Institute, Cambridge, MA, USA; Delft Bioinformatics Lab, Delft University of Technology, Delft, 2628 XE, The Netherlands; Computer Science and AI Lab, Massachusetts Institute of Technology, Cambridge, MA, USA; Department of Molecular Microbiology and Center for Women’s Infectious Disease Research, Washington University School of Medicine, St. Louis, MO, USA; Harvard Medical School, Boston, MA USA; Harvard-MIT Health Sciences and Technology, Cambridge, MA, USA

## Abstract

The ability to detect and quantify microbiota over time has a plethora of clinical, basic science, and public health applications. One of the primary means of tracking microbiota is through sequencing technologies. When the microorganism of interest is well characterized or known *a priori*, targeted sequencing is often used. In many applications, however, untargeted bulk (shotgun) sequencing is more appropriate; for instance, the tracking of infection transmission events and nucleotide variants across multiple genomic loci, or studying the role of multiple genes in a particular phenotype. Given these applications, and the observation that pathogens (e.g. *Clostridioides difficile, Escherichia coli, Salmonella enterica*) and other taxa of interest can reside at low relative abundance in the gastrointestinal tract, there is a critical need for algorithms that accurately track low-abundance taxa with strain level resolution. Here we present a sequence quality- and time-aware model, *ChronoStrain*, that introduces uncertainty quantification to gauge low-abundance species and significantly outperforms the current state-of-the-art on both real and synthetic data. ChronoStrain leverages sequences’ quality scores and the samples’ temporal information to produce a probability distribution over abundance trajectories for each strain tracked in the model. We demonstrate Chronostrain’s improved performance in capturing post-antibiotic *Escherichia coli* strain blooms among women with recurrent urinary tract infections (UTIs) from the UTI Microbiome (UMB) Project. Other strain tracking models on the same data either show inconsistent temporal colonization or can only track consistently using very coarse groupings. In contrast, our probabilistic outputs can reveal the relationship between low-confidence strains present in the sample that cannot be reliably assigned a single reference label (either due to poor coverage or novelty) while simultaneously calling high-confidence strains that can be unambiguously assigned a label. We also analyze samples from the Early Life Microbiota Colonisation (ELMC) Study demonstrating the algorithm’s ability to correctly identify *Enterococcus faecalis* strains using paired sample isolates as validation.

## Introduction

The human microbiome is involved in many aspects of human health and disease and exhibits a great level of diversity within and across host environments [1]. One of the most basic forms of analysis performed on any sample in a microbiome study is determining what bacteria are present and at what abundance. Although some applications call for coarser-grained taxa identification at the Operational Taxonomic Unit (OTU) or species level [2, 3], newer studies increasingly focus on more fine-grained resolution at the strain, or even Single Nucleotide Variant (SNV) level [4], including: (1) tracking *Clostridioides difficile* (*C. diff*) infection transmission events in Intensive Care Unit (ICU) patients through Single Nucleotide Polymorphism (SNP) calls [5]; (2) studying the role that individual genes play in infant gut microbial community development following antibiotic treatments [6]; and (3) detailed phylogroup analysis of *Escherichia coli* (*E. coli*) strains identified in longitudinal fecal samples from recurrent Urinary Tract Infection (rUTI) patients [7]. Since these studies try to draw conclusions about strain fitness, stability and/or competition, they rely on accurate quantifications of strains in time-series.

The process of converting bulk shotgun sequencing reads to taxa abundances usually involves some aspect of mapping or aligning reads to reference sequences [8–13]. An alternative is to perform metagenomic assembly [14], though for low-abundance taxa including most gastrointestinal pathogens of interest, this is unlikely to generate scaffolds of sufficient quality to produce reliable strain-level insights. Unfortunately, state-of-the-art methods quantifying strain-level abundances have a multitude of shortcomings when used to track low-abundance taxa, and these shortcomings become more evident when used to study longitudinal samples.

Only a select few methods report a statistic using the raw sample that can be directly interpreted as a strain’s predicted abundance. Instead, many approaches typically report *pile-up* statistics for SNPs across reference genomes or gene-specific loci [12, 15]. This then still leaves a large gap between pile-up information and the interpretation of results as strain dynamics in a longitudinal study. There are several methods that are specifically designed to de-convolve corresponding allele counts from pile-ups into abundances [9, 16]. However, per-locus pile-ups do not encode quality scores and no longer contain any information about SNV co-occurrences.

No existing method simultaneously leverages the temporal information in a longitudinal study design with the corresponding raw sequencing data. Indeed, base quality score information is often only used for pre-filtering low quality reads in bioinformatics pipelines [10, 17, 18]. Furthermore, no method to date utilizes the fact that multiple samples may have come from the same donor at different timepoints. Both sources of information can help overcome ambiguity when mapping or aligning reads. Notably, there are only a few methods that provide uncertainty quantification (e.g. confidence/credibility intervals) for abundance estimates. This is typically accomplished through Bayesian modeling, but previous methods take only a single sample as input. Meanwhile, recent works in other domains of computational biology have successfully demonstrated that such techniques can help overcome data sparsity and improve accuracy [19]. Furthermore, previous Bayesian methods that attempt to model read sequences to be more sensitive to SNVs are computationally burdensome [17].

To address these unmet needs, we developed ChronoStrain: an *uncertainty-aware*, time-series strain abundance estimation algorithm. It is, to our knowledge, the first fully Bayesian algorithm that fits all the above specifications. Chronostrain takes as input raw reads with associated quality score information and sample labels (host and time of collection) to output mixtures of strain calls that quantifies uncertainty in the strain abundances over time. Furthermore, Chronostrain works with any user defined set of marker genes for strain calls. To be precise, we define a *strain* as a cluster of subsequences from reference genomes, with a representative reference genome used to name it [15, 17, 20]. The threshold used to define these clusters is a user defined variable. In this work we use a range of thresholds from 0.998 similarity to just a single-nucleotide variant. We will use the words “strain cluster” and “strain” interchangeably throughout the rest of this text.

We demonstrate the superior performance of our algorithm on semi-synthetic data as well as data from two human studies. The first human study is the rUTI microbiome (UMB) project, a year long longitudinal study of women with a history of rUTI with a matched healthy cohort [7]. With the UMB study we focus on the increased utility and interpertability one has with Chronostrain compared to other state of the art methods when tracking strains over time. The second set of samples we apply Chronostrain to come from (what we are calling) the United Kingdom - Early Life Microbiota Colonisation (ELMC) study, which collected and sequenced between one and six fecal samples per subject (mean of 2.5) over the first few months of a child’s life (PRJEB32631, PRJEB22252) [21]. With the ELMC data, we demonstrate an improved lower limit of detection for Chronostrain for *E. faecalis* strains in fecal samples, using isolates from the original ELMC study to aid in validating our method.

## Results

### Overview of ChronoStrain

ChronoStrain, outlined in Figure 1a, consists of three main components: (1) a custom database that is constructed from user defined seed gene sequences, (2) a bioinformatics step for pre-filtering and processing raw reads, and finally (3) the core Bayesian inference algorithm. The software implementing ChronoStrain is written in Python and is available on Github (https://github.com/gibsonlab/chronostrain). ChronoStrain’s input is provided in three pieces: the raw fastq files from sequencing, a metadata file containing sample host and temporal information, and a database constructed from the FASTA-formatted *seeds*.

**Figure 1:**
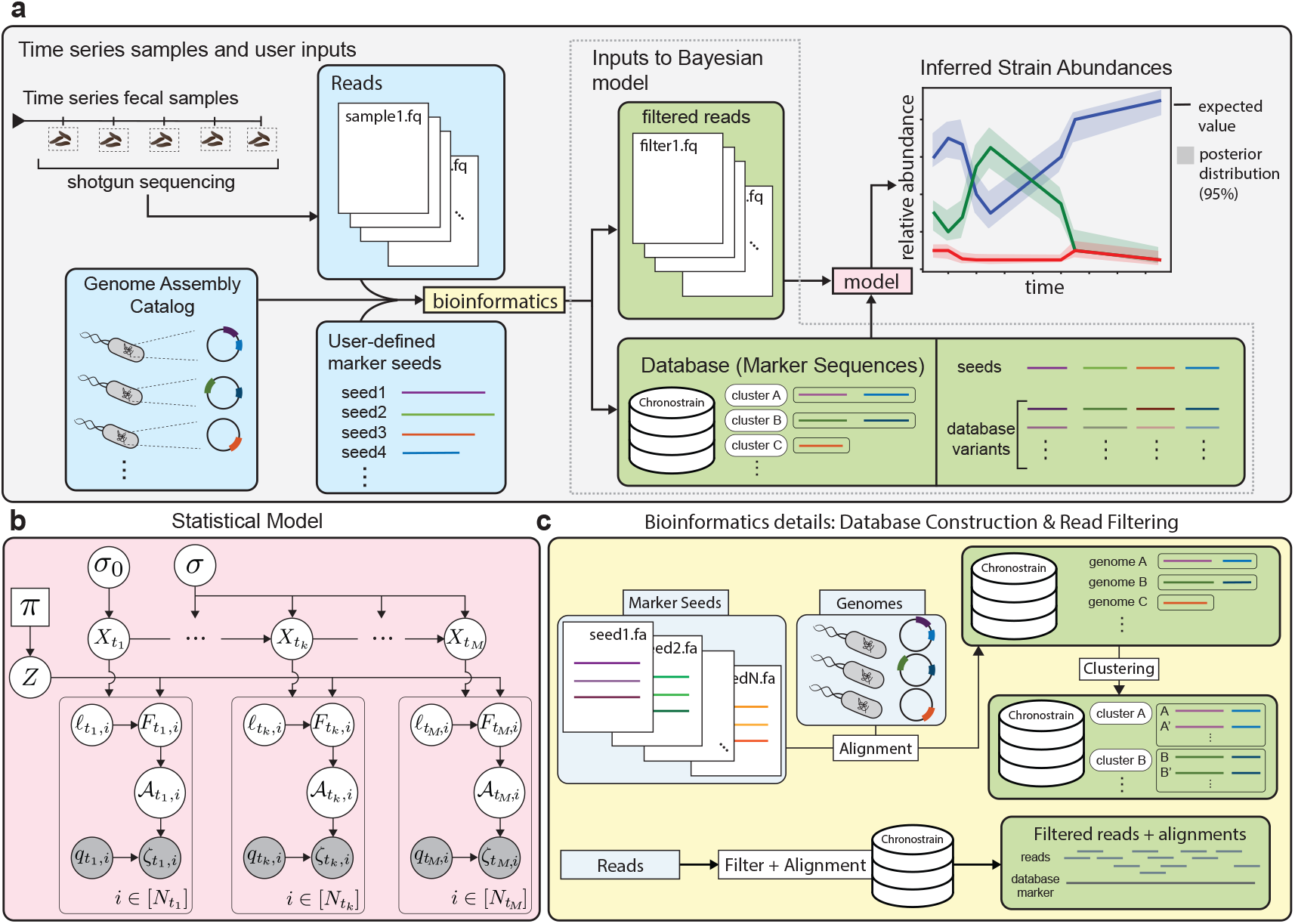
Overview of ChronoStrain. (**a**) A high-level schematic of the ChronoStrain pipeline. The pipeline constructs a database from marker sequence seeds, and uses it to filter reads to be passed as input to a Bayesian inference algorithm. As output, it returns a probability distribution over trajectories. (**b**) A graphical representation of the probabilistic model (Methods - Bayesian Model) used by ChronoStrain. White circles are latent random variables, gray circles are observations, squares are hyper-parameters (not all model parameters shown). (**c**) The “bioinformatics” step shown in panel (a) first generates a database representation of strains’ marker sequences; a thorough search is performed to account for all variants or similarities. A marker-similarity clustering is performed to address database redundancy. Only reads with high alignment identity to this database are kept.

Each strain in our model is specified via a collection of marker sequences, where a *marker sequence* is a subsequence of the genome that shares high sequence identity with a user-defined list of seeds. In turn, a *seed* is any sequence that the user specifies as key for distinguishing strains. We provide two example seed collections: one for *E. coli* and another for *E. faecalis* (Methods - Database Construction). These sequence seeds need not be genes core to the species, nor are expected to be single-copy, and thus may be chosen by the user depending on what they wish to quantify. One strength of our database’s design is that the marker selection can be guided by prior knowledge, which enables hypothesis-driven characterizations of strains. For example, one can use strain-specific genes that encode antibiotic resistance, fimbriae or toxin production which characterize pathogenicity; previous “marker” definitions often exclude these useful, interpretable regions [12].

A database of reference genome-marker combinations is constructed from a catalog of chromosomal assemblies (e.g. downloaded from NCBI’s assembly database) and high-identity hits to marker seeds via BLAST (Figure 1c); this functionality is provided by our software. ChronoStrain then clusters the references by a user defined level of marker sequence similarity, arbitrarily close to 100%. These two functionalities provide the ability to specifically target defined sequences/genes of interest, as well as the ability to differentiate reference sequences by whatever level of nucleotide identity the user wishes to use to specify “different” strains [22]. Finally, before passing the reads onto the core inference algorithm, our software filters out reads that do not align to our custom database beyond a user defined identity (default: 97.5%), which helps protect against contamination; see Figure 1c and Methods - Read Filtering for more details.

Our Bayesian model, for a single time series, is shown in Figure 1b. Strains are modeled using a stochastic process governing the latent taxa abundance vector 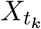 at time *t*_*k*_ (which is indexed by each strain *s* as 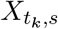), together with a model inclusion variable *Z* (which is also indexed as *Z*_*s*_). Then, at each time point *t*_*k*_, the *i*-th read is modeled as a nucleotide sequence 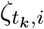 with its corresponding quality score vector 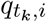. The sequence 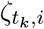 is modeled through the variables 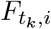 (the source nucleotide sequence fragment of the read), 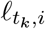 (the random length for a sliding window along the markers that determines which fragment is measured) and 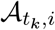 (the fragment-to-read substitution/indel error profile). A complete description of the model can be found in Methods - Bayesian Model.

Our model is closely related to a time-series topic model [23] with an extra component that models measurement noise. The connection is best drawn using an analogy: strains are *topics*; each abundance profile is a strain/topic mixture that produces a bag of fragments (*words*), whereby each fragment/word is measured with some quantifiable error per nucleotide (words are observed with mistakes). Both the time-series topic model and our model account for dependencies across time using a latent Gaussian process, transformed into mixture weight vectors of topics/strains via a softmax function.

To achieve scalability for our Bayesian inference (a limitation in some previous works that estimate Bayesian posteriors, see Supplement §A.4), we implemented Automatic Differentiation Variational Inference (ADVI [24]) as opposed to slower sampling algorithms (especially for large models such as ours) like Markov chain Monte Carlo. ADVI is a gradient-based optimization method that is easily implementable on a graphics processing unit (GPU). For our model, inference was further accelerated by compressing a component of the likelihood function into a more efficiently computable one (Supplement §B.1).

ChronoStrain outputs a posterior distribution over abundance *trajectories* across database strains. This is in contrast to algorithms which output point estimators (for instance, the most likely trajectory calculated via Expectation-Maximization). Specifically, in the context of longitudinal studies, ChronoStrain offers two big advantages in terms of interpretability. First, the user no longer has to stitch together outputs of an algorithm that has been run on each timepoint’s sample independently. ChronoStrain is capable of jointly modeling an entire longitudinal dataset from a single host, and thus produces abundance estimates which are more consistent throughout time than existing methods. Second, our posterior distributions offer visualizations of uncertainty that illustrate when labels are possibly ambiguous (suggesting that the sample contains a strain with a novel combination of SNVs) or if coverage is too small when the variationally-fit variance is large.

### ChronoStrain outperforms other methods in semi-synthetic experiments

For benchmarking, we created a *semi-synthetic* dataset that combines *in silico* (simulated) reads with real experimental data [15]. The *in silico* reads were created as follows: six Phylogroup A strains were selected at random, then ∼ 0.2% of the bases from those strains were randomly changed, so that ground truth strains are not identical to strains in the reference database. Then, noisy reads were simulated from those *in silico* mutant reference strains. The simulated reads were then combined with the first six timepoint samples from UMB18 which lack detectable phylogroup A strains^1^; these samples only contain phylogroup B2 and D *E. coli* strains. With this setup, we have a ground truth subset of six strains from phylogroup A that are not identical to strains in the database and where the algorithms must also contend with other real experimental reads from a complex microbial background that includes distantly related *E. coli*. The above process of *in silico* strain selection was repeated 10 different times; for each set of strains we simulated two distinct sets of random reads per read depth, resulting in 10 × 2 × 5 = 100 different read sets for benchmarking.

For comparison, we included StrainGST [15], StrainEst [20] and the mGEMS pipeline [25]. For a discussion on methods that did not make it into the benchmark, refer to Supplement §A. StrainGST identifies clusters by performing fast comparisons between read *k*-mer counts and *k*-mer counts from a database of strain genome assemblies. StrainEst deconvolves allele frequencies (generated from pile-ups against a small collection of reference genomes) into abundance estimates by setting up a large linear regression problem. mGEMS first performs *pseudo-alignments* using Themisto [26], then performs point estimation for a Bayesian model of Themisto’s outputs using mSWEEP [27], which outputs a modal posterior estimate assuming a Dirichlet prior on abundances. To ensure all metrics were on equal footing, each database was tuned so that the number of phylogroup A strain clusters were roughly equal. Note that mGEMS’ output is meant to be post-processed using `demix_check` [28]. However, the scores were unusable due to the mismatch between the binned reads and the nearest reference genome (see Methods - Semisynthetic).

We measured the algorithms’ performances using root-mean-square error (RMSE) of log abundance estimates (RMSE-log), area under receiver operating characteristic (AUROC) for detecting the *in silico* strains out of phylogroup A, and runtime (Figure 2). The RMSE-log was calculated in two ways: (1) normalized across the six strain clusters that coincide with the ground truth strains, and (2) normalized across the ∼340 Phylogroup A strain clusters.

**Figure 2:**
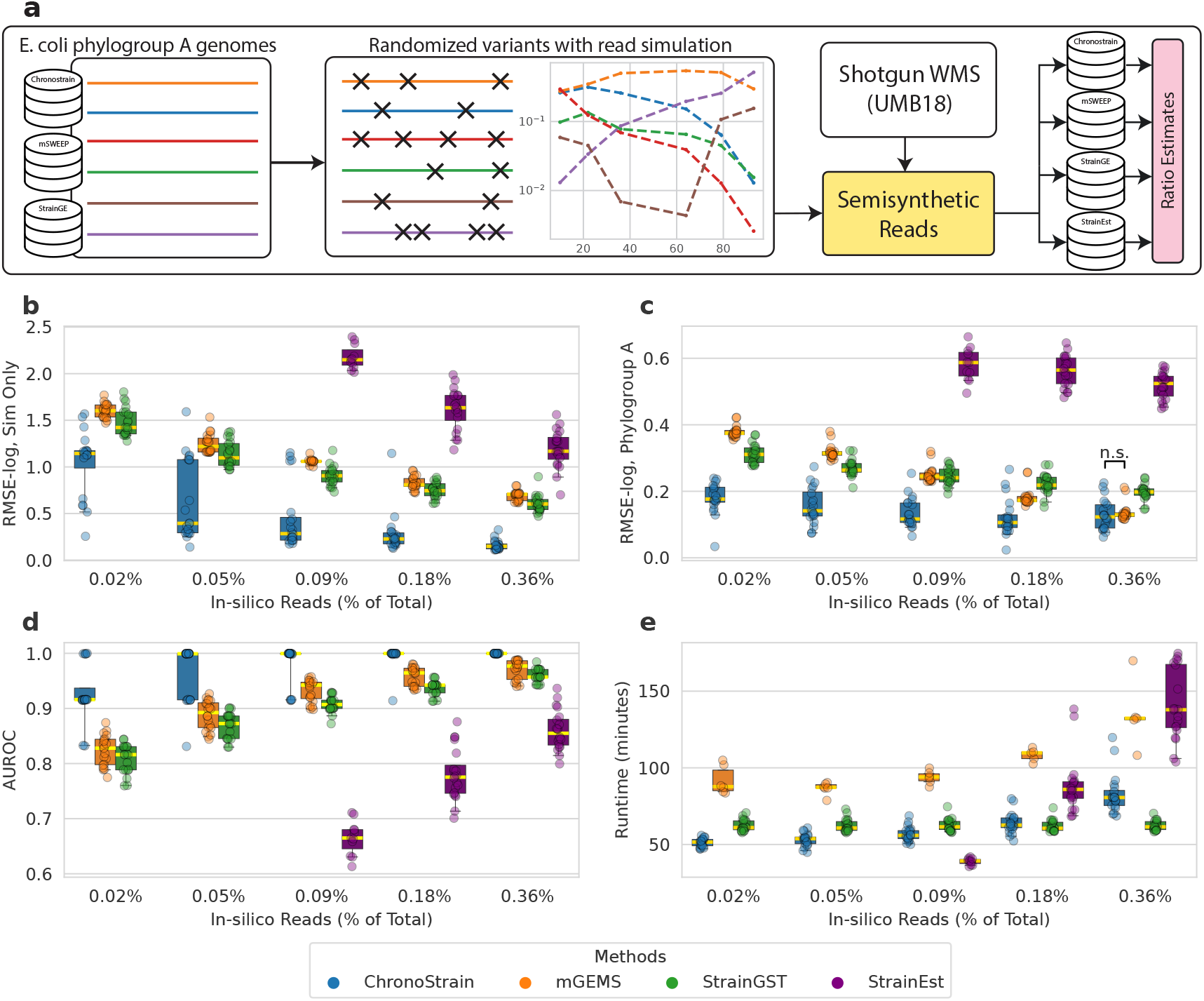
Chronostrain outperforms other state of the art methods with semi-synthetic data. All comparisons to Chronostrain in (b,c,d) are statistically significant at level 0.05, after two-sided, paired Wilcoxon tests with Benjamini-Hochberg (BH) correction, unless otherwise noted with an n.s. (**a**) Semisynthetic Data construction: Six phylogroup A genomes were selected from our genome database. Those genomes were then mutated, from which reads were generated with synthetic read counts 2500, 5000, 10k, 20k, 40k (0.02%, 0.05%, 0.09%, 0.18%, 0.36% of total reads, combined with UMB18 samples). The time-series trajectory is fixed across the 20 experimental replicates. We evaluate RMSE for model log-predictions **(b)** averaged over the six synthetic strains, and **(c)** averaged over all phylogroup A strains in the database which include false positives from the background (real) reads. (**d**) AUROC for synthetic strain detection normalized over phylogroup A. (**e**) Comparison of algorithm run-times.

ChronoStrain significantly outperforms all other methods for all simulated read depths in terms of RMSE and AUROC, with the exception of one scenario, all while maintaining a comparable runtime to the other methods (Figure 2, Supplemental Data Table 1). Note that StrainEst was unable to identify any of the *in silico* strains when 0.05% reads or less were sampled from them. With the RMSE separated across the six target strains, the performance trends were similar to the combined error metrics just discussed, with Chronostrain significantly outperforming the other methods for a majority of the read depths (Supplemental Figure S1, Supplemental Data Table 2). By binning the RMSE contributions according to the synthetic strain’s sample abundance into four equal-width bins, one begins to see a distinct significant advantage for Chronostrain in the lower-abundance bins (Supplemental Figure S2, Supplemental Data Table 3).

A sensitivity analysis on free design parameters was performed for the top three performing methods: Chronostrain, StrainGST, and mSWEEP (Supplemental Figure S3, Supplemental Data Table 4). For Chronostrain, we varied the prior probability for strain inclusion in the model *π*, where *Z*_*s*_ ∼ Bernoulli(*π*), and the threshold 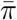 on the posterior *p*(*Z*_*s*_ |*R*) determining whether *s* is given a zero or nonzero prediction. For StrainGST, we varied the `score` parameter which is used as the metric for determining if a strain should be counted as present or not. For mGEMS, the free design parameters we varied were the beta-binomial mean hyperparameter *µ*_*β*_ (“*π*_*k*_” in the paper [27]) that models pseudoalignment counts, and the post-inference abundance threshold *ε*_LOD_ for filtering out false positives (see Methods - Semisynthetic).

For Chronostrain, varying the prior probability by several orders of magnitude, *π* ∈ {10^−4^, 10^−3^, 10^−2^, 10^−1^, 0.5} was largely insensitive and remained significantly better with the three posterior probabilities we tested 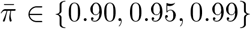. Decreasing mGEMS’ *ε*_LOD_ threshold did not affect the RMSE-log when calculated across the six *in silico* strains, but the RMSE-log increased when computed over all Phylogroup A predictions, suggesting that mGEMS is less effective at distinguishing low abundance from noise. The mean parameter *µ*_*β*_ had a similar, but weaker effect. StrainGST performed best when `score` was set to zero. Finally, we tested the sensitivity of the methods to varying levels of *in silico* strain mutation rates (relative to the reference strain in the database) testing mutation rates of 0.2%, 0.6%, 1.0%, 1.4%, and 1.8% and Chronostrain significantly outperformed all other methods in this experiment as well (Supplemental Figure S4, Supplemental Data Table 5).

### Analysis of UMB dataset with ChronoStrain provides interpretable results with more consistent correlations over time

The UMB project monitored 31 women in two cohorts, “rUTI” (multiple UTIs in past year) and “healthy” (no recent history of UTI), over the course of a full year [7]. Each participant provided a stool sample once a month, with outgrowth cultures grown from rectal and urine samples taken at the first month for all participants. For those participants who were clinically diagnosed with a UTI, additional urine samples and outgrowth cultures were taken on the days of diagnoses when possible. Beyond this, metadata about participants’ self-reported dates of last known antibiotic administration and the dates of infection are available. In addition to the original samples, we have added a new data modality. For specific samples from the rUTI cohort for which blooms were identified by the original StrainGST analysis [7], cultures from stool samples plated on MacConkey agar (favoring Gram-negative bacteria including *E. coli*) were sequenced. We applied ChronoStrain and StrainGST to all 31 time-series in the UMB study (Supplemental Data Trajectories U1-U31) with model outputs for UMB participant 18 shown in Figure 3. See Methods - UMB Analysis for details on the analysis pipeline.

**Figure 3:**
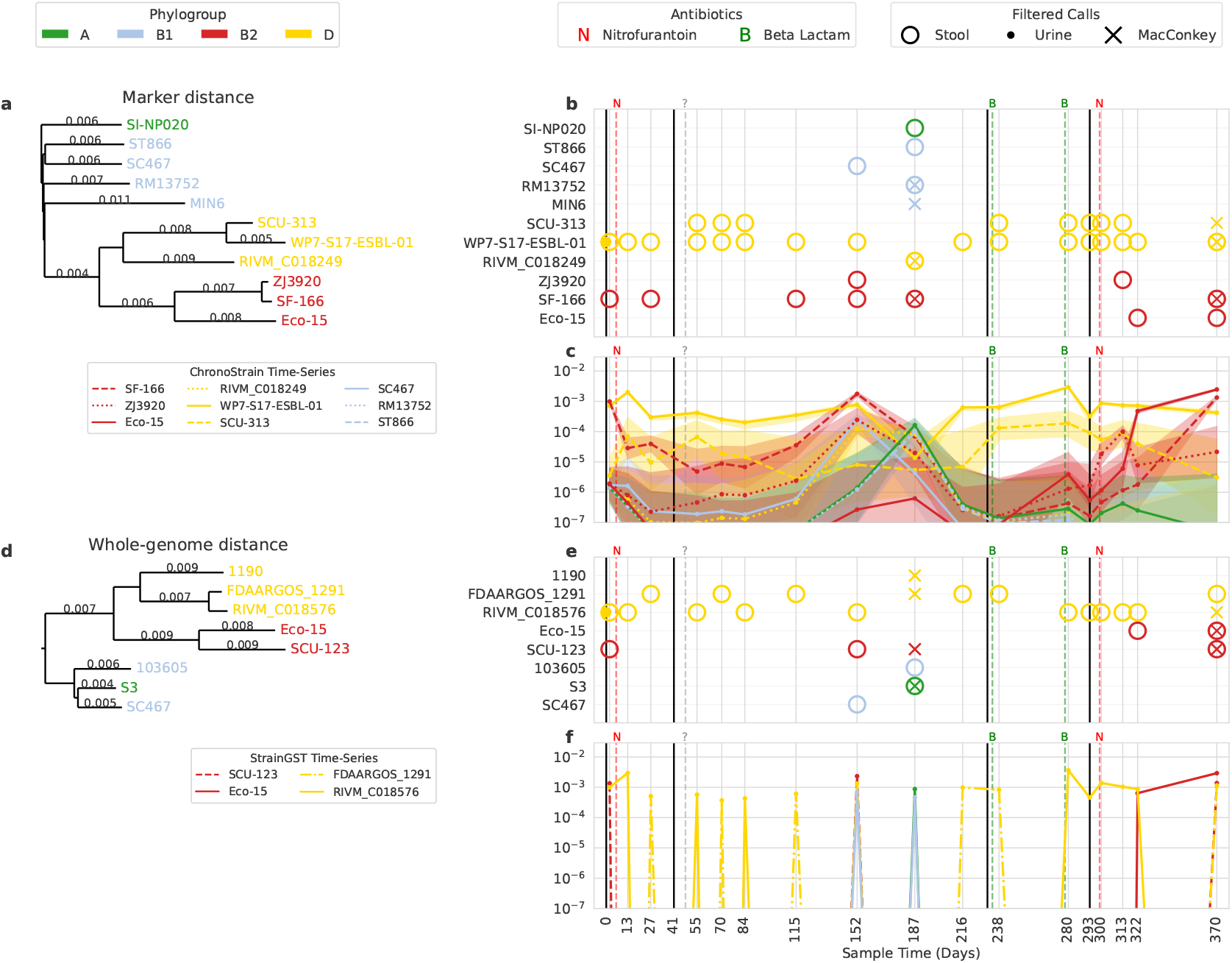
A visualization of ChronoStrain’s and StrainGST’s outputs for UMB18, a rUTI-positive participant. **(a**,**d)** Phylogenetic sub-trees of strains, computed using two different metrics: marker-specific k-mer proportion distance, and whole-genome k-mer Mash distance. **(b**,**e)** A scatterplot of strain detections across timeseries for the two respective methods. Different markers indicate sample modality (Stool, MacConkey culture from stool, urine). Solid vertical lines indicate dates of UTI diagnoses, dotted lines indicate self-reported, last-known dates of antibiotic administration. **(c**,**f)** Plots of estimated *overall* relative abundances in stool. Credible intervals are calculated using 5,000 posterior samples.

The output of ChronoStrain (Figure 3b,c) suggests that the initial infection most likely came from a Phylogroup D strain (closed circle on day 0) whose marker signature is given by cluster `WP7-S17-ESBL-01`. After multiple rounds of antibiotics, the phylogroup D strains (yellow) are no longer detectable in the urine but still persist in the GIT. Two of the Phylogroup D clusters are called across multiple timepoints (`WP7-S17-ESBL-01`: solid yellow line, and `SCU-313`: dashed yellow line), where the dominant cluster is the same as the one called in the urine sample and is also the most abundant strain in 13 of 17 timepoints. Another prominent cluster is `SF-166` (dashed red line) of phylogroup B2, which shows differing responses to the antibiotics; this cluster is re-capitulated in both of the enriched MacConkey-culture samples (x’s in Figure 3b with posterior abundance estimations in Supplemental Figure S5). The initial dose of nitrofurantoin and the unknown antibiotic reported by the participant before day 55 fail to clear this strain from the GIT; it is present with abundance above or near 10^−5^. Around the time of the third and fourth round of antibiotics, which were beta-lactam inhibitors, all the B2 reference strains’ abundances drop well below 10^−6^. However, the fifth round of antibiotics is a re-dosage of nitrofurantoin near day 300, for which ChronoStrain identifies the rapid growth of both the old SF-166 cluster but also `Eco-15` (solid red line) which is a newly dominant strain but was previously undetected. These results suggest beta-lactam more effectively cleared the B2 taxa in the GIT than nitrofurantoin. This is consistent with prior literature [29, 30] suggesting nitrofurantoin has higher host bioavailability, and thus accumulates less in the GIT in comparison to beta-lactam.

Interpreting the output of StrainGST (Figure 3d,e) one sees adjacent timepoints call two phylogroup D (yellow) clusters `FDAARGOS_1291` and `RIVM_C018576` in a mutually exclusive manner. This time-series/temporal inconsistency (alternating presence absence over time) is something that is absent in the joint analysis done by ChronoStrain. Furthermore, the fecal sample analysis on day 187 suggests that `SCU-123` is present in previous samples but not on that date, yet the MacConkey culture sample on that same timepoint suggests otherwise. Indeed, in ChronoStrain’s analysis the corresponding dominant B2 cluster `SF-166` was still detected on that particular date – which suggests that our method’s joint analysis had the correct detection call. The lack of a coherent cross-sample consensus makes it difficult to evaluate the sensitivity of different strains to antibiotics or to determine the presence of new strains from a bloom. Furthermore, the lack of a credibility (or confidence) interval hampers the interpretability of StrainGST.

In the analysis of UMB18 just presented, we defined distinct strain clusters as those with less than 0.998 nucleotide identity over the database marker sequences. This choice in threshold was so as to coincide with StrainGST’s level of nucleotide identity used to define clusters in their original work [7]. To demonstrate the utility as well as interpretability of our method we performed the same analysis as above (UMB18) but with reference genomes divided into distinct strains for any single nucleotide difference over our markers, Supplemental Figure S6. The overall story from before is the same: there is a dominant Clade D strain, a Clade B1 (or C) strain blooms in the middle, and a second B2 strain becomes dominant. With this finer resolution view, however, we do call more strain clusters. One can however directly see that several of the strain’s credible intervals are entirely overlapping. This suggests that the model is having trouble differentiating those strains, and that they should not be considered to be distinct. Practically, one might consider collapsing those strains, which we would have done had we not already started with a coarser strain clustering threshold in our direct comparison to StrainGST.

In order to quantify ChronoStrain’s qualitatively observed increased temporal consistency in strain detection compared to StrainGST, we computed the time-lagged Spearman correlation (the abundance rank correlation between adjacent timepoints) for all *E. coli* strains across all participants in the UMB study, Figure 4.^2^ For the top four most abundant phylogroups (A, B1, B2, D), ChronoStrain produced a significantly higher time-lagged correlation coefficient (medians typically at or above 0.75) when compared to StrainGST’s outputs (approximately 0 correlation and sometimes even negative), Figure 4a. The other phylogroups (C, E, F, G) were much rarer and thus were not statistically significant. Previous literature has used temporal correlations to demonstrate that a healthy adult microbiome is stable^3^ over time at the Operational Taxonomic Unit (OTU) level [32]. More recent work has shown that many of the macro-ecological properties seen at the OTU and species level (like stability) also extend to the strain level [33]. Thus, it should not be a surprise to find positive temporal correlations for *E. coli* strains in the UMB cohort. Finding negative temporal correlations, on the other hand, would be.

**Figure 4:**
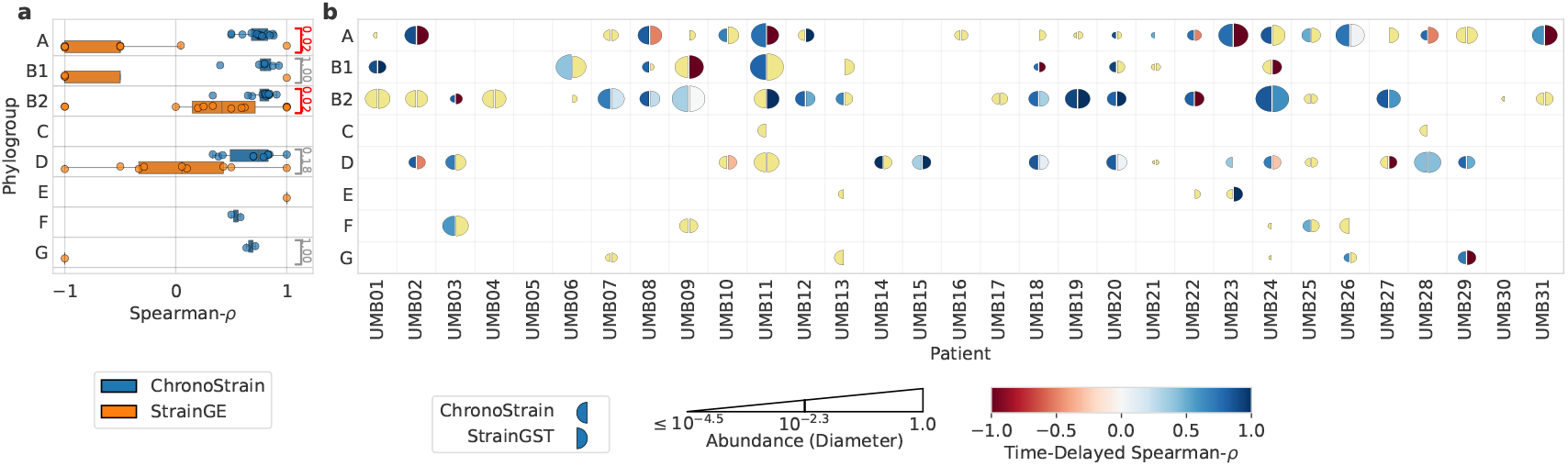
ChronoStrain provides time-consistent predictions. To quantify the improvement in the consistency across time compared to state-of-the-art predictions, we computed the time-delayed, phylogroup-specific correlation *ρ* for all participants (Spearman-*ρ* is computed for each pair of adjacent timepoints, then averaged across time). For ChronoStrain, we calculated *ρ* for the median trajectory of the posterior. (**a**) By aggregating across all participants, we performed paired, two-sided Wilcoxon tests for each phylogroup. Significance (red brackets) was determined via BH correction at a 5% false discovery rate; adjusted p-values are denoted. This result suggests that StrainGST’s output strains from phylogroups A, B1, B2, D are inconsistent across time (**b**) For each phylogroup, we plotted *ρ* and maximal abundance estimates for each participant. The size of each semicircle depicts a log-scale abundance level for the corresponding method, relative to the entire sample (details in Supplement), while the color depicts *ρ*. Yellow semicircles indicate NaNs.

### ChronoStrain calls isolates from infants with more accurate abundance estimation and is more robust to database mismatch

The ELMC study collected and sequenced longitudinal fecal samples from 596 full term babies during the neonatal and infancy period, with additional paired samples from a subset of the mothers [21]. Each infant in this study had between one and six fecal samples collected with a majority of the neonatal samples taken on days 4, 7, and 21. In this study, 805 isolates were obtained from fecal samples of 189 infants, of which 349 isolates were *E. faecalis* (321 from infants, 28 from mothers). We applied the mGEMS pipeline as well as ChronoStrain to the subset of 189 infants’ time-series fecal samples with a database that incorporated the isolate genomes.

The database for ChronoStrain was constructed with a collection of 2,026 European reference genomes [34] and all 349 ELMC *E. faecalis* isolate genomes, just like the mGEMS analysis in [28], plus additional genomes from *Enterococcaceae* (see Methods - ELMC Analysis). The genomes were then clustered at 0.998 similarity, the same as in previous sections. We also re-ran the entire mGEMS pipeline, meaning that (1) we first ran a species-level analysis to bin reads into the *E. faecalis* species (using a large 640,000-genome index [35]), then (2) we used an index constructed from the same 2,375 (2,026 + 349) *E. faecalis* genomes for the strain-level analysis. To ensure that both methods would be performing inference at the same level of strain granularity we manually tuned PopPUNK [36] (used by mGEMS for clustering) so that the number of strain clusters containing ELMC isolates was the same for both methods.

After ensuring that both methods’ databases were on equal footing, we performed inference on the infant fecal metagenomic samples. For mGEMS, we used the same hyper-parameters as described in [28] with *ε*_LOD_ = 0.01 and only reported calls with `demix_check` score 2 or better (smaller). To compare the methods at roughly equal sensitivities, we tuned ChronoStrain’s post-inference threshold so that the two methods reported the same number of infant *E. faecalis* isolates, settling on a posterior threshold 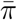 = 0.95 and only retaining clusters with *E. faecalis* abundance ratio at least 0.065 (Figure 5).

**Figure 5:**
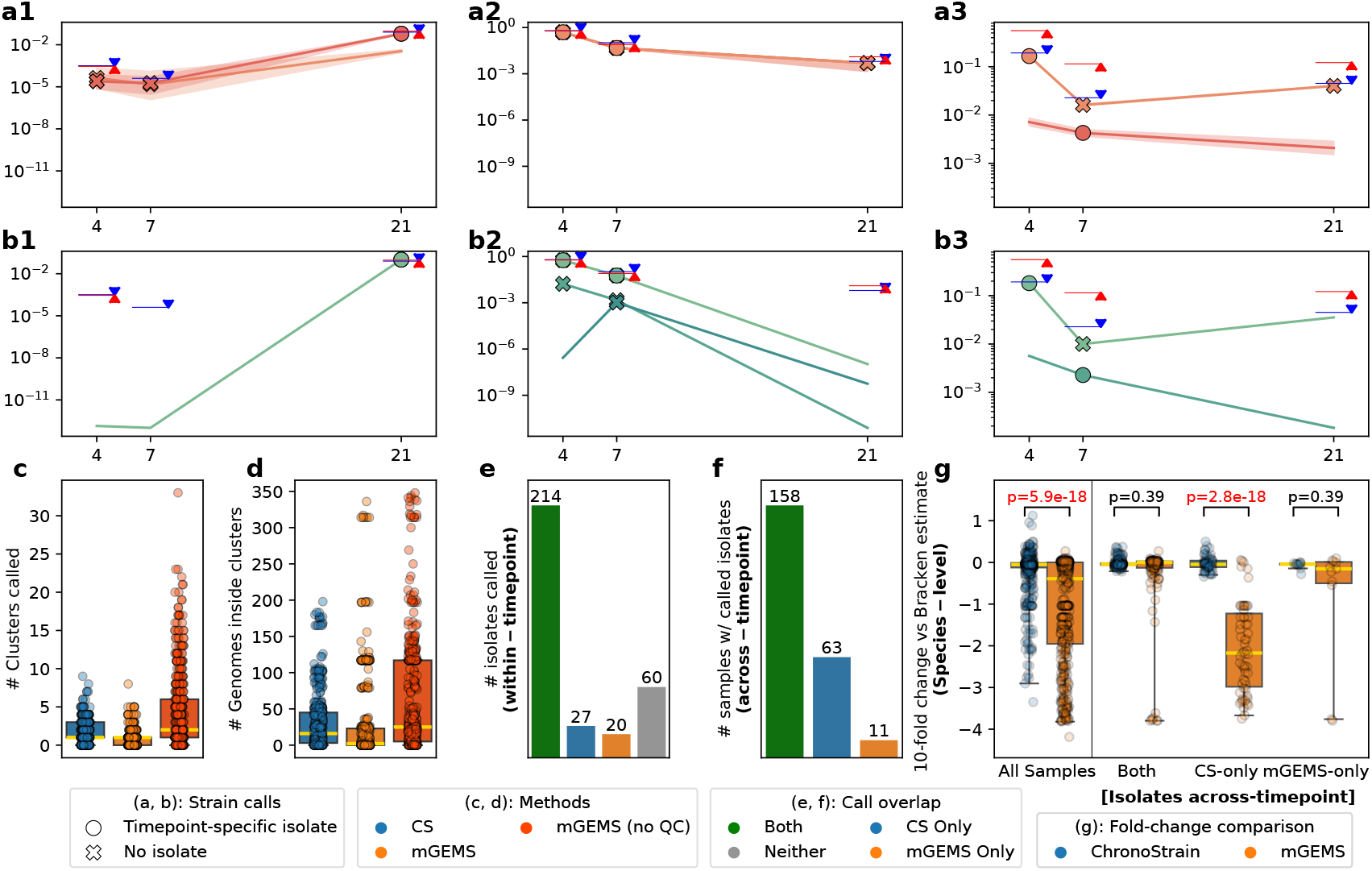
ChronoStrain calls isolates from infants across time with more accurate abundance estimation than mGEMS. **(a1-a3)** Example ChronoStrain *E. faecalis* strain abundance estimates for infants `A01077`, `B00053`, `B02273`. **(b1-b3)** mGEMS estimates for the same infants. For each cluster, we drew its trajectory only if it passed the method’s respective filter in at least one sample. Each timepoint-specific strain call is denoted, depending on whether the cluster contains an isolate culture from that timepoint (O) or not (X). Blue triangles + horizontal lines are Bracken *E. faecalis* species abundance estimates, Red triangles + lines are MetaPhlAn4 species estimates. **(c, d)** Number of clusters passing the filter and total genomes within those clusters. **(e)** Number of strain calls with a paired isolate in that same sample (321 total). **(f)** Number of strain calls with a paired isolate in a *different* sample from the same time series, counting cross-timepoint predictions. **(g)** For each category from (f), we checked how far the species predictions are from Bracken’s. Wilcoxon test (paired, two-sided and BH-corrected) p-values above brackets.

As intended, the number of strain calls per sample (Figure 5c) as well as the number of strain clusters corresponding to a cultured isolate from the same time-point (Figure 5e) were similar for both methods. However, we did notice a stark difference in the abundance estimates. We illustrate this with example trajectories from three infants `A01077`, `B00053`, and `B02273` in Figure 5a,b (complete set in Supplemental Data Trajectories E1-E21). To provide an independent comparison, we used Kraken2+Bracken (KB) [37, 38] and MetaPhlAn4 [39] to estimate *E. faecalis* **species** abundances (triangles in 5a,b) which yield approximately the same estimates for almost all samples. mGEMS often produces under-estimates relative to KB across the ELMC infant dataset (5g, “All Samples”), with the largest discrepancy between KB and mGEMS occurring for those samples where ChronoStrain makes a strain call to a paired sample isolate but mGEMS does not (“CS-only”). To better understand this discrepancy, we plotted *E. faecalis* abundance fold-changes relative to KB (and MetaPhlAn) for each sample (Supp. Fig. S7). At approximately 0.01 relative abundance and below^4^, mGEMS readily outputs zero (or near-zero) *E. faecalis* abundances, unlike KB, MetaPhlAn and ChronoStrain.

Finally, we tested the robustness of the methods when the reference database no longer contains genomes identical to the strains we are trying to track. Having the isolate genomes gives us the opportunity to perform an experiment where we mutate isolates used in the database (similar to the semisynthetic setting, but mutating the database instead of the reads) while still maintaining a notion of ground-truth strain calls. Specifically, we mutated 117 of the ELMC isolates, chosen from those already called by mGEMS (Supplement §B.6.1), and then performed inference with both methods using the same hyper-parameters and thresholds as before. The results from that experiment (Fig. 6 and Supp. Fig. S9, discussed in more detail in Supplement §B.6.2) show that mGEMS calls decreased from 117/117 to only 45/117 strains, but ChronoStrain’s results largely stayed the same (108/117 to 109/117). We emphasize that we intentionally chose isolates with a paired sample that already had an mGEMS call, regardless of ChronoStrain, which is why we report 108 calls for the original run instead of the full 117 for CS. The drop in the number of strain calls by mGEMS is due to an expected behavior: the `demix_check` scores became worse (increased) with mutated isolates in the database; see Figure 6b. One can increase the number of correct strain calls by allowing larger `demix_check` scores, but this comes at the cost of specificity. With the `demix_check` score threshold increased from 2 to 4 mGEMS correctly calls 93/117 of the strains, but the median number of calls per sample becomes six times larger than ChronoStrain (Supp. Fig. S9c).

**Figure 6:**
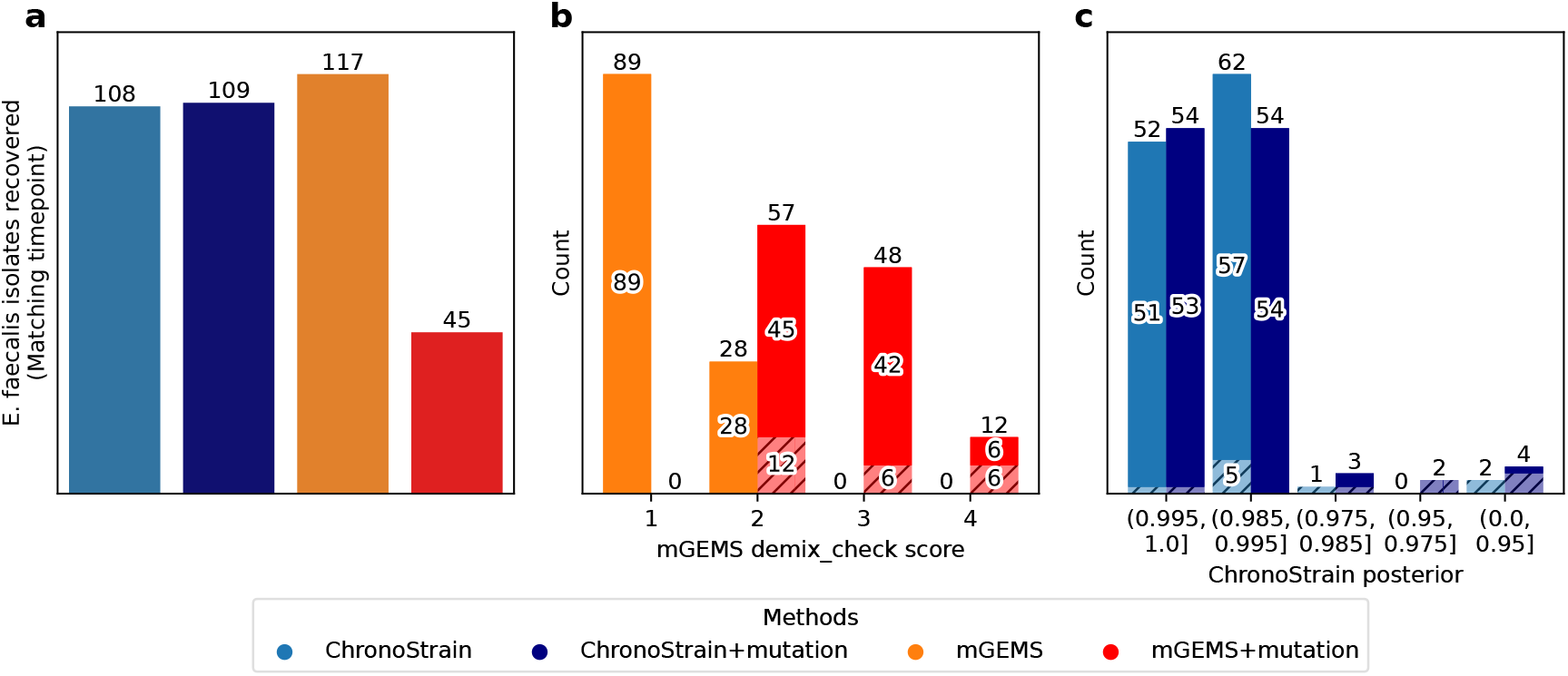
Chronostrain more robust than mGEMS to mismatches between database genomes and sample reads. We performed inference on the ELMC data where 117 of the ELMC isolates were mutated (genome mutation rate 0.002) before including them in the database (Supplement §B.6.1). **(a)** The raw number of isolate clusters called by each method (filters used for a call - ChronoStrain: 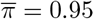, ratio ≥ 0.065; mGEMS: `demix_check` score 1 or 2, ratio ≥ 0.01). Note that mGEMS calls more isolates due to the experimental design: the 117 isolates were initially chosen using mGEMS predictions, even if ChronoStrain didn’t call them. **(b)** We show the `demix_check` score distribution for all 117 isolate clusters; “1” is best, “4” is worst. Each bar is divided into two sections with the solid upper region those strains with an abundance ratio ≥ 0.01 and dashed lower region those strains with abundance ratio < 0.01. The mutated genomes caused a precipitous increase in the `demix_check` scores. **(c)** ChronoStrain’s posterior probabilities, solid upper region are those strain calls with an abundance ratio ≥ 0.065 and dashed lower region those strain calls with abundance ratio < 0.065.

These experiments demonstrate that ChronoStrain has more accurate abundance estimates for lower abundance strains, and that it is more robust to database discrepancies when making strain calls (without losing specificity). mGEMS is sensitive to having the strain of interest being isolated and sequenced for database inclusion (Figure 6b) and the pipeline does not reliably estimate abundances below 0.01 (Supp. Fig. S7). These differences can affect how the strain dynamics are interpreted in statistically significant ways. For instance, when looking at strain turnover^5^ within *E. faecalis*, ChronoStrain estimates that twice as many infants had at least one turnover occurrence in the first month compared to mGEMS (40/189 versus 19/189, χ^2^-test *p* = 0.0046, Supplemental Data Table 6).

## Discussion

There are two major differences between ChronoStrain and comparable methods. First, ChronoStrain outputs a distribution over strain abundance trajectories, while the others provide a single point estimate. Since ChronoStrain is a full end-to-end Bayesian model of the reads’ errors across multiple samples, it explicitly propagates measurement uncertainty and temporal correlation. This allows for a qualitatively different approach to interpreting results with ChronoStrain, making it possible to directly visualize uncertainty through the plotting of credible intervals or to possibly use the entire posterior distribution in downstream analyses. It also helps quantify the uncertainty induced by the database; for almost all real data applications, the database will not contain the full catalog of strains that encapsulate all strains in the samples. Without isolates incorporated into the database, there will always be discrepancies between the nearest strain in the database and the genomic variants present in the sample.

Second, our method specifically tracks variants and combinations of genes, and so has applications where one wants to type strains beyond conventional genes (serotyping, sequence typing, or species core genes). In this vein, our method makes a very purposeful trade-off: we perform more computation per read to account for nucleotide *and* presence/absence variability, but in return, we gain (1) accuracy for low-abundance (low-coverage) strains for which assembly might not be possible and (2) a more customizable database that one can use to define strains via interpretable features. In our example databases, each genome’s markers form around 1.5% of the median species-level genome length. It may be surprising/counter-intuitive that the performance improvements seen in the benchmarks are being achieved with a far smaller database than other methods. However, the insight that “compressed” views of the genome (e.g. discriminatory genomic regions) confers significant statistical and algorithmic advantages is a commonly occurring theme in computational biology [40].

The two biggest limitation of our model are (1) that one needs to define a set of gene seeds for the model to use when constructing its database and (2) that it is reference based. The seeds should be chosen appropriately given the context of the problem (Methods - Analysis Details). Moving beyond a reference database, it would be useful to adopt a methodology for *de novo* assembly of short genes for inclusion in the database, which does seem feasible given that we are not tracking an entire chromosome. In future work, we also plan to address the use of long reads (e.g. Oxford Nanopore, PacBio Hifi) as these technologies become commonplace in metagenomic studies. Even with state-of-the-art accuracy (e.g. 60% of reads being Q30 or better, as recently attained by [41]) this still leaves 40% of reads with 50 errors per 50000 kb^6^. Our model could also be used to overcome ambiguity for scenarios where the forward and reverse reads overlap, which occurs in 16Sv4 Amplicon sequencing. In addition, budget/experimental constraints typically enforce a limitation on sequencing depth, especially if multiple samples in time-series are to be analyzed. ChronoStrain would be an appropriate choice for overcoming ambiguity, especially in these use cases. We close by reemphasizing that no method is a panacea. With ChronoStrain, we suggest always inspecting the full time-series posterior distribution. Similar strains having highly overlapping credible intervals suggests that some strain clusters should be collapsed, as previously discussed with the fine-grained clustering results for UMB18 (Supp. Fig. S6). With that in mind, having the strain clustering thresholds learned jointly with the rest of the model could be an interesting and valuable addition to the model.

## Conclusion

Uncertainty quantification is an often overlooked concept in computational biology. Our method, ChronoStrain, incorporates uncertainty and time-awareness throughout its Bayesian probability model and exhibits significant performance improvements over current state-of-the-art. This allows for a more interpretable representation of how raw reads were likely to come from different genomes, and how that impacts our ability to estimate their abundances over time. We believe these results will have direct impacts on biological insights on microbial stability and competition, while the performance improvements will have nontrivial implications for the performance-to-cost ratio of time-series strain quantification experiments.

## Supporting information

Supplemental Data - Trajectories

## Code and Data Availability

The software and all analyses are available at https://github.com/gibsonlab/chronostrain. All UMB-related sequencing data, including the new MacConkey-culture sequencing experiments, are available under BioProject ID PRJNA400628. The ELMC sequencing reads were downloaded from the European Nucleotide Archive under accession PRJEB32631, and isolates under accession PRJEB22252. The ∼640k genome Themisto index was downloaded from Zenodo (7736981). Databases and raw outputs for all real-data analyses are available on Zenodo (10932690, 10932762). Databases and analyses can be reproduced using our codebase.

## Human Subjects

The UMB study [7] was conducted with the approval and under the supervision of the Institutional Review Board of Washington University School of Medicine in St. Louis, MO (IRB No. 201401068). Informed consent was obtained from all participants. Further details on the UMB cohort can be found in the Methods section of Reference [7] under the subheading Study design and sample collection - Enrollment.

## Contributions

YK Experimental Design. Statistical Model. Inference Algorithm. Software. Data analysis. Writing. Reviewing.

CW Experimental Design. Data analysis. Reviewing.

SA Inference Algorithm. Software.

LRvD Data analysis. Reviewing.

DA Software.

ZG Software.

PA Experimental design.

KD Experimental design. Data acquisition.

SH Experimental design. Data acquisition. Reviewing. Project management.

GKG Statistical model. Inference Algorithm. Data analysis. Reviewing.

AME Experimental design. Data acquisition. Data analysis. Reviewing. Project management.

BB Statistical model. Writing. Reviewing. Project management.

TEG Experimental Design. Statistical model. Inference Algorithm. Data analysis. Writing. Reviewing. Project management.

## Online Methods

### Overview of ChronoStrain

ChronoStrain is a Bayesian inference algorithm that learns the posterior distribution of bacterial strains’ relative abundances from metagenomic shotgun reads, by focusing on a subset of them which map to particular sections of the genome (e.g. marker sequences). Precise details of our model and its implementation is given in the next section; here we only give the high-level overview and explain key modeling decisions.

As input, one provides a database 𝒮 of strains and their marker sequences, a list of timepoints 𝒯 and each timepoint *t*’s collection of *N*_*t*_ corresponding reads. For each timepoint *t* ∈ 𝒯 and *i* ∈ [*N*_*t*_], each observed read *r*_*t,i*_ = (*ζ*_*t,i*_, *q*_*t,i*_) is specified as a (nucleotide sequence, phred quality score vector) pair.

The core machinery of the algorithm is based on a Bayesian model describing the joint distribution of (*X, Z, R*), where *X* = (*X*_*t*_)_*t*∈𝒯_ is a latent representation of the unobserved abundance profile at time *t, Z* is a vector that decides which strains are included in the population, and *R* = (*R*_*t*_)_*t*∈𝒯_ each is the sub-collection of reads of size *N*_*t*_ which align to our database. The latent representation (*X*_*t*_)_*t*∈𝒯_ is modeled using a standard Gaussian random walk (Wiener process [42]). The model inclusion vector *Z* = (*Z*_1_, …, *Z*_𝒮_) is a collection of i.i.d. Bernoulli(*π*) random variables. These determine the abundance trajectory using a sigmoidal function, resembling dynamic topic models [23]. Each read in *R*_*t*_ is modeled as being conditionally independent, given *X* = (*X*_*t*_)_*t*_. For the reads, we use the Phred model with random insertions and deletions, together with a model that considers a random choice of a position on the genome from the latent population that the read is measuring. We believe that the read model is especially important for strain-level resolution, since we care about single-nucleotide variants of genes and thus one *must* consider the inevitability of reads mapping to multiple genomes or loci with varying scores.

ChronoStrain performs variational inference to approximate the posterior distribution *p*(*X, Z* | *R*). We note that the algorithm implemented in this paper is specifically optimized and tailored for short Illumina reads; in the next section, we will point out exactly where this assumption is encoded. This leaves room for a model variant that accommodates long reads for future work.

The main difficulty for this task is in making the resulting algorithm scale efficiently with model size, which depends on the marker lengths, strains, and reads. Indeed, if implemented naively, practical computers would not have the required memory to store the full probabilistic model (easily taking up to hundreds of gigabytes for the database that we used). Our solution involves a heuristic sparsification of the data likelihood function (mathematically derived in Supplement §B.1). We show, using provided benchmarks on semi-synthetic data (Figure 2), that our heuristic works well in practice.

### Bayesian Model

First, we model the unobserved abundances using latent representations 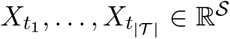 and a single vector *Z* ∈ {0, 1}^𝒮^. The *X*_*t*_’s are discretized observations of a Weiner process:

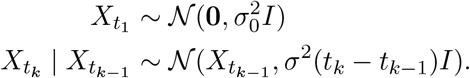

The scalar variance parameters 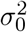, *σ*^2^ are each assigned independent instances of Jeffrey’s (improper) prior [43] for the variance of a Gaussian with known mean:

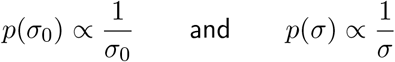

This prior was chosen to fulfill the role of a “non-informative” prior, and for its transformational invariance property. For users of our method, this means that the choice of measurement units of time – whether it is minutes, hours or days – does not matter. The elements of vector *Z* = (*Z*_*s*_)_*s*∈𝒮_ are independent Bernoulli(*π*). To transform the real-valued vectors *X*_*t*_ into relative abundances, we take 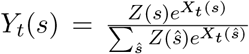, so that *Y*_*t*_ ∈ Δ ^𝒮−1^, the *S*-component probability simplex. This “exponential renormalization” mechanism (often called “softmax”) is used in machine learning, and was previously used to capture time-series relationships in continuous-time dynamic topic models [23, 44].

Next, we model the reads in two steps: the random position of the reads’ source fragments and then the sequencing noise. We model reads *r*_*t,i*_ = (*ζ*_*t,i*_, *q*_*t,i*_) without a mate pair as being a noisy measurement of a single randomly chosen *fragment f*_*t,i*_. The model for reads *with* mate pairs is cumbersome to describe, so for this section we assume unpaired reads; full details for the paired-read model are in Supplemental Text §B.2. In our model, a “fragment” is a substring of a marker sequence, representing the nucleotides which later get measured into reads. The primary assumption that we will make here is that each read’s source fragment overlaps with *some* marker in the database, thus necessitating a filtering step as described in Methods - Read Filtering. Conditioned on the *Y*_*t*_’s, we model the (timepoint *t*, index *i*) read *r*_*t,i*_ independent from all other reads as described below.

First, we introduce a few definitions. Allowing each marker sequence of strain genome *s* to be padded with *β* |*ζ* | of “empty” nucleotides on both ends, let 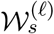 be the collection of all length-*ℓ*, position-marked sliding windows of markers of *s* for positive integers *ℓ*. We say that 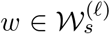 <u>induces</u> *f* if *f* is the string obtained from *w* by removing all padded bases; in particular, *f* is always at most as long as *w* (|*f* | *<* |*w*|). For instance, *β* = 0.5 guarantees that we only consider fragments *f* induced by ≥ 50% of the (short) read. Let 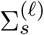 denote the set of all *f* induced by each 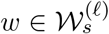. Finally, let 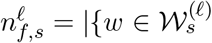: *w* induces *f*}| be the number of times *f* is induced, and let 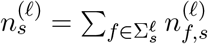 be its sum across all *f*.

Using the above definitions, we describe the fragment model. For each read *r*_*t,i*_, let *ℓ*_*t,i*_ be NegBinomial(*R*_NB_ *>* 0, *P*_NB_ ∈ [0, 1])-distributed. We model *f*_*t,i*_ as being sampled proportional to the frequency at which it is represented in the population at time *t*. More precisely (dropping the subscripts *t, i* to make it easier to read):

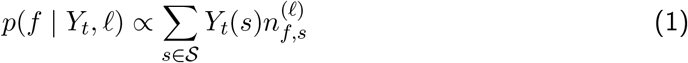

Note that this proportionality represents a normalization across all fragments *f*, and the normalization denominator 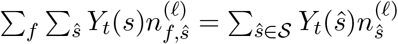 is a function of *Y*_*t*_ and *ℓ*. Algorithmically, a certain approximation of this (Supplement §B.1.2) results in an efficient correction for strains whose markers are over-represented in the database. As an aside, we remark that this is precisely the part of our method specially tailored for short reads. When operating on long reads (typically ∼10-25 kb or longer), our approximation fails to hold and thus our algorithm must be adapted to a different strategy. Furthermore, long reads can span *multiple* markers, thus requiring an adjustment in the fragment count definition.

Each read *r*_*t,i*_ = (*ζ*_*t,i*_, *q*_*t,i*_) ∈ *R*_*t*_ is modeled in the following way, treating *q*_*t,i*_ as a fixed observation. Conditioned on *f*_*t,i*_, we model the per-base sequencing error given by indels and substitutions, which we represent using alignments. For any alignment 𝒜_*t,i*_, meaning an arbitrary alignment of *ζ*_*t,i*_ to *f*_*t,i*_ represented by a 2 × *K* array of nucleotides (for some *K*) and gap symbols ‘-’, we model

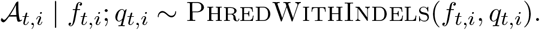

We temporarily drop the subscripts for exposition. The PhredWithIndels(*f, q*) distribution is supported over *feasible* alignments in the theoretical search space of the Needleman-Wunsch dynamic programming algorithm [45]. In particular, 𝒜 must satisfy max(|*s* |, | *f* |) ≤ *K* ≤ |*s*| + |*f*| and (# matches)+(# mismatches)+(# deletions) = |*f*|. Its likelihood function is given by a formula assuming fixed indel error rates *ε*_ins_ and *ε*_del_ and the standard phred score model:

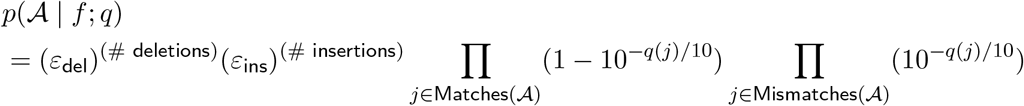

for any feasible alignment 𝒜. The parameters *ε*_ins_, *ε*_del_ are specific to the sequencing machine and may depend on whether the reads are forward or reverse in the mate pair.

Finally, conditional on 𝒜_*t,i*_, each mismatched/inserted base in read *r*_*t,i*_ is sampled uniformly at random; the likelihood of the read’s nucleotides *ζ*_*t,i*_ is the product

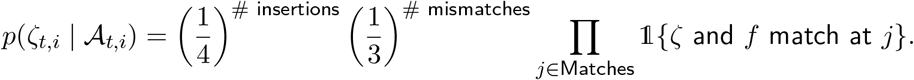

In its entirety, this model has hyperparameters *π, R*_NB_, *P*_NB_, *β, ε*_ins_, *ε*_del_. Our choices are explained in the supplemental text §B.3.

## ChronoStrain’s Database

In ChronoStrain’s model, a strain *s* ∈ 𝒮 is specified by a (multi-)set of markers ℳ_*s*_, where a “marker” *m* ∈ *ℳ*_*s*_ is simply a nucleotide sequence specific – but not necessarily unique – to that strain. Such a sequence could be, for example, a variant of a known gene encoding a particular target function of interest. Qualitatively speaking, an ideal marker is a gene, or any substring inside the genome, that has within-species variation (is *not* conserved species-wide) <u>and</u> presence/copy number variation (including non-”single-copy core” genes). Simultaneously, marker sequences from the target species should exhibit variation when compared against members outside the target species (e.g. homology with mutations). However, high sequence similarity outside the target species is inevitable — especially if we deviate from the usage of “core” genes — and thus our database construction includes steps to mitigate these confounders.

We pay no attention to markers’ ordering on the chromosome, but we do care about their multiplicity and their exact nucleotide sequence. In a later section (Methods - Analysis Details), we give a detailed explanation of the marker seeds used in our analyses. We also provide example statistics (e.g. number of reference genomes used and number of resulting clusters) and explain some of the idiosyncracies of the markers and the methodologies that we chose.

A key assumption required of our database (implicitly encoded in Equation 1) is that each ℳ_*s*_ includes all substrings of *s*’s chromosomes that share sufficiently high % identity. Stated another way: if a fragment *f* is in the genome of *s*, all occurrences of it in ℳ_*s*_ must be accounted for in the model, even if these mapped regions do not necessarily correspond to the same functional gene. Therefore, satisfying this constraint requires some careful consideration, especially when seeds represent genes with multiple known copies and/or homology outside of the species.

To construct such a collection, we require a FASTA file containing a reference sequence (or a multi-fasta file containing many known variants or homologs) for each seed. We allow for the seeds to represent genes which have homologs outside of the species of interest, and also appear multiple times within the same genome. For instance, all of our markers are genes specified by annotations or PCR primers. This comes with a myriad of complications: genes may have homologs within other species, vary in copy number, are possibly mis-annotated, or the PCR primers themselves may have mutations (known to be true for our *E. coli* genes [46]).

To account for these issues, our database construction begins by downloading all available assemblies from the same family as our target species (e.g. *Enterobacteriaceae* when analyzing *E. coli*). For each seed, we run a local BLAST query; the hits are thresholded by percent alignment identity (by default, 75%) and minimum length 150 (a typical read length), with `--max_target_seqs` = 10 × #(database genomes) to report a generous number of hits per marker seed query. The “best” percent identity threshold is expected to vary by gene and by target species. However, this is merely meant as a first-pass protection against out-of-species confounders; the main mechanism is our read filtering step (Methods - Read Filtering) for restricting reads which occurs after database construction.

Next, since the BLAST hits found above may overlap and/or may be contiguous, we process these hits by merging all pairs of hits which overlap on the corresponding reference genomes. For instance, if positions (35000-42000) and (41000-50000) are BLAST hits for genome *g* on the same contig/chromosome, then we merge them into a single hit spanning positions (35000-50000) forming a single marker on *g*. This is a crucial step: during our inference algorithm, reads may map to overlapping regions. These theoretically should only count *once* in the likelihood function, but if we were to use the raw BLAST hits they have the potential to be counted multiple times.

Lastly, to reduce the redundancy of the collection of strain genomes, we cluster them. Note that it is unclear how the markers constructed in the above steps are evolutionarily related (e.g. which markers are homologous to which other markers), nor is there a clear solution for forming a phylogeny when multiple copies of genes are present. Thus, it is important that the clustering method *does not* rely on multiple sequence alignment. To evaluate a distance metric *d* between each pair of reference sequences in an alignment-free manner, we used the tool `dashing2` [47] on the multi-fasta file of markers for each genome, which computes approximations of multiplicity-aware *k*-mer (multi)-set distances. This tool is the most compatible with ChronoStrain compared to other sketching methods such as Mash [48], because it incorporates multiplicities of fragments that occur across the markers. Using the pairwise distance matrix output by this tool, we ran scikit-learn’s implementation of agglomerative clustering with “complete” linkage; this is parametrized by distance threshold. Each cluster 𝒞’s representative strain *s*_rep_(𝒞) was chosen as the strain whose distances most closely resembles the whole cluster:

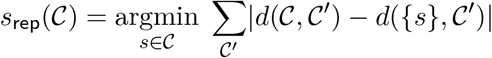

where 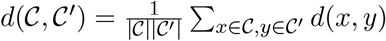 is the average distance between two clusters. Overall, assuming each strains contains *O*(1) sequences that approximately match each marker seed, the database size grows linearly in the product (# Clusters) *×* (# Marker Seed Length).

### Read Filtering

The read collection *R* of reads that map to database marker sequences is determined via alignment. In this step, we only require a one-sided test, in which we rule out all reads which definitely do not map to some marker of a known strain. Out of speed considerations, we use `bwa-mem2` to obtain these hits. Note that this step merely serves as a first-pass filter; exact analysis of these alignments and multiple mapping positions are left for a downstream step (using `bowtie2`, Supplement §B.1.1). In principle, any alignment program which outputs a standardized SAM/BAM file can be substituted in either step.

The alignment parameters were chosen specifically to suit our needs. More precisely, the match/mismatch penalties were assigned the log_2_-odds ratio of errors from the `Phred-WithIndels` model (assuming a rather pessimistic quality score of 20) in relation to a uniformly random sequence of nucleotides:

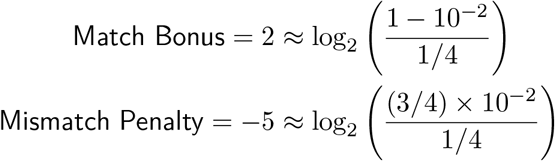

Assuming that indels are randomly distributed across each read according to indel error rates *ε*_ins_, *ε*_del_ (Supplement §B.3), we set the gap open penalty to zero and the extend penalties to − log_2_(*ε*_ins_), − log_2_(*ε*_del_).

Based on these alignments, we only kept reads that aligned to some marker sequence, where the alignment mapped with at least 97.5% identity. The % identity is calculated using a fitting alignment of the entire read, even if the alignment program produced local alignments. Specifically, in the case of `bwa` which only produces local alignments, we complete the alignment by re-attaching clipped bases.

### Target Posterior Approximation

Based on the Bayesian generative model, we aim to estimate the posterior *p*(*X, Z* |*R*). We employ ADVI [24], which uses stochastic optimization on Monte-Carlo estimates of the evidence lower bound (ELBO) objective. Using standard VI notation *q* to denote a generic approximate distribution: we used the factorized family *q*(*X*)*q*(*Z*) of densities for our variational fit. For the log-space variables *X*, we provide a fully-joint (|𝒯||𝒮|)-dimensional Gaussian approximation (a “full” covariance matrix) *q*(*X*); this is because we specifically wanted the time-associated uncertainty to show up in the posterior samples. Such posteriors are typically avoided due to their computational cost. However, we use a heuristic for pruning parts of the model based on the data available (Supplement §B.4), and were still well within limitations of a personal computer (Methods - Comp. Resources). For the scenario where memory is a limiting constraint, one can choose to factorize *q*(*X*) = Π_*t*∈𝒯_ *q*(*X*_*t*_), which we used for longer time-series (e.g. UMB stool samples). For the model inclusion posterior, we implemented a mean-field factorization *q*(*Z*) = *q*(*Z*_1_) … *q*(*Z*_𝒮_), where each *q*(*Z*_*i*_) is an independent “Gumbel-Softmax” relaxation [49].

The objective function was implemented and optimized using the JAX library [50] and the Adam gradient descent algorithm. The default settings, equivalent to the settings that we used for this paper, are as follows. The posterior approximation of *X* is initialized to have mean zero (corresponding to a uniform abundance profile for all timepoints) and covariance equal to the identity matrix. The learning rate decay schedule *η*(*t*) was initialized to *η*(0) = 0.0005. This was set to decay by a factor of 0.1, down to a minimum of 10^−7^, when the ELBO value (a Monte-Carlo estimate using *n* = 100 samples) no longer improved between epochs (1 epoch = 100 iterations) after at least five epochs. The Gumbel-Softmax is parametrized by a temperature parameter *τ*, initialized to 10.0 and is annealed by a factor of 0.95 every epoch, down to a minimum of 10^−4^. The optimization was set to stop after 1000 epochs, or sooner if the ELBO stopped improving by a multiplicative factor of 10^−7^. These settings were decided empirically, based on whether or not the gradient optimizer reliably improved the ELBO across real and semisynthetic data. We calculated all statistics using *n* = 5000 samples from this estimated posterior.

### Detection Classifier

For all real data analyses, we applied the following post-hoc method to interpret the approximated po<u>st</u>erior *p*(*X, Z* |*R*). First, we computed the collection of strain clusters 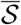 where each 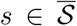 satisfies 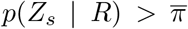. We chose 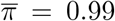, equivalent to a Bayes Factor threshold of 10^5^ when the prior is *π* = 10^−3^. Then, we sampled from the conditional posterior 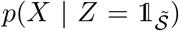, meaning that we conditioned on only 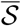 appearing in the model; the partial factorization of the variational solution makes this sampling trivial. For each timepoint *t*_*k*_ in the time-series, a strain *s* is marked as “detected” (as in Figures 3b and 5) if the resulting database-normalized relative abundance estimate `median`(*Y*_*t*_(*s*)) exceeds a cutoff *ρ*. Unless stated otherwise, *ρ* was chosen to be 5%; the “optimal” cutoff is database- and dataset-specific and we found that this simple value often yields a reasonable tradeoff.

### Analysis Details

Here, we describe the analysis details for each dataset (UMB, ELMC and semisynthetic), including marker seeds for the two databases used in this work^7^ and the settings that were used for each method. The precise database construction workflow for ChronoStrain is implemented as a Jupyter notebook for each dataset, available to view in our codebase. Analysis on real data for all methods (including the background reads from UMB18 for semisynthetic) were run on trimmed and de-contaminated data – see Methods - Sequencing & Real Data Processing.

### UMB Analyses (*E. coli* quantification)

For *E. coli* strain abundance estimation found in this work, our database seeds were:

1. genes from all *E. coli* MLST schemes on PubMLST [51],
2. genes used by the ClermonTyping tool [52] for phylotyping,
3. the O-antigen gene cluster, flanked by the JumpSTART and `gnd` primers [53],
4. H-antigen encoding (flagellar) genes annotated with names `fliC`, `flk*`, `fll*`, `flm*`,
5. annotated fimbrial genes `fim*`, and
6. annotated Shigatoxin genes `stx*`.

We note that there are currently two ST schemes for *E. coli* on PubMLST; we simply included all of the genes from both. Our catalog of reference genomes consisted of 5,405 whole-chromosome assemblies from the *Enterobacteriaceae* family, of which 2,063 were *E. coli*. After the BLAST and redundancy & overlap correction, our markers make up ∼1.5% of the genome when averaged across *E. coli* entries. For both UMB and semisynthetic analyses, we chose a 99.8% weighted marker *k*-mer frequency similarity (`dashing2` with ProbMinHash sketching) as a cutoff for the agglomerative clustering heuristic. After this process, we ended up with a database of 2,325 *Enterobacteriaceae* strain cluster representatives and their marker sequences; 842 are *E. coli*.

The systematic BLAST search & overlap correction steps were critical. For 2, 4, 5 and 6, we relied on genbank annotations; we found that several genes suffered from mis-annotations and/or unconventional naming schemes (e.g. `stx1A` versus `stxA1`), and thus the overlap correction helped correct for redundancies. Furthermore, the primers for the O-antigen gene cluster are known to have mutations in different sub-clades of *E. coli* [46], so the systematic BLAST search helps identify genes missed by the in-silico primer matching step (we used EMBOSS `primersearch` [54] which helps, but does not guarantee, finding all hits).

We ran ChronoStrain using this database with default inference settings, with prior *π* = 0.001 Results were interpreted using posterior threshold 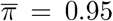 for semisynthetic and 0.99 for UMB; the latter was more stringent to aid visualization for a subset of the cohort. We ran StrainGST with default settings (*k*-mer length 23, *n* = 5 iterations and score threshold of 0.02) and using a database of Escherichia & Shigella genomes. We did not run the next tool in the StrainGE pipeline (StrainGR) which characterizes novel SNVs from the reads, since our goal was only to compare abundances. Note that while StrainGR does take multiple inputs simultaneously (across timepoints), it is not a tool designed for abundance estimation (although existing versions of the software do report estimates based on raw alignment/mapping counts using e.g. `bwa-mem`).

### ELMC Analysis (*E. faecalis* quantification)

For *E. faecalis* strain abundance estimation, our database seeds were:

1. genes from all *E. faecalis* MLST schemes on PubMLST [51],
2. PCR primer-specified pathogenicity/virulence-marking/polymorphic genes from [55],
3. a subset of 39 genes from the infants’ *E. faecalis* isolates.

To determine the genes to be used for #3, we first performed database construction and clustering at 1 − 10^−8^ similarity using just #1 and #2 (resulting in ∼450 *E. faecalis* clusters out of ∼660 total). Using these genes, many of the isolates across *different* infants are co-clustered even at this extreme of a threshold. It is possible, but less likely, that distinct infants have identical strains. Therefore, we operated on the assumption that most pairs across infants were indeed genetically distinct on some level, but #1 and #2 alone were not sufficient to distinguish them. In order to locate genes which finely separates the isolates, we annotated all of the isolate assemblies using NCBI’s `pgap` tool [56], and then for each gene name *g* we computed a multiple alignment using `MAFFT` [57]. For each gene *g*, cluster 𝒞 and for each pair of infants *i, j* whose isolates were in 𝒞, we computed a certain metric 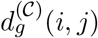 which quantifies how well *g* splits them. To explain it, let 𝒞 [*i*] denote the set of isolates from infant *i* contained within 𝒞, and for any isolate *x*, let *g*[*x*] denote the aligned sequence of the gene *g* in *x* (if *x* has multiple copies of *g*, then pick the last one in the GFF annotation; if *x* has zero copies of *g*, then take a sequence of gap characters). The formula for 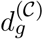 is

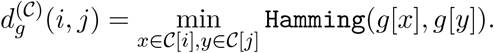

Finally, we picked the top *k* = 3 genes *g*_1_, …, *g*_3_ maximizing the number of nonzero entries of the distance matrix 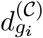 repeating this for each 𝒞 gave us 39 genes for marker group #3. The intent here is that genes *g* with many nonzero distances 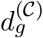 are those genes that will distinguish isolates from different infants. We made no attempt to “optimize” *k* in this work; the above scheme was a heuristic simply to get the analysis to run while having reasonable runtime and memory constraints.

Our reference genomes consisted of 1,087 complete chromosomal assemblies from the *Enterococcaceae* family (excluding RefSeq *E. faecalis*) to account for sequence similarity-induced confounders, plus the 2,026 + 350 *E. faecalis* isolates from Europe and ELMC as in the original mGEMS analysis [28]. Averaged across *E. faecalis* entries, our markers make up ∼1.6% of the genome. We used a 99.8% marker similarity cutoff, which resulted in 533 *Enterococcaceae* strain cluster representatives, of which 387 are *E. faecalis*. Of these, 83 contain at least one ELMC isolate.

We ran ChronoStrain using the above database and default inference settings, with prior *π* = 0.001. Results were interpreted using posterior threshold 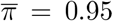. For the sake of comparison, we re-ran the mGEMS pipeline (themisto + mSWEEP + mGEMS bin extraction) in a hierarchical style as in [28]. This means that we first ran the pipeline to bin reads by species, then a sub-analysis on the *E. faecalis* bin, and finally `demix_check` to check the quality of the strain bins. Finally, we only kept strain bins with abundance greater than 0.01, and removed bins with poor confidence scores (3 or 4).

We re-ran the mGEMS analysis so as to have access to the intermediate outputs (the species-level estimates and read bins) when running Themisto v3.2.1, mSWEEP v2.0.0 and `demix_check`. We first reproduced the original results using the original database used in [28], but due to a version incompatibility, we used an updated pair of pre-compiled indices. For the species-binning step, we used the recently published index of ∼640,000 genomes [35] compatible with Themisto v3.2.1. For the strain-binning step, we used a faithful reproduction of the *E. faecalis* index of (a subset of) the original 2,026 European isolates plus 350 ELMC isolates. Both were provided by the authors of [28]; we verified that using the above reproduced the original results before proceeding.

We note that the above *E. faecalis* index contains only 1,229 of the 2,026 European assemblies (due to an undocumented filtering step in [28]). Furthermore, only 37 out of 168 clusters contains ELMC isolates, so the clustering is quite coarse. Thus, for our main results, we compiled a third index for *E. faecalis* quantification using all 2,026 European isolates. To create it, we re-ran PopPUNK using the `threshold` method with threshold 0.00036; this was tuned manually, so that PopPUNK produces exactly 83 clusters containing at least one infant isolate just like ChronoStrain. Interestingly, this produced 808 total clusters (much larger than ChronoStrain’s 533), confirming that our earlier gene-picking heuristic successfully produced markers that are particularly discriminative for ELMC isolates. Additional species-level quantification was done twice, once using Kraken2 v2.1.3 + Bracken v2.9 (database:`k2_standard_16gb_20240112`), and once using MetaPhlAn4 v4.0.6 (database:`202307`). We found that both third-party method outputs agreed for many samples on *E. faecalis* and the results are largely similar regardless of which of the two we chose. Finally, we re-ran both methods using a mutated database to test their robustness to the database on real data; refer to Supplement §B.6 for details.

### Semisynthetic benchmark

For the purpose of benchmarking, we generated a “semi-synthetic” dataset by merging synthetic reads with real ones. We created a database for each method using their own provided (or documented) algorithm for creating strain clusters. For ChronoStrain, we provided the same database as in the UMB analysis. To ensure that all clustering methods ended up with the same granularity for the benchmark, we tuned each method’s distance threshold so that they yielded roughly the same number of phylogroup A clusters (341 *±*2) as ChronoStrain. Specifically, for mSWEEP we ran PopPUNK [36] with distance thresholding (`--fit-model threshold`) to control the granularity of the *Enterobacteriaceae* assembly catalog. For StrainGST, we ran its built-in clustering on the *Escherichia* and *Shigella* genomes (the program is designed to warn the user if we simultaneously include other genomes that are not closely related). For StrainEst, we ran their database construction method, which requires picking two sets of representative genomes: (1) a core list of ten genomes, which are used build a list of SNV loci via whole-genome multiple alignment, and (2) a clustering of all genomes and their representatives, to characterize the nucleotides observed at these pre-calculated loci. For (1), we used the authors’ suggested method of performing hierarchical clustering by Mash distance and picking a cutoff that results in ten genomes. For (2), the recommended method is to again perform mash clustering; to reduce some variability between clusterings (which affects the dataset generation), we re-used the clusters generated by PopPUNK. Though the PopPUNK metric is more intricate than the Mash metric, both are *k*-mer-based; thus, we had little reason to suspect it would be particularly disadvantageous for StrainEst.

Overall, our dataset is made up of ten “genome” replicates (sampling six genomes and random mutations), times two “sequencing” replicates (Art is re-run with a different seed), across five possible simulated read depths, totalling 100 distinct overall replicates. We describe this process now. First, we sampled six phylogroup A *E. coli* genomes at random for each “genome” replicate, that are not a part of the original database. To do so, we first ensured that each cluster is fairly represented across the replicates by assigning to each genome a probability weight proportional to the reciprocal of the root-mean-square of the respective databases’ cluster sizes (so that each cluster is chosen with roughly equal weight regardless of the method used). We pick six random genomes one at a time without replacement using these weights, and for each we remove all other genomes that share a cluster with it (for all clustering schemes) at each step, so that no cluster is chosen twice. To each chosen real genome, we introduced random mutations by flipping a coin of bias *p*(Heads) = 0.002 for each base. If Heads, we chose one of the three remaining nucleotides at random to substitute.

For each “sequencing” replicate, we took these six synthetic genomes and simulate reads using Art [58] and its built-in HiSeq (length 150) error profiles. The reads were sampled according to the counts given by a Multinomial(*N*, 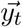) distribution, where *N* is the chosen simulation read count and 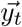 are the abundance ratios for the six synthetic genomes for timepoint *t*. The time-series ratios 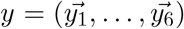 were chosen and fixed beforehand; these trajectories span a wide range from 0.1 to 10^−3^, where some strains fluctuate between different orders of magnitude. The simulated reads span six timepoints, and they were merged with the first six timepoints from UMB18’s stool sample sequences. This choice was made based on a preliminary analysis using the methods, which suggested that phylogroup A was either absent or undetectable in these samples. What makes this dataset extra challenging is that there are several trajectories that dip into an “ultra-low” species-level ratio (10^−3^). At the lowest simulated read count (*N* = 2500), this approximately amounts to ∼10^−7^ overall relative abundance, after accounting for the ∼10 million real reads.

Due to the difficulty & expectations of this benchmark, we were required to make some changes to the settings of all methods except ChronoStrain. First, StrainGST calls at most 5 clusters by default; to allow this method to infer strains beyond what it would have returned on the background samples alone, we raised this value to 20. For StrainEst, we noted that when run on default settings, the program failed to recall any of the simulated clusters when *N <* 20000, so we raised its sensitivity slightly (`-p 5 -a 3`) which incurs runtime cost. We ran mGEMS using the hierarchical approach described in [28]; we first used a large bacterial genome catalog to bin reads by species, then ran *E. coli* strain analysis on the *E. coli* binned reads. The species binning step used a large ∼640k genome index, which is also used in the ELMC analysis. When we ran `demix_check` on the *E. coli* bins, all scores were 4 (the worst possible score) for all bins, including the ground-truth bins. Note that this is expected behavior for sufficiently in-silico mutation rates: these scores are between 1 (best) to 4 (worst) and measure *novelty* of the binned reads compared to the database; the scores are biased in favor of bins which already contain the sample’s latent genomes. Thus, thus we did not factor these scores into the analysis.

For measuring the error in abundance estimation, we computed the RMSE-log after re-normalizing across a subset of clusters (the six source clusters, or the all clusters labeled as phylogroup A). For an estimate *ŷ*, the RMSE-log metric is given by the formula

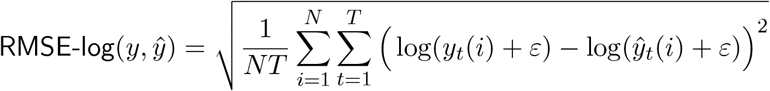

where *ε* = 10^−4^, an order of magnitude smaller than the smallest ratio in *y*.

For mGEMS, the raw mSWEEP estimate contains a considerable amount of noise, coming from the fact that many reads pseudo-align to a majority of database strains. We built an estimator *ŷ* using mGEMS outputs to help it minimize the RMSE-log error (which is sensitive to noise at or above *ε*), in the following manner. Starting with the raw mSWEEP estimate 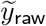, we build the phylogroup-only vector (or the target-only, length-six vector) 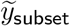 by restricting the vector to the desired entries and renormalizing. Then, we zeroed out all entries below *ε*_LOD_ and renormalized once more; this helps remove the effect of noise from the RMSE-log. Since this threshold is applied after an initial re-normalization on the target subset, we chose *ε*_LOD_ = 0.001 so that the lowest ground-truth ratio ∼ 0.0025 was allowed to appear. This parameter is necessary to control the RMSE-log error for mSWEEP (Figure 2b,c), because the error tends to be large when false positives are not accounted for (Supplemental Figure S3b).

To evaluate the classification metric (AUROC), we turned each method into a classifier by applying an abundance threshold to determine “detection”. For ChronoStrain specifically, we varied the abundance threshold *ρ* described previously (Methods - Detection Classifier), keeping the posterior threshold 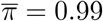 fixed. Note that the varying the abundance threshold for mSWEEP here means controlling the limit of detection parameter *ε*_LOD_.

### Computational Resources

For benchmarking, all four methods were run on stock Alienware Aurora R15s (Intel 12900KF with 128 GB of RAM). ChronoStrain’s inference step, in particular, was run on a single RTX 3090; other benchmarked methods were not designed with GPU hardware in mind. GPU memory size is the primary limiting factor that requires us to cluster the database. If one includes many more markers, more database clusters and/or more samples, a GPU with more memory is required. All benchmark analyses were able to fit on the RTX 3090 (typical CPU memory footprint was less than *∼*10GB CPU RAM).

### Sequencing & Real Data Processing

MacConkey-cultured samples were sequenced in the same manner outlined for the original UMB dataset [7]. Starting with the raw reads, we used the demultiplexed, whole-genome portion of the UMB dataset for all participants. Just prior to analysis, all reads from UMB were preprocessed using the KneadData pipeline (v0.11.0, https://huttenhower.sph.harvard.edu/kneaddata/), which invokes Trimmomatic v0.39 [59] to trim adapters and low-quality bases at the ends (phred ≤ 10) and invokes Bowtie2 [60] to discard reads which align to the human genome. The ELMC metagenomic reads available online did not require trimming and decontamination.

## Footnotes

1 Lack of phylogroup A was determined using StrainGST. Even if phylogroup A were actually present but rare, any potential gaps in performance between the methods are expected to close as we add more synthetic reads.

2 By “temporal inconsistency”, we are referring to the fact that for the samples between `2016-02-23` and `2016-09-13`, StrainGST shows an alternating presence-absence (mutually exclusive) pattern for two of the Clade D strains in participant UMB18’s time-series (Figure 3f,g). The pattern for ChronoStrain is the opposite: we do not see this alternating and mutually exclusive pattern for pairs of strain clusters over time.

3 By “stability”, we mean “asymptotically stable”. This means perturbations do not cause an arbitrary deviation in the system from its unperturbed state (equilibrium point) and upon removal of a perturbation, the system returns to its prior equilibrium point [31].

4 This is separate from the strain-level parameter *ε*_LOD_ = 0.01, which is only applied for strain calls after inference. The behavior we are demonstrating occurs for the species abundance estimate, which occurs before the strain-level analysis; the choice of *ε*_LOD_ appears to be a coincidence.

5 Strain “turnover” is defined as an event where the most abundant strain at one timepoint is no longer the most abundant strain at an adjacent timepoint, computed here only within the context of *E. faecalis*.

6 For comparison, *E. coli* markers add up to *∼*74k bases per genome in this work.

7 When building the reference genome catalog, out of caution, we preferred to use *complete* chromosomal assemblies for our reference database (e.g. downloaded from NCBI’s `Assembly` database). The primary reason was that retrieval of scaffolds or contig-level assemblies often also included MAGs. In general, MAGs are often mis-assembled or contain contaminations (e.g. from the human host). In theory, for bacterial quantification, using bacterial-specific markers and filtering metagenomic reads by alignment at 97% similarity should provide some protection; however, our seed construction step was “loose” enough at 75% identity threshold that we could not rule out these as potential confounders.

