## Supplemental Figures for "Strain tracking with uncertainty quantification"

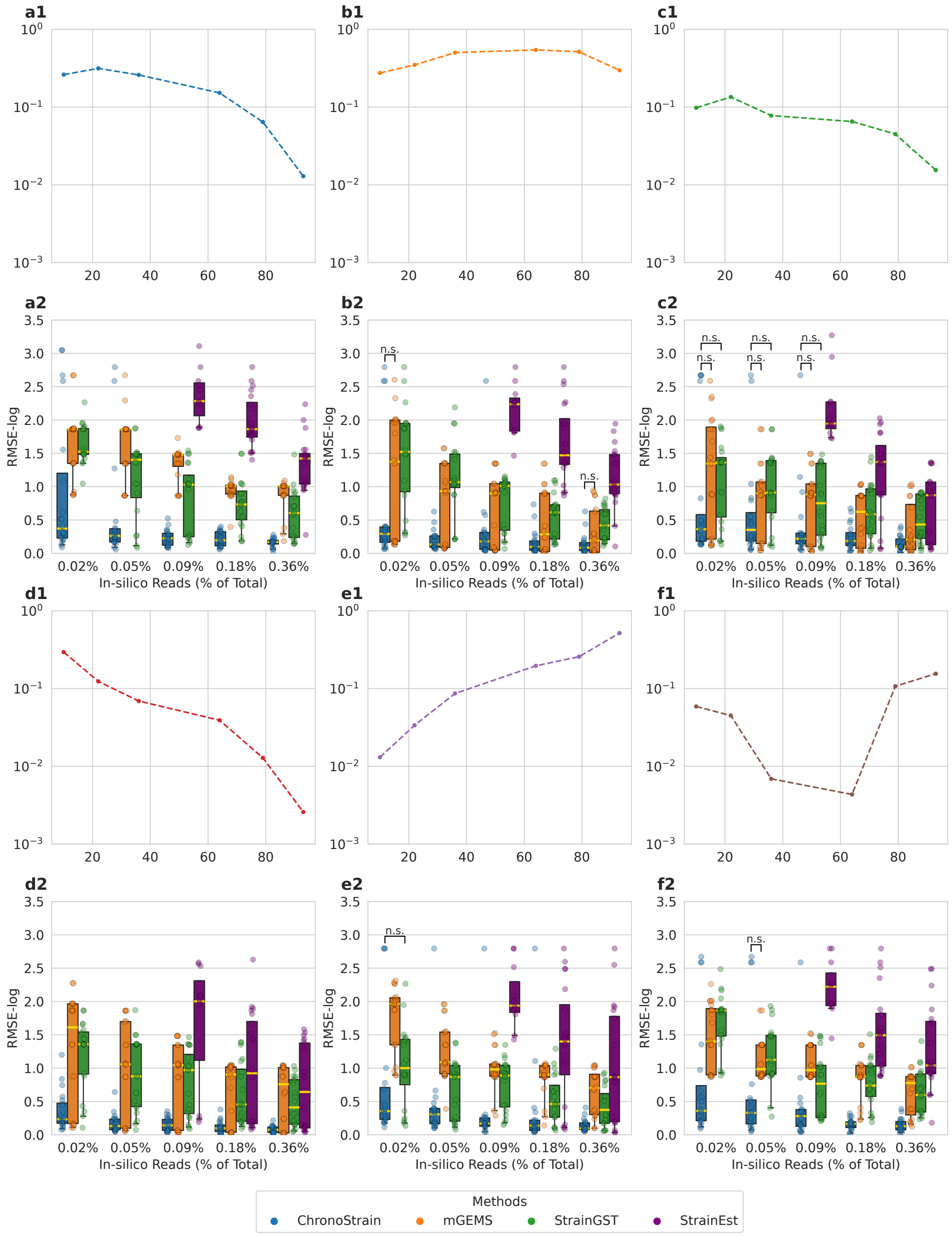


**Supplemental Figure S1: (Semisynthetic benchmark) Target strain-specific RMSE-log error performance. (a1-f1)** The ground-truth trajectories, individually plotted. **(a2-f2)** The corresponding RMSE-log error for each trajectory estimate, representing a per-strain breakdown of the RMSE-log metric in Figure 2a. The simulated genome is randomized across replicates as described in Methods, so each trajectory is not one fixed strain, but rather a family of randomized genomes and reads sampled at specific ratios across time. Significance was determined via two-sided Wilcoxon test $p$-values with significance level $0.05$ after BH correction; only non-significant pairs are marked via "n.s.".

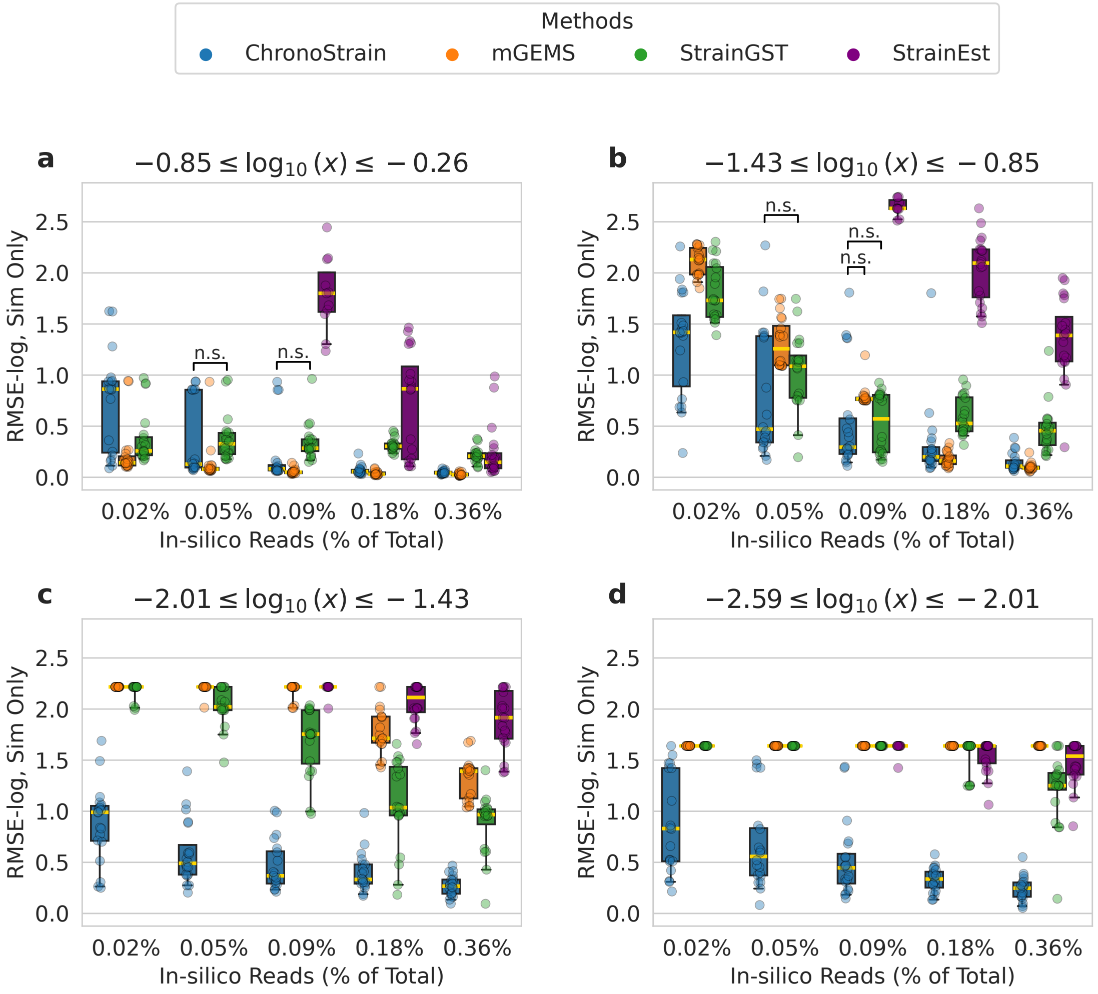


**Supplemental Figure S2: (Semisynthetic benchmark) Abundance-specific RMSE-log error performance.** To break down the contribution of the estimates to the RMSE error by abundances, we evaluated the RMSE-log error after binning (timepoint, synthetic strain) pairs by ground-truth abundance ratios. **(a-d)** Bins are ordered from "most abundant" strain in timepoint to "least abundant", where $\log_{10}(x)$ indicates the log-ratio. The RMSE-log was evaluated after adding $10^{-4}$ to predictions to handle zeroes. Significance was determined via two-sided Wilcoxon test $p$-values with significance level $0.05$ after BH correction; only non-significant pairs are marked via "n.s.".

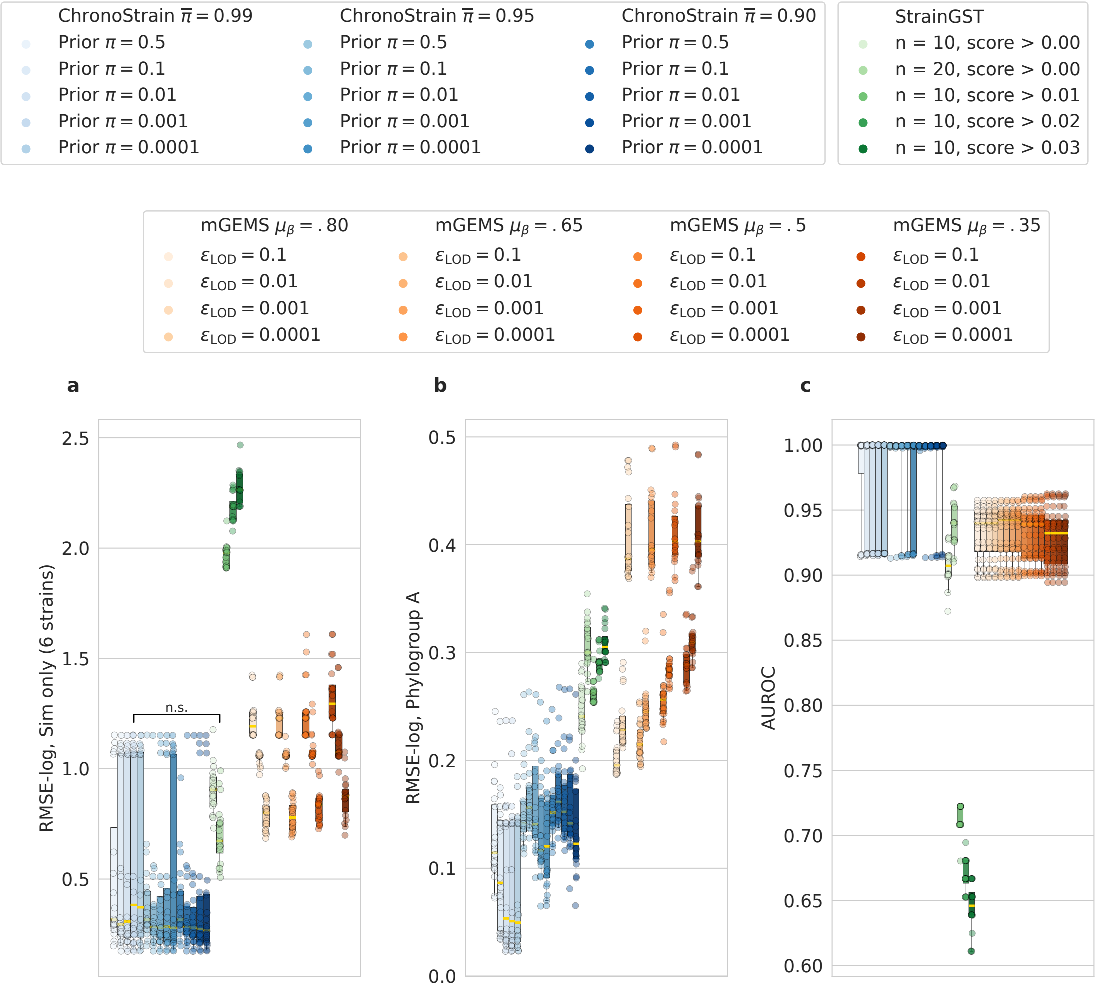


**Supplemental Figure S3: (Semisynthetic benchmark) Sensitivity of methods to method-specific parameters for 10,000 simulated reads.** For ChronoStrain, we varied the parameter $\pi$ of the model inclusion prior and the post-inference threshold $\overline{\pi}$. Like the main result (Main Figure 2), the AUROC for any given pair $(\pi, \overline{\pi})$ is evaluated by varying an abundance threshold. For StrainGST, we varied the number of iterations, which is an upper bound on the number of strains it will output. For mSWEEP, we varied the abundance threshold parameter $\varepsilon_{\mathrm{LOD}}$ which is applied after inference. The errors are defined the same as in Fig. 2: **(a)** The RMSE-log error on the six target clusters, **(b)** the RMSE-log error using all phylogroup A clusters, and **(c)** AUROC by varying the abundance threshold. ChronoStrain's median errors are largely insensitive to the parameters, though the variance shrinks as we lower the posterior threshold (thus removing the extra uncertainty induced by the posterior presence/absence variable $Z$ but increasing false positives for a fixed abundance threshold). The paired, two-sided Wilcoxon test was only performed between the parametrization used in the paper ($\pi = 0.001, \overline{\pi} = 0.99$) versus every StrainGST & mGEMS parametrization, with BH correction applied. In all plots, all BH-corrected $p$-values were below $0.05$.

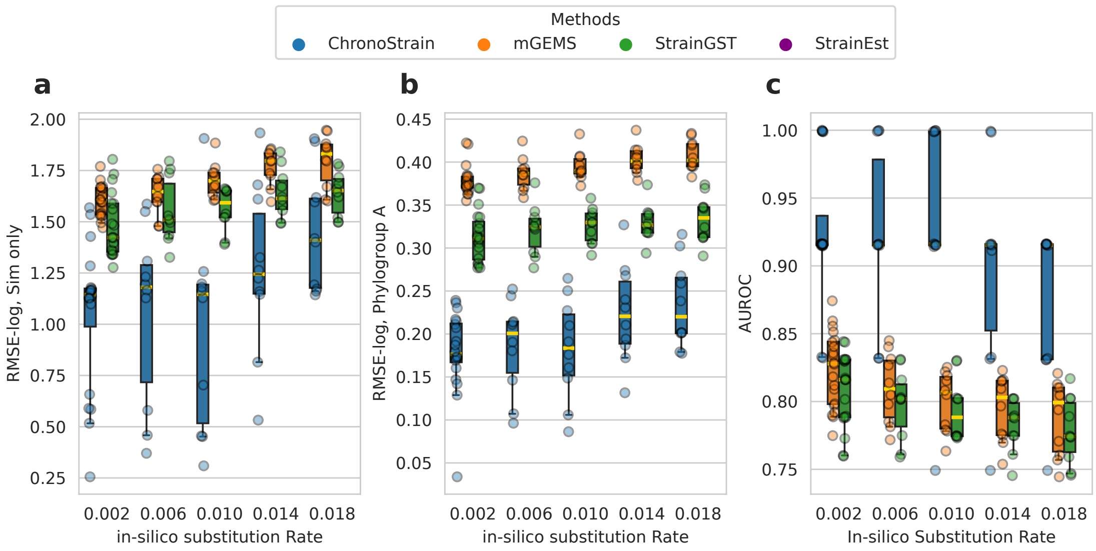


**Supplemental Figure S4: (Semisynthetic benchmark) Sensitivity of methods to an in-silico substitution rate on the target genomes.** We compared the performances of each method by varying the in-silico substitution rate of the semisynthetic dataset. To illustrate the scenario where the decay in error would be the most drastic, we used a low simulated read count $(2500)$ which is expected to be the hardest and largest-variance scenario in our benchmark. The errors are defined the same as in Fig. 2: **(a)** The RMSE-log error on the six target clusters, **(b)** the RMSE-log error using all phylogroup A clusters, and **(c)** AUROC by varying the abundance threshold.

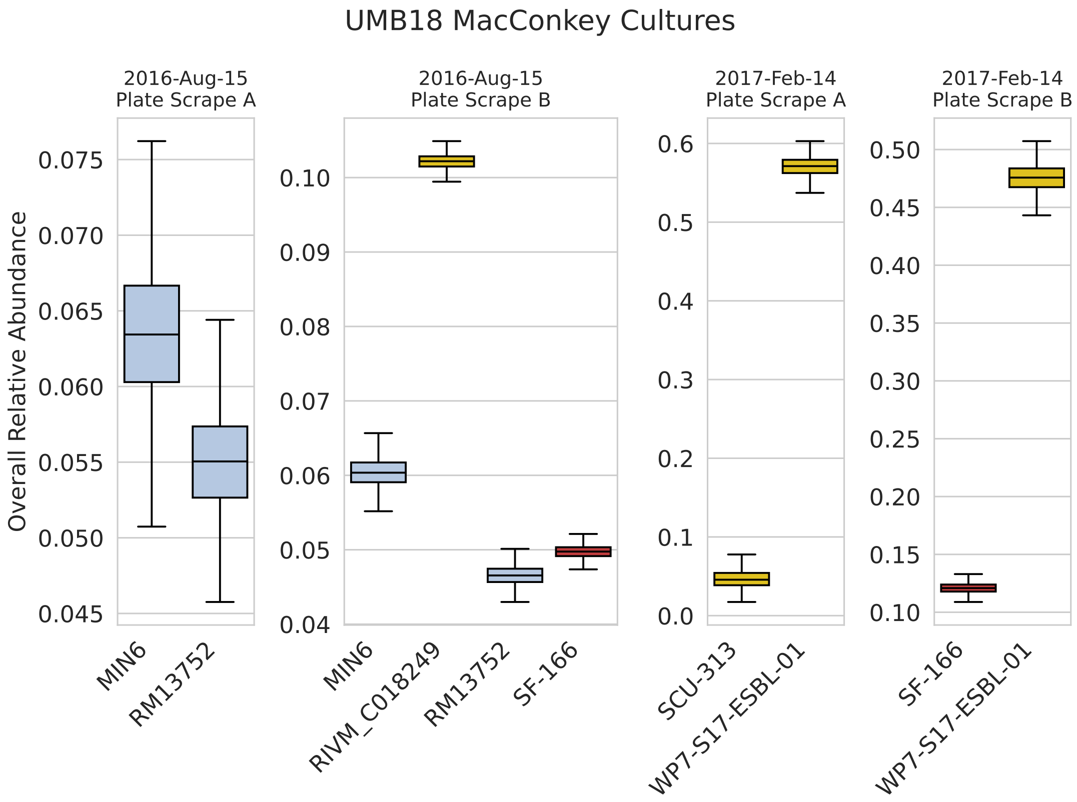


**Supplemental Figure S5: (UMB) ChronoStrain's raw estimates for the *E. coli* abundances of MacConkey-cultures grown from UMB18 stool samples.** The $y$-axis quantifies the strain clusters shown on Main Figure 3b, originally drawn using X's for the corresponding timepoint. Colors are chosen using the same phylogroup-based color palette.

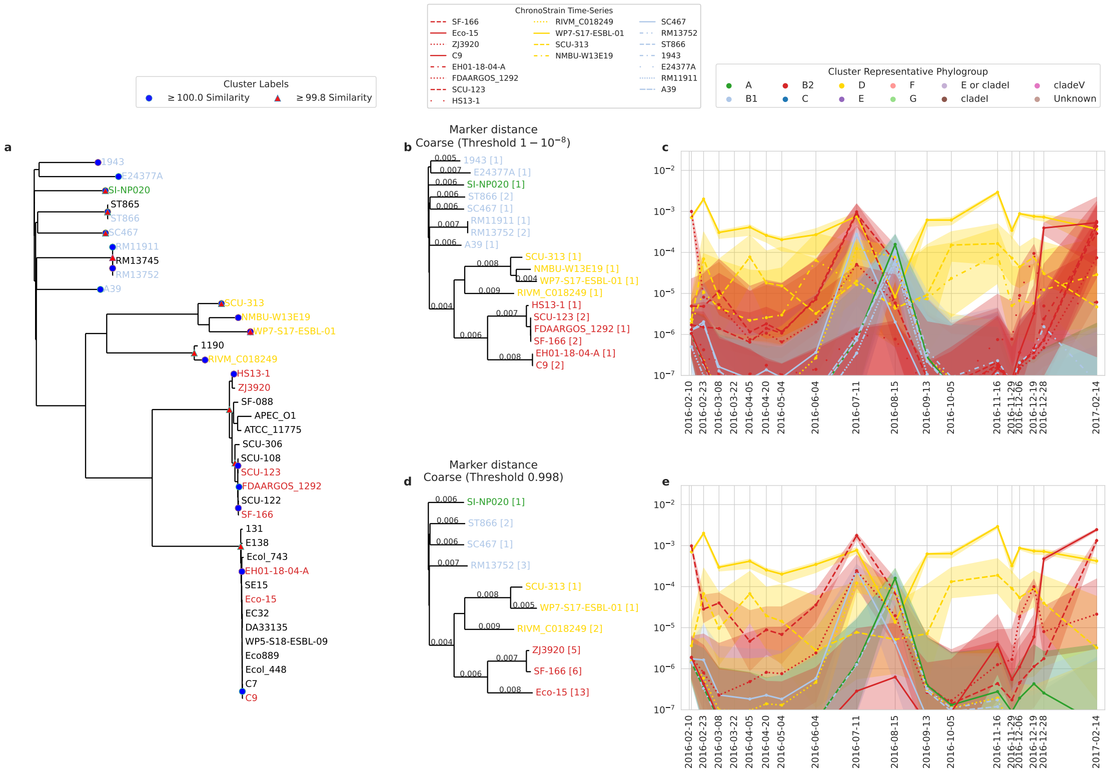

**Supplemental Figure S6: (UMB) A side-by-side comparison of ChronoStrain run with the 99.8% thresholded database and a run with near 100% threshold for UMB18.** **(a)** A phylogenetic tree (constructed from marker $k$-mer distances) of all cluster representatives and their constituents; only the representatives' leaves are labelled with names. The least common ancestors of the clusters' members are annotated with markers, distinguished by the threshold. **(b, d)** The threshold-specific subtree of cluster representatives called by ChronoStrain in the time-series; the size of each cluster is indicated in brackets. **(c, e)** The threshold-specific result corresponding to the trees; the fine-grained run is above the coarse-grained result. Both runs were filtered using posterior threshold $\overline{\pi} = 0.95$ and abundance threshold $= 5\%$. There are more clusters in the fine-grained result (top) due to the increased granularity, but the top plot has *increased specificity* in the sense that the called clusters' sizes sum up to fewer members than the bottom (23 versus 34). In particular, only 3 members out of the 99.8%-thresholded `Eco-15` cluster (13 members) are represented in the sample.

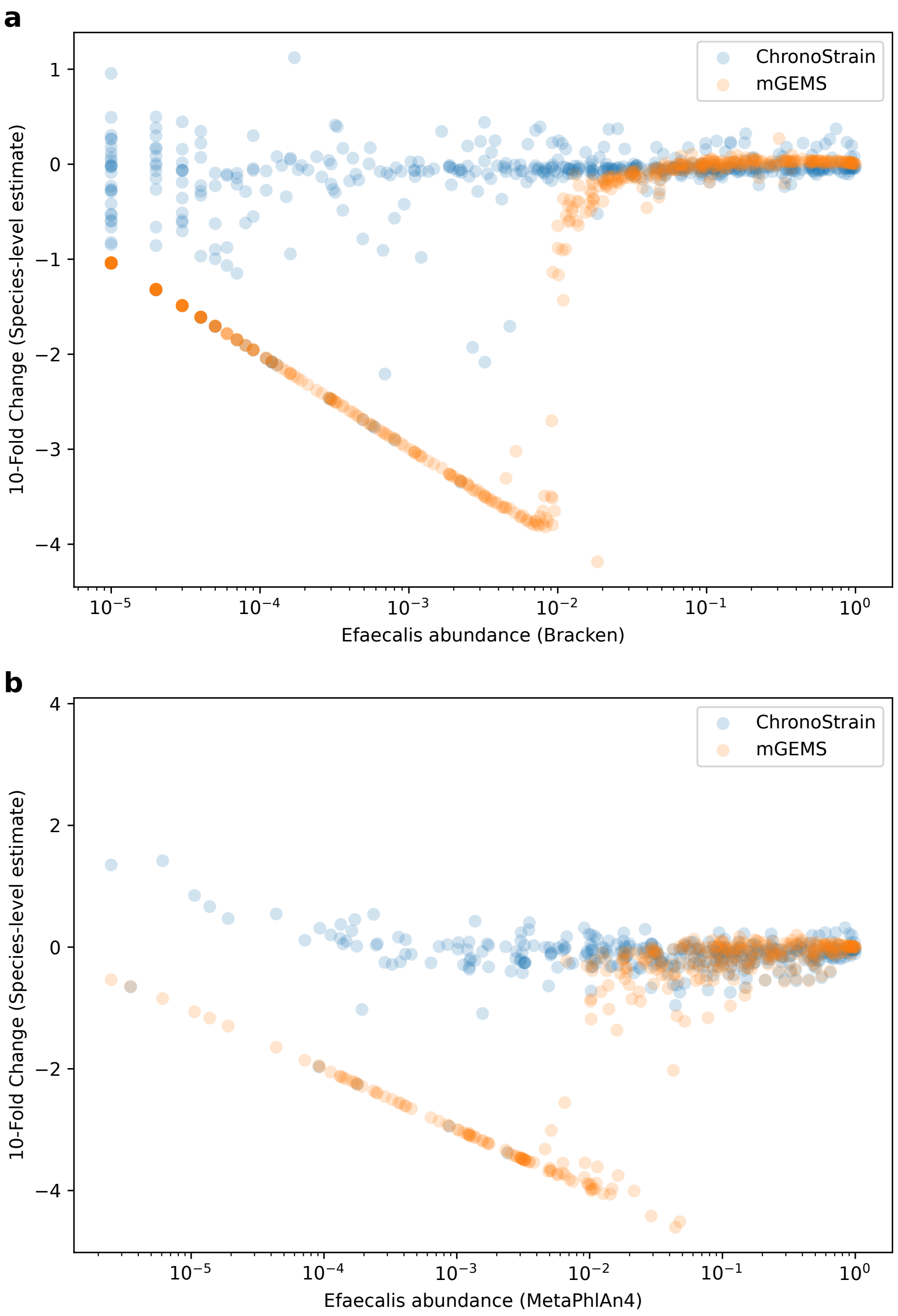


**Supplemental Figure S7: (ELMC) A comparison of the fold-change versus third-party methods for ChronoStrain and mGEMS' *E. faecalis* species estimates.** We ran two methods for species-level comparison: **(a)** Kraken2 + Bracken and **(b)** MetaPhlAn4. Each dot is a single infant sample. In both plots, the fold change is computed as the difference between $\log_{10}$ of the predictions plus $\varepsilon = 10^{-6}$ to avoid NaN's. The dots forming a straight line on the bottom of the scatterplots (the line $\log_{10}(x) + y = -6$) are an artifact of this $\varepsilon$, where either ChronoStrain (resp. mGEMS) or Bracken (resp. MetaPhlAn4) – but not both – estimates zero/near-zero abundance for *E. faecalis*. For mGEMS, a sharp drop-off just above $10^{-2}$ is visible in both **(a)** and **(b)**, suggesting that the method has a detection threshold at that location. In contrast, ChronoStrain generally agrees with both third-party methods all the way down to $10^{-5}$, with higher spread as *E. faecalis* becomes rarer in the sample.

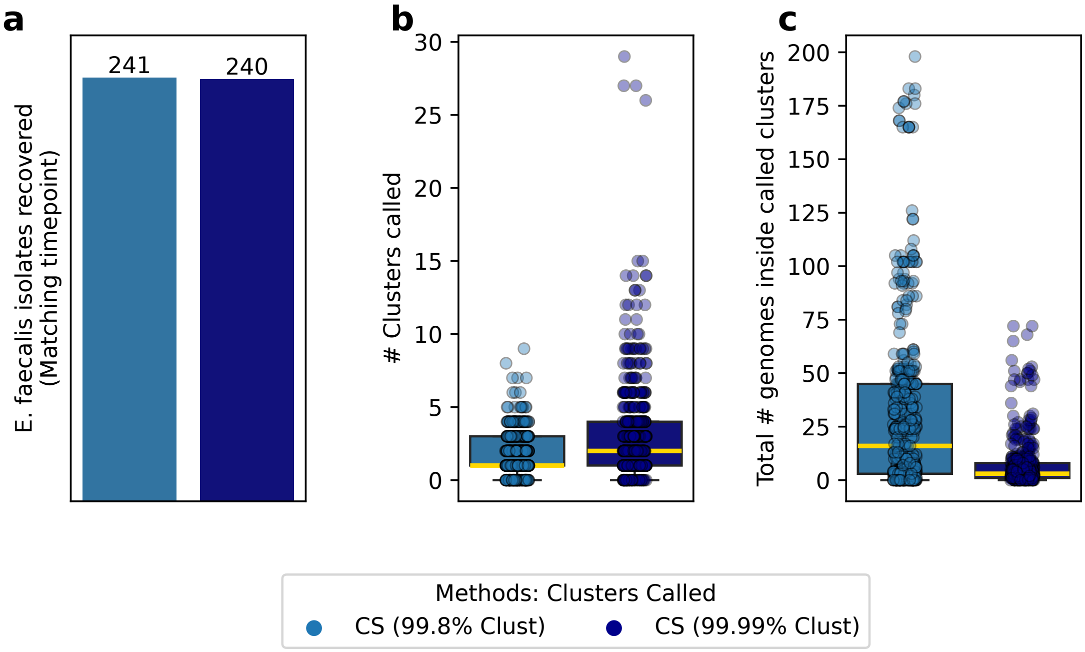


**Supplemental Figure S8: (ELMC) Counts of the number of called clusters, and the total size of these clusters, contrasted across ChronoStrain with two different databases.** One is the 99.8% identity thresholded run from the main result, the other uses a 99.99% similarity threshold on the clusters, which breaks many of the infant isolate clusters into smaller parts. **(a)** At the same posterior threshold $\overline{\pi} = 0.95$, the strain abundance threshold was re-calibrated to $0.025$ (required to account for change in database size) so that the number of within-timepoint isolate calls are approximately equal. **(b)** The fine-grained run calls a median of 2 clusters which is one more than the median of 1 for the coarse-grained run. **(c)** However, the fine-grained run produces a median of 3 genomes in total when adding up the cluster sizes. Compared to 16 from the original run, this suggests overall improved specificity.

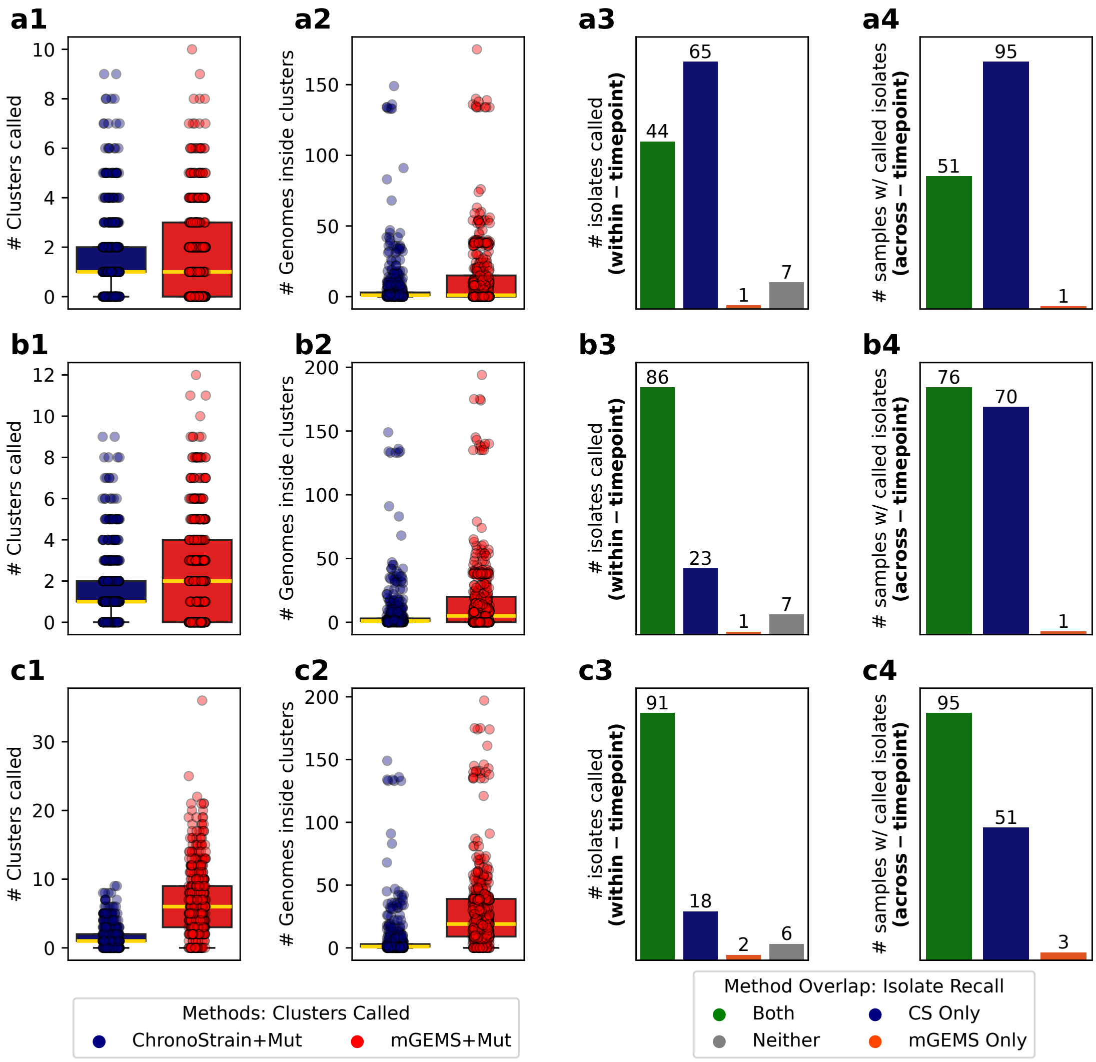


**Supplemental Figure S9: (ELMC) Inference result comparison using the mutated database of 117 infant isolates (Supplement B.6), in the style of Figure 5(c,d,e,f).** The first row (**a**) only retains mGEMS predictions with `demix_check` quality scores 2 or better. The second row (**b**) retains 3 or better, third row (**c**) is 4 or better. ChronoStrain thresholds are held fixed in all three rows (ChronoStrain: $\overline{\pi} = 0.95$, ratio $\geq 0.065$). One may loosen the `demix_check` threshold in order to obtain comparable numbers of isolate calls (**a3**,**b3**,**c3**) at the cost of calling more clusters (**a1**,**b1**,**c1**), whereas ChronoStrain remained largely the same from the unmodified run from Figure 5 (Fig 6).

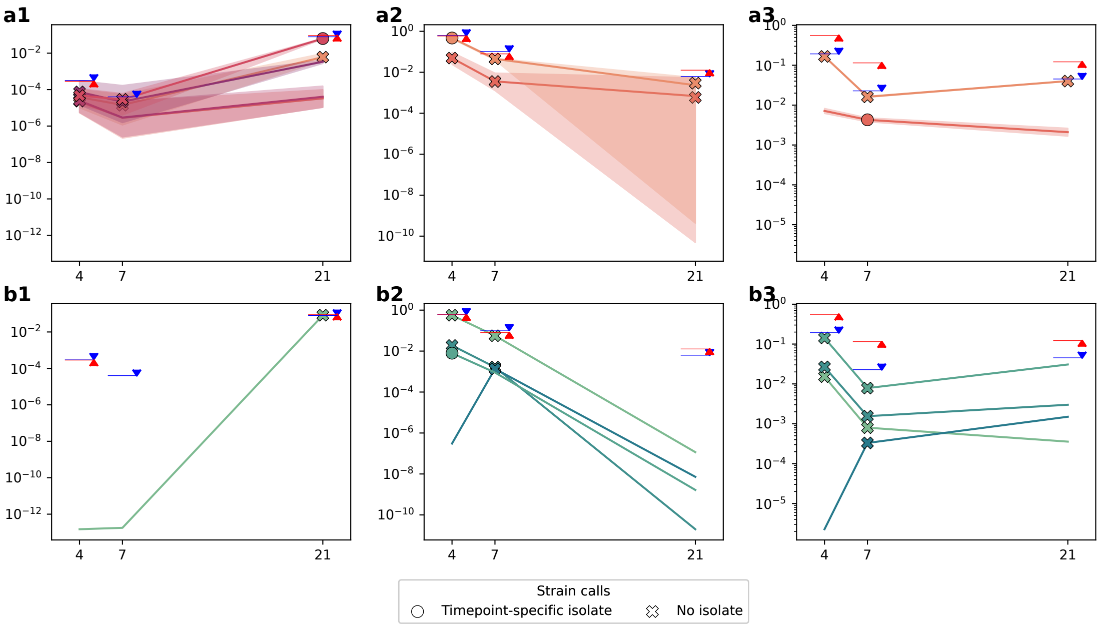


**Supplemental Figure S10: (ELMC) Abundance trajectories of clusters using mutated databases.** Using the mutated analysis results, we plotted the inferred trajectories for the same three examples shown in Fig. 5a,b, drawn in the same style. Just like before, the trajectory for a cluster is rendered only if it passes the filter for the respective method in at least one timepoint. For each trajectory, for each timepoint at which the cluster passes the filter, we mark it with an O (contains an isolate cultured at that timepoint) or X (no isolate for that timepoint). After mutation, mGEMS' filter no longer calls the corresponding isolates in infants `A01077`, `B02273` (**b1, b3**) whereas ChronoStrain remains unchanged from the original inference (**a1, a3**). For `B00053` (**a2, b2**), the isolate is called correctly at the first timepoint for both methods but in mGEMS is no longer the dominant strain, whereas both methods in the original analysis agreed that it was the dominant strain in the sample (Figure 5, panels a2, b2).
