## Supplemental Text for "Strain tracking with uncertainty quantification"

#### A Related Works

According to the overview of strain detection methods in [1], ChronoStrain can be categorized as a *read alignment-based* method. Here, we focus on the *abundance estimation* aspect of strain inference (which is different from *detection*), and thus organize the methods slightly differently. More specifically, we emphasize three orthogonal features of existing methods: (1) *whole-genome based* versus *gene based*, (2) *read modeling* versus *allele frequency modeling*, and (3) *genotype-learning* versus *reference calling*.

In this work, we largely omit from the discussion methods which rely on metagenomic assembly. This is because assembling *low-abundance* strains (such as UPEC in the gut) requires an extremely high read depth to be successful. Furthermore, existing methods [2] that nevertheless attempt this strategy produce contigs that are binned into metagenome-assembled genomes (MAGs). However, when genomes exhibit sufficient *overall* similarity — such as with phylogroup B2/F/G *E. coli* strains — it can be challenging to resolve contigs into MAGs correctly. Still, the general idea of using assembly (or a related algorithm) using reads still has some merit; we mention it in Section A.3.

##### A.1 Whole Genome vs Gene-based methods

Methods that use whole genome information sometimes rely on genome-to-genome alignments; examples include StrainEst [3], which relies on pairwise alignments, and the marker gene database construction step in BIB [4] (discussed below). In general, there is a limitation that applies to *any* method that relies on a genome-to-genome alignment, because they can be confounded by large-scale rearrangements and recombinations, even across variants of the same species. Indeed, these events are not uncommon in *E. coli*, a major focus of this paper. On the other hand, some multiple alignment algorithms that try to account for this issue (such as progressiveMauve [5]) simply fail to scale with input size; the problem is far from being resolved at this point in time.

Examples of whole-genome based methods that do not use genome (pairwise or multiple) alignments are Sigma [6] (which only uses read-to-genome alignments), StrainGE [7] (which uses $k$-mer frequencies) and mSWEEP [8] (which uses pseudo-alignments from either Kallisto [9] or Themisto [10]). Using whole-genome information is not foolproof; these methods come with their own sets of biases and limitations — these differences tend to stem from the manner in which each method computationally represents, distinguishes and utilizes "conserved" nucleotide patterns versus "non-conserved" ones present in the sequencing data and/or a reference database.

Another approach that avoids these issues altogether is to associate taxonomies with a single gene (e.g. 16S for Karp [11]), or collections of genes (e.g. ConStrains [12], StrainFinder [13], the inference algorithm in BIB [4] and our method ChronoStrain). Beyond using just the 16S gene, the task of finding useful collections of genes is implemented by MetaPhlAn [14], which defines a group of "core" genes for each clade. Either approach enables gene-specific variant calling to define strains (StrainPhlAn [15], which produces pile-ups and does not resolve strain genotype-specific abundances; see the discussion below about allele frequency-based methods).

Our method is built on the latter idea, with one key difference: our belief is that the genes encoding interesting phenotypic signatures (e.g. hypothesized pathogenicity of UPEC strains) are precisely those that are *not* core to the species. In fact, this is the same reason why whole-genome methods have been developed; but our approach fits nicely in between the two extremes (core-genome vs. pan-genome) and has the potential to utilize a significant — and clinically relevant — fraction of the pan-genome.

#### A.2 Read Modeling vs Allele Frequency Modeling

The phrase *read modeling* refers to methods that models reads. One choice is to directly utilize the read nucleotides, usually in a probabilistic model without any intermediate "proxies" for reads (e.g. BIB, Sigma, Karp and ChronoStrain). Another choice is to model *read mapping counts*, where mappings are typically derived using some kind of alignment tool. Different genes have different levels of conservation within a species, and mutation rates in many genes are unknown, and thus one must take care to carefully model and account for multi-mappings (e.g. mSWEEP and ChronoStrain).

The above is in contrast to *allele frequency modeling*, whereby a method will typically align reads to a collection of reference genomes, and attempt to de-convolve a SNV frequency matrix into abundance ratios (StrainFacts, StrainFinder, StrainEst fall into this category). The primary issue with operating only on allele frequencies is that the action of computing a frequency matrix causes the loss of evidence of SNV-to-SNV correlations that each read provides. Deconvolving frequency matrices into strain genome & abundance profiles is, generally speaking, statistically challenging with just reference information. A useful example to consider is the task of deconvolving a frequency matrix on $L$ loci produced by two distinct genomes at equal abundances; this special setting requires picking one of $2^{L-1}$ possible combinations of SNVs. Mathematically, this is equivalent to the *haplotype phasing problem* from the context of human genome measurements [16], for which reads are known to be extremely helpful, and population-wide correlations (e.g. linkage disequilibrium) alone generally cannot solve every instance. Furthermore, when calling alleles across whole genomes (and not gene-specific alleles), one is effectively performing genome-to-genome comparisons, and thus this approach is subject to the same challenges discussed in the previous section.

#### A.3 Genotype Learning vs Reference Calling

We use the phrase "reference calling" to describe methods that use a reference database of assemblies, and attempt to output abundance estimates of *reference labels* that are most appropriate for the latent strains in the sample. Several methods (e.g. StrainGST, StrainEst, Sigma, BIB and Karp), including ChronoStrain fits in this category. The main limitation with this approach is that it is not obvious what to do for experimental data when there are *novel* strains. (The following points are discussed in the main text, but we recapitulate them here.) It is possible that a new strain may not deserve any *one particular* reference label; it may be mixtures of different known variants across loci, or it may contain previously unobserved variants. Typically, one digs further using allele frequency estimators such as StrainPhlAn and StrainGR (which is packaged with StrainGST), bringing us back to the allele frequency modeling setting. Another alternative is to output mixtures of labels through a posterior distribution (BIB and ChronoStrain); point estimates are also possible (StrainGST, Sigma, Karp), but when comparing across multiple samples, one may end up with inconsistency in the labels.

One potential advantage of several "genotype-learning" methods (ConStrains, StrainFinder, StrainFacts, which all happen to operate on allele frequencies) is that these methods attempt to jointly estimate strain-specific genotypes and abundance ratios. The word "potential" is used to emphasize the fact that this is a *much* more general task than what the reference-calling methods are attempting; thus, the inherent *dimensionality* of these problems is much higher, and there are more ways that these methods can go wrong. It is important to note that this harder problem is not limited to the allele-frequency modeling approach; by utilizing reads directly, one can treat this as a variant of an assembly problem (either *de novo* metagenomic assembly or as an analogue of a *polyploid* haplotype assembly).

ChronoStrain fits squarely into the category of reference calling, which is why it currently outputs mixtures of references, particularly on the UMB dataset. We do not attempt to learn genotypes *de novo* in this work, but we leave it as a crucial future direction as discussed in the Discussion section of the main text. To our knowledge, a bona-fide "time-series"-aware (genome or genotype) assembly algorithm across multiple metagenomic samples (e.g. beyond merging samples into one giant read set) has not yet been invented, but we imagine it would be a quite helpful contribution to the field.

##### A.4 Method-specific Comparison

We note that out of all works mentioned, ChronoStrain is the only one to explicitly encode timepoints $t$ at which samples were taken, even if several other works perform joint estimation across multiple samples (ConStrains, StrainFinder, StrainFacts). Excluding these features, exactly two methods agree with ChronoStrain for all categories: BIB and Karp. We summarize their differences at a high level.

BIB leverages read alignments to a database of core genes (genes shared by all strains of a species or higher clade) via a likelihood model on the read's nucleotides and phred scores; it is the *only* other method known to us that implements an approximate posterior probability distribution over abundance profiles. BIB has two limitations that do not extend to our method. First, its database construction of core genes is seeded by a whole-genome multiple sequence alignment (MSA) and thus has trouble scaling to species with more than a few dozen strains. Second, our method includes a position-encoding, fragment sliding-window model Eq. (1), whereas BIB does not. It was excluded in that method for the sake of computational complexity (since the method wants to fully consider reads that don't map uniquely to their database), but finding a tractable approximation (Section B.1.2) is one of our key contributions.

Karp was designed for 16S reads and builds upon Metakallisto's [17] usage of $k$-mer based 16S pseudoalignments: it was developed after observing that utilizing base quality can be helpful (ChronoStrain also includes quality information). Since it is designed for 16S and not whole-metagenomic sequencing, we do not expect it to work well (and confirmed this by running it on UMB18). A close cousin of these methods is mGEMS [18] which is an end-to-end pipeline connecting mSWEEP and a pseudoalignment output (either Kallisto or Themisto).

In the main text, we draw comparisons primarily to two methods, which to our knowledge are the current state-of-the-art. These methods are: StrainGST (an algorithm in the StrainGE package [7]) and mGEMS. mGEMS is a whole-genome method pipeline which models read pseudoalignment counts: the pseudoalignments are computed using Themisto [10] and these counts are then passed as input to a Bayesian model [8]. Besides the "whole-genome" versus "gene-based" comparison, another major difference between modeling (pseudo-)*alignment counts* and ChronoStrain's model is that the latter directly models sequencing noise. Since pseudoalignments only contain *mapping* information (queries of the form, "which genome(s) $g$ could read $r$ have come from?"), count-based models cannot capture fine-grained, quantifiable information about how much a read maps "better" to one reference genome than another. In particular, this directly explains one of ChronoStrain's advantages: estimating abundances of *low-abundance* strains (Supp. Fig. S2). More explicitly stated, the connection here is that rare strains might have only a few reads in the sample, and thus their abundance estimates are more sensitive to sequencing noise, especially if modeled indirectly through (say) pseudoalignment counts.

StrainGST follows a common paradigm in bioinformatics: designing *fast*, low-memory footprint algorithms by working in $k$-mer space. It operates iteratively, by repeatedly calling successive reference labels whose $k$-mer profile has the highest "overlap" with a remaining set of $k$-mers formed by the reads. However, like allele frequencies, read-derived $k$-mer frequencies loses *some* SNV-to-SNV correlations, and there is a natural tradeoff. If $k$ is too large, then the speed/memory advantages becomes negligible and the algorithm loses robustness to sequencing noise. If $k$ is too small, one sacrifices some ability to resolve long-range, multi-locus correlations, especially for strains at perpetual low-abundance. As a consequence, untangling the sequences and abundances of multiple similar, co-abundant strains becomes challenging. In theory, the following latent information ought to be inferrable from raw data, but becomes extremely challenging if one uses algorithms that are lossy, which is the case with allele or $k$-mer counts. These are: (1) correlation in time-series (*which SNVs are correlated across time?*), and (2) correlation between loci (*which SNVs occur on the same molecule?*). It is an algorithmic challenge to efficiently resolve both simultaneously, particularly for low-abundance strains, without resorting to sequencing depths likely far beyond what is theoretically required. This is precisely the type of scenario where Bayesian methods — such as ChronoStrain — tend to shine, and these concerns helped shape our algorithmic design.

#### A.5 Methods Excluded from Analysis

We remark upon tools mentioned above that did not make it into our semi-synthetic benchmark. First, BIB's database construction did not scale well to the scale of genomes being used in the semisynthetic scenario. Since ChronoStrain's database requires one to specify marker seeds without worrying about copy number or homologies, it took ~2 hours to construct the full database of ~5400 *Enterobacteriaceae* genomes and $\frac{1}{2}$ hour to agglomerate this into the final 1225 at 99.7% marker nucleotide identity. Furthermore, the original StrainEst paper ran ConStrains [12] and Sigma [6] for its own comparisons. Unfortunately, in that work, the authors found that these two tools fail to properly learn sub-species mixtures of *E.coli* and/or relies on very high coverages, and thus we do not expect them to perform better than what StrainEst can already accomplish. Furthermore, we could not run ConStrains to completion due to its reliance on outdated software.

We also made an attempt to run at least one estimation tool that provided genotype deconvolutions jointly with abundance estimates. In particular, we ran StrainFacts [19], which is a relatively new allele-based method, using the pipeline documented in the manuscript. Running this method on the semisynthetic dataset came with a caveat: the main StrainFacts algorithm (which outputs a maximum a posteriori estimator) requires specifying a value $k$ which represents the desired number of genotypes. Since we actually mixed the synthetic reads with real reads, this value could not be determined.

Instead, we ran it on **only** the simulated reads, with $k = 6$. Note that the method attempts to output abundance estimates of potentially *novel* combinations of alleles, which has no guarantee of perfectly matching our ground truth strains. To engineer a fair metric for this purpose, we calculated the following, adjusted $\ell^1$ error metric

$$\text{Error}(\hat{x}) = \min_{\pi \in S_6} \sum_{t \in \mathcal{T}} \sum_{i=1}^{4} |x_t(i) - x_t(\pi(i))|$$

where $S_6$ is the set of all permutations of six elements, $x$ is the ground truth abundance ratio, and $\hat{x}$ is the abundance estimate. This is the same error metric applied to the other methods, but minimized across all ways to mix-and-match the inferred genotypes with the ground truth strains. Unfortunately, it consistently provided poor results (TV close to 1.0, the theoretical maximum across all coverages, indicating that the algorithm did not actually learn the simulated strains) for all coverages.

This suggests that allele deconvolution methods cannot be directly compared to other types of methods using the simulated benchmarks. For instance, it is quite possible that StrainFacts learned at least one "fuzzy" genotype (mixtures of alleles per loci) that was a mixture of the simulated strains. This is problematic, since it is rather unclear how the six fuzzy genotypes relate to the six synthetic genotypes, and since it is not clear how to further untangle these into individual abundances. (Our main hope was that the cross-sample correlation provided sufficient information to resolve non-fuzzy genotypes from the allele frequency matrix.) Another potential issue might have been that the software was designed using a bi-allelic assumption, whereas the true synthetic genotypes — and the underlying database — are multi-allelic after multiple genome alignment. Keeping these issues in mind, we stopped short of re-engineering parts of the code ourselves. The core problem of allele deconvolution is a strictly harder one than what we are after (after incurring some information loss by converting reads into allele counts, as discussed in Section A), so our benchmarks' assumptions/requirements may have been too optimistic.

### B Supplemental Methods

#### B.1 Objective Function

The posterior that we are after is $P(X, Z \mid R)$ (conditional on $R$ aligning to the database), where $R$ is the subcollection of reads for which we condition on as originating from our markers. As mentioned in the main text, ChronoStrain implements Automatic Differentiation Variational Inference (ADVI) [20], although any appropriate inference algorithm can take its place. A core ingredient of ADVI is the Monte-Carlo estimate to the Evidence Lower Bound objective (ELBO):

$$\begin{aligned} \mathsf{ELBO}(\varphi) &= \mathbb{E}_{(X,Z)\sim q_\theta}[\log p(R, X, Z)] + H[q_\varphi] \\ \rightarrow \widehat{\mathsf{ELBO}}(\varphi) &\approx \left(\frac{1}{M}\sum_{m=1}^{M} \log p\left(R, \widetilde{X}^{(m)}, \widetilde{Z}^{(m)}\right)\right) + H[q_\varphi] \end{aligned}$$

where $q_\varphi$ is an approximating likelihood function parametrized by $\varphi$, $(\widetilde{X}^{(m)}, \widetilde{Z}^{(m)})$ indexed by $m$ are i.i.d. samples from $q_\varphi$, and $H(\cdot)$ is the Shannon entropy function. Note that the above is not "stochastic optimization" in the commonly understood sense of Machine Learning literature [21], since we do not sub-sample the reads for optimization (we do a full pass on the entire dataset). Subsampling would help us scale the optimization to much larger datasets, but the above calculation helps minimize variability across optimization seeds.

The core principle of ADVI is that one maximizes this function by pushing it through standard, automatic-differentiation algorithms as a black box. Since such a heuristic would call for evaluating this function many times — preferably using fresh samples for each iteration — it is critical to minimize the computational complexity required to estimate the ELBO. Otherwise, this becomes a rather expensive algorithm which might not run, even on high-performance computing resources. This is in terms of runtime *and* in terms of the memory required when storing all of the necessary gradients.

We assume that the likelihood of a read set $R$ given $X, Z$ can be written as the product

$$\begin{aligned} p(R \mid X, Z) &= \prod_{t \in \mathcal{T}} p(R_t \mid Y_t) && \text{(Cond. Indep. over } \mathcal{T}\text{)} \\ &= \prod_{t} \prod_{i=1}^{N_t} p(r_{t,i} \mid Y_t) && \text{(Cond. Indep. over reads).} \end{aligned}$$

Each individual term can be expressed as a marginalization over fragments, e.g.

$$p(r_{t,i} \mid Y_t) = \sum_{f_{t,i}} p(r_{t,i} \mid f_{t,i}) p(f_{t,i} \mid Y_t).$$

Dropping the subscripts $(t, i)$ from $r$ and $f$ for readability, we expand this according to the model described in Methods:

$$\begin{aligned} p(r \mid Y_t) &= \sum_{f} p(r \mid f) \sum_{\ell} p(f \mid Y_t, \ell) p(\ell) \\ &= \sum_{f} p(r \mid f) \sum_{\ell} p(\ell) \left( \frac{\sum_{s \in \mathcal{S}} Y_t(s) n_{f,s}^{(\ell)}}{\sum_{\hat{s} \in \mathcal{S}} Y_t(\hat{s}) n_{\hat{s}}^{(\ell)}} \right) \\ &= \sum_{s \in \mathcal{S}} Y_t(s) \sum_{f} p(r \mid f) \left( \sum_{\ell} p(\ell) \frac{n_{f,s}^{(\ell)}}{\sum_{\hat{s} \in \mathcal{S}} Y_t(\hat{s}) n_{\hat{s}}^{(\ell)}} \right) \end{aligned}$$

This expression for $p(r \mid X_t)$ looks complicated, but it can be broken down into two major pieces.

1. The calculation of fragment-to-error likelihoods $\varepsilon_{r,f} \overset{\text{def}}{=} p(r \mid f)$, and
2. The calculation of the (weighted) strain-specific fragment frequencies

$$\omega_{t,f,s} \overset{\text{def}}{=} \sum_{\ell} p(\ell) n_{f,s}^{(\ell)} \left( \frac{1}{\sum_{\hat{s} \in \mathcal{S}} Y_t(\hat{s}) n_{\hat{s}}^{(\ell)}} \right)$$

In plain English: the first piece characterizes the *error likelihood of reads* enumerated across the database, and the second piece encodes the *genetic diversity* of each member of the database. The inverse term (containing the sum indexed by $\hat{s}$) can be understood as a correction that accounts for the bias induced by the choice of marker seeds. Roughly speaking, in the posterior distribution, this term ensures that strains aren't unfairly boosted just by the pure virtue of having more marker sequences.

Let $\mathcal{F}$ be the collection of all possible fragments in the model. Algorithmically, if we are given both pieces in the form of two matrices

$$W_t = \left(\omega_{t,f,s}\right)_{f \in \mathcal{F}, s \in \mathcal{S}} \qquad \text{and} \qquad E_t = \left(\varepsilon_{r,f}\right)_{r \in R_t, f \in \mathcal{F}}$$

then the likelihood computation of the reads at timepoint $t$ can be reduced to the evaluation of the product $E_t W_t Y_t$ across $t \in \mathcal{T}$. Symbolically, this is a simple linear algebraic operation, and thus easily black-boxed using standard tensor libraries.

In practice, however, the matrices $W$ and $E$ can be extremely large. Furthermore, $E$ can be pre-computed but $W$ cannot, since the latter explicitly depends on the $Y_t$'s which we are trying to estimate. The key observation is that for biologically plausible models (e.g. variants of markers need to be somewhat similar), $W$ can be approximately decoupled from $Y_t$ via a sparse sum and $E_t$ is "close" to being sparse, in the sense that all but $O(S \times |R_t|)$ entries are vanishingly close to zero. We carefully designed heuristics for sparsely approximating $E_t$ (Section B.1.1) as well as $W$ (Section B.1.2), which are key ingredients for scaling up our model to accommodate a large database of ∼2000 strain clusters or more.

##### B.1.1 Per-read error likelihood calculation

In this section, we explain how we estimated the sparse matrix $E_t$. The overall goal here is to identify which rows $f \in \mathcal{F}$ support the row vector corresponding to read $r_{t,i}$. For the sake of exposition, we drop the subscripts and write $r = (\zeta, q)$ to represent a generic read, where $\zeta$ is the read's nucleotide sequence. Expand

$$\begin{aligned} \varepsilon_{r,f} &= p(r \mid f) \\ &= \sum_{\text{Alignments } \mathcal{A}} p(\zeta \mid f, \mathcal{A}) \times p(\mathcal{A} \mid f; q) \end{aligned} \tag{S.1}$$

for the error likelihood, which includes marginalization over $\mathcal{A}$. Instead of exhaustively listing out all theoretically plausible $f$, we narrowed down the search using an alignment-based heuristic.

We restrict the calculation to candidate fragments $f$ and alignments $\mathcal{A}$ for which this likelihood value is significantly larger than their alternatives. More precisely, this heuristic assumes the right hand side of Equation S.1 is dominated by several orders of magnitude by a single term:

$$\varepsilon_{r,f} \approx p(\zeta \mid f, \mathcal{A}^*) \times p(\mathcal{A}^* \mid f; q)$$

where $\mathcal{A}^*$ is an optimal global alignment between $f$ and $\zeta$. For those reads $r$ for which all of the above products are sufficiently small ($< e^{-500}$), we simply round $\varepsilon_{r,f}$ to zero. Tuning this threshold controls the sparsity of the matrix.

Note that this truncation applies on a *per-fragment* basis for each read, so we must search for all candidate fragments $f$ which align well to $r$. To carry out this strategy, we aligned the filtered reads to the marker database. We use the same parametrization as in the filtering step – but this time configured to output all available alignments instead of just the best one. We remark that these exact parameters tell the alignment program to return pairs $(\mathcal{A}, f)$ which (more or less) optimize the log-likelihood $\log_2 p(\mathcal{A} \mid f; q)$ amongst feasible alignments $\mathcal{A}$ of the read $r = (\zeta, q)$ to fragments $f$ of the database.

Since speed is less of a concern than in the filtering step operating on *all* of the reads, by default we use *bowtie2* [22]. This choice differs from that of the filtering step (Methods - Read Filtering) because we found that this tool returns a more comprehensive set of alignments (`bowtie2 -a` versus `bwa-mem2 mem -a`) even after tuning various parameters and accounting for tool-varying quirks in the resulting SAM outputs[8]. If the alignment clipped nucleotides of marker $m$, we re-included those bases into the alignment without indels; this is to ensure that we properly model the *whole* read. The aligned marker's substring, with gaps removed, was included in the model as a candidate fragment $f$ (the support of $r$).

##### B.1.2 Fragment frequency

In this section, we describe how we compute the sparse matrix $W_t$, whose entries are $\omega_{t,f,s}$. Here, $s$ denotes an arbitrary strain, but we may now assume that $f$ is a supporting fragment of $E_t$ for some $t \in \mathcal{T}$, since we only wish to describe non-zero entries in the product of the two matrices. We rewrite each entry $\omega_{f,s}$ for convenience:

$$\omega_{t,f,s} \stackrel{\text{def}}{=} \sum_{\ell=0}^{\infty} p(\ell) n_{f,s}^{(\ell)} \left( \frac{1}{\sum_{\hat{s} \in \mathcal{S}} Y_t(\hat{s}) n_{\hat{s}}^{(\ell)}} \right).$$

[8]However, we note that `bwa-mem2` provides a more amenable SAM output, since it excludes redundant alignments that yield the same fragment (sequence of matches/mismatches/insertions/deletions, which is more specific than the CIGAR string). For this reason, neither tool is perfect for our use case. We opted to use the computationally expensive yet more "sensitive" one.

Note that $n_{f,s}^{(\ell)} = 0$ if $\ell < |f|$, since each fragment $f$ can only be induced by windows at least as long as $f$. There is some room for truncation in the above sum. The negative binomial distribution is heavily concentrated using the parameters we set. They are in fact near-Gaussian, and taking $\ell$ ranging from $0$ to $\mu + 2\sigma$ captures roughly $98\%$ of $\ell$'s probability mass.

To evaluate $n_{f,s}^{(\ell)}$, we ran `bwa fastmap` to search for exact matches of $f$ within the database. Since the only way $f$ can be induced by a window longer than $|f|$ (by removing padded bases) is if it maps to an edge on a marker, we take

$$n_{f,s}^{(\ell)} = \sum_{m \in \mathcal{M}_s} \sum_{h \in H_f} \mathbb{1}\{h \text{ maps } f \text{ to the edge of } m\} \vee \mathbb{1}\{|f| = \ell\}$$

where, as a reminder, $\mathcal{M}_s$ is the set of markers in strain $s$, $H_f$ is the set of exact-match mappings to the marker sequences, and $\vee$ denotes the binary OR operator.

Next, we consider the denominator. Note that we can expand the expression by explicitly computing $n_{\hat{s}}^{(\ell)}$:

$$\begin{aligned}\sum_{\hat{s} \in \mathcal{S}} Y_t(\hat{s}) n_{\hat{s}}^{(\ell)} &= \sum_{\hat{s} \in \mathcal{S}} Y_t(\hat{s}) \sum_{m \in \mathcal{M}_{\hat{s}}} (|m| + \ell - 2\beta + 1) \\ &= \left( \sum_{\hat{s} \in \mathcal{S}} Y_t(\hat{s}) L_{\hat{s}} \right) + (\ell - 2\beta + 1) \left( \sum_{\hat{s} \in \mathcal{S}} Y_t(\hat{s}) |\mathcal{M}_{\hat{s}}| \right)\end{aligned}$$

where $L_s$ is the total nucleotide length across all markers of $s$. These terms have simple interpretations. The first sum is the (weighted) mean total marker content across strains in the database. The second term is the total number of padded window positions that we introduce; in particular, $\beta$ is the budget parameter that determines how willing we are to incorporate edge-mapped reads into our model.

Since with high probability $\ell$ cannot be too large (e.g. consider the rationale behind the suggested truncation $\ell \leq \mu + 2\sigma$), and assuming that the total *length* of markers for each strain typically greatly exceeds the total number of padded bases (which is generally true in our case), we provide the approximation

$$\frac{1}{\sum_{\hat{s} \in \mathcal{S}} Y_t(\hat{s}) n_{\hat{s}}^{(\ell)}} \approx \frac{1}{\sum_{\hat{s} \in \mathcal{S}} Y_t(\hat{s}) L_{\hat{s}}}$$

In particular, this expression does not depend on $\ell$, and thus we end up with

$$\omega_{f,s} \approx \left( \frac{1}{\sum_{\hat{s} \in \mathcal{S}} Y_t(\hat{s}) L_{\hat{s}}} \right) \sum_{\ell = |f|}^{\mu + 2\sigma} p(\ell) n_{f,s}^{(\ell)}$$

In log-likelihood space (which we need for numerical stability when computing the ELBO), the expression is

$$\log \omega_{t,f,s} \approx \log \left( \sum_{\ell = |f|}^{\mu + 2\sigma} p(\ell) n_{f,s}^{(\ell)} \right) - \log \left( \sum_{\hat{s} \in \mathcal{S}} Y_t(\hat{s}) L_{\hat{s}} \right)$$

This heuristic reasoning gives us an algorithmic advantage, namely that the above expression is now *decoupled*:

- the complicated summation over $\ell$ does not depend on $Y_t$, so it can be *precomputed*, and

- the second term is easily computable via a single vectorized operation during ADVI.

Furthermore, from a purely algorithmic perspective, this expression is rather nicely interpretable. The first term can be thought of as the "bag-of-words" component from a standard topic model ($\log \mathbb{P}(\text{word} = f \mid \text{topic} = s)$), and can be thought of as to the approximate (truncated) expectation $\widetilde{\mathbb{E}}_\ell[n_{f,s}^{(\ell)}]$. The second term corrects for database-specific bias; it penalizes strains that are over-represented in terms of marker content. Indeed, having too few markers is *not* an intrinsic property of the strain, it is a property of the database being used.

#### B.2 Reads with mate pairs

For mate pairs $\vec{r} = (r_1, r_2)$, we modify the model described in the previous sections. We introduce a dependency between these two reads by modeling the pair as occurring jointly given a *paired* fragment $\vec{f} = (f_1, f_2)$:

$$p(\vec{r} \mid X_t) = \sum_{\vec{f} \in \mathcal{F}_2} p(\vec{r} \mid \vec{f}) p(\vec{f} \mid Y_t)$$

We model mate pairs as being conditionally independent given $\vec{f}$:

$$\varepsilon_{\vec{f},\vec{r}} \stackrel{\text{def}}{=} p(\vec{r} \mid \vec{f}) = p(r_1 \mid f_1) p(r_2 \mid f_2) = (\varepsilon_{f_1,r_1})(\varepsilon_{f_2,r_2}).$$

The paired fragment occurs at random from the multi-set of all fragment pairs (which we denote using $\mathcal{F}_2$) that appears simultaneously in the same strain:

$$\mathcal{F}_2 = \{(f_1, f_2) \in \mathcal{F} \times \mathcal{F} : n_{f_1,s}^{(\ell_1)} > 0 \text{ and } n_{f_2,s}^{(\ell_2)} > 0 \text{ for some } s \in \mathcal{S}, \ell_1, \ell_2\}.$$

A fully rigorous way to model the pair $\vec{f}$ would be to have it be randomly induced by all pairs of possible windows observable from a paired-end sequencer, given a sequencing library of inserts that are fragments from the population $Y_t$. Moreover, each pair of windows could be determined by a insert length (or a distribution of insert lengths from a sequencing library) and measuring a certain number of bases, while taking into account any non-marker regions that the insert might span. However, ideally we would like to write down an approximate formula that is (1) efficient to compute, (2) doesn't require prior knowledge of the library's insert lengths, and (3) makes minimal assumptions about the lengths in between marker regions that these genomic inserts would span – all while being quantatively reasonable.

To this end, we make the simplifying assumption that the fragment pair $\vec{f}$ appears proportional to its *joint* frequency, which we further assume factors nicely: $n_{\vec{f},s}^{(\vec{\ell})} = n_{f_1,s}^{(\ell_1)} n_{f_2,s}^{(\ell_2)}$, but we restrict this calculation to the special case where $f_1, f_2$ are each supported by alignments to $r_1, r_2$ based on the read-error likelihood calculations for strain $s$ (§B.1.1). To this end, we assume a simple factorized model where $\vec{f}$ is determined by a paired length $\vec{\ell} = (\ell_1, \ell_2)$:

$$p(\vec{f} \mid Y_t) = \sum_{\vec{\ell}} p(\ell_1) p(\ell_2) \frac{\sum_s Y_t(s) n_{f_1,s}^{(\ell_1)} n_{f_2,s}^{(\ell_2)} \beta_{\vec{f},s}}{\sum_{\hat{s}} Y_t(\hat{s}) \left( \sum_{f_1,f_2} n_{f_1,s}^{(\ell_1)} n_{f_2,s}^{(\ell_2)} \beta_{\vec{f},\vec{\ell},s} \right)}$$

In the above, (1) $\ell_1, \ell_2$ are iid copies of the negative binomial fragment length variable used in the previous sections, and (2) $\beta_{\vec{f},s}$ is a fixed zero-one valued parameter, where it is one if and only if the pair is observed as a paired-end read alignment to markers of strain $s$. To simplify the calculation, we make the approximation[9] $\sum_{f_1,f_2} n^{(\ell_1)}_{f_1,s} n^{(\ell_2)}_{f_2,s} \beta_{\vec{f},\vec{\ell},s} \approx L_s$. This yields the algorithm evaluating the likelihoods of mate pairs via the matrix product $E_t^{\text{pair}} W_t^{\text{pair}} Y_t$, where $\varepsilon_{\vec{f},\vec{r}}$ are the entries of $E_t^{\text{pair}}$ and

$$\log \omega_{t,\vec{f},s} \approx \log\left(\widetilde{\mathbb{E}}_{\ell_1}[n^{(\ell_1)}_{f_1,s}]\right) + \log\left(\widetilde{\mathbb{E}}_{\ell_2}[n^{(\ell_2)}_{f_2,s}]\right) - \log\left(\sum_{\hat{s}\in\mathcal{S}} Y_t(\hat{s}) L_{\hat{s}}\right)$$

are the entries of $W_t^{\text{pair}}$ whenever $f_1$ and $f_2$ arise out of alignments of some read pair $\vec{r}$, and $-\infty$ otherwise.

#### B.3 Model hyperparameters

The model inclusion prior $Z(s) \sim \textsc{Bernoulli}(\pi)$ is given the parameter $\pi = 0.001$ by default, so that for a typical database of order $10^3$ there are at most ten (or up to a small multiplicative factor) clusters in expectation of the prior. The negative binomial parameters $R_{\text{NB}}, P_{\text{NB}}$ for fragment lengths are fit from adapter-trimmed data and the marker database. In brief detail: we take the union of all fragment lengths that are induced by sliding windows of length $\ell$ across a fictitious marker of length $m$, with $bm$ padded bases on either end of the marker (as specific by $\textsc{WindowsWithPaddings}$). Using these collection of "simulated" fragment lengths, we fit a negative binomial using `statsmodels`.

For this fitting scheme, we vary $\ell$ across a range by taking an equally-spaced subsample of size 100 of read lengths after sorting them. This choice is informed by the fact that the not all reads are of equal lengths, partially due to adapter trimming but also because the sequencer may not have been run with identical settings across all samples. We choose $m$ to be the median marker length across the whole database. We take $\beta = 0.5$, meaning that for each fragment-read pair $(r, f)$ allowed by the model, the fragment $f$ aligns with at least half of the read. It is theoretically possible to specify a sample-specific negative binomial distribution; this choice was made for simplicity in the initial version of the software.

The insertion and deletion error rates $\varepsilon_{\text{ins}}, \varepsilon_{\text{del}}$, depending on whether the read was forward or reverse in the pair, are set by default to the empirical insertion and deletion error rates (on the order of $10^{-6}$) from [23] for Illumina reads. We remark that that work used a different HiSeq dataset to obtain estimates. However, the inference results of this work were not very sensitive to these two parameters within one order of magnitude. Thus, these parameters should be re-tuned if, say, the sequencing platform has an abnormally high indel error rate ($\varepsilon_{\text{ins}}, \varepsilon_{\text{del}} \geq 10^{-5}$).

#### B.4 Model trimming

To further improve the scalability of the algorithm, we trim down parts of the theoretical model that don't need to be included into the calculation. The trimming occurs in two stages. In the first stage, we remove all strain clusters $s$ which do not satisfy

$$s = \operatorname*{argmax}_{\hat{s}} p(r|\hat{s})$$

[9] A "naive" approximation would be quadratic (e.g. $L_s^2$), since the expression counts pairs. However, the linear approximation $L_s$ is more realistic. This is because the number of possible fragment pairs are endpoints of inserts from a sequencing library — even if we do not know the insert length distribution — and consequently for each "forward" fragment there are only a small number of "reverse" fragments paired to it. This is the information encoded in $\beta_{\vec{f},\vec{\ell},s}$. Even if we assume a missing constant factor, it does not appear in the gradient-based optimization, and thus it suffices to use $L_s$.

for some $r$ in the entire dataset. This is a rather conservative condition, as this only needs to occur *once* across all samples for a strain to be kept. In English, this is designed to cull strains which do not have any "decisive" evidence as given by the data and often (theoretically) results in near-zero posterior probabilities.

In the second stage, we consider the empirical read-likelihood correlation matrix $C = (C_{ij})$ indexed by $\mathcal{S}^2$ (or whatever is left over after the first stage)

$$C_{s,s'} = \widehat{\mathrm{Corr}}(\alpha_{r,s}, \alpha_{r,s'}).$$

Here, $\widehat{\mathrm{Corr}}$ indicates letting $r$ range across the existing filtered reads and computing the Pearson correlation coefficient. The term $\alpha_{r,s} = \frac{p(r|s)}{\sum_{\hat{s}} p(r|\hat{s})}$ is the normalized probability, which represents what the posterior $p(s|r)$ would be if all clusters had equal abundance. Using this matrix $C$ as a measure of similarity, we run scikit-learn's agglomerated clustering with "complete" linkage and 99% correlation cutoff, forming data-driven, "secondary" clusters beyond what was already done in the database.

In short, this stage identifies groups of strain clusters that are near-impossible to tell apart using the existing data. These are then reported in the software's output. In our figures, we unraveled these ad-hoc clusters for data interpretation, by dividing the abundances equally across all members of this secondary clustering. Users may interpret these clusters as a diagnostic of what they are un-able to estimate from the existing data. Sometimes, especially when data is sparse, the SNVs that can be used to tell apart two clusters will have zero read coverage — and the dataset will yield a high correlation $C_{s,s'}$.

#### B.5 An estimator for overall relative abundance

Our method estimates the ratio of strains $Y_t = \texttt{softmax}(X_t)$ which is normalized across the database only. Since the whole population of strains, including those species not in the database, is unknown, we derive a simple estimator for the overall relative abundance (the absolute abundance of each strain divided by total bacterial abundance in the sample) via the calculation

$$\begin{aligned}\mathbb{P}(\text{Strain} = i \mid Y_t) &= \mathbb{P}(\text{Strain} \in \mathit{DB} \mid Y_t) \times \mathbb{P}(\text{Strain} = i \mid \text{Strain} \in \mathit{DB}, Y_t) \\ &\approx \left(\frac{n_{\text{DB-chr}}}{n_{\text{marker}}}\right) \times \left(\frac{n_{\text{marker}}}{n_{\text{total}}}\right) \times (Y_t)_i. \qquad \text{(Estimator \#1)}\end{aligned}$$

Here, given a particular timepoint $t$, $n_{\text{DB-chr}}$ is the number of reads that map to all database chromosomes, $n_{\text{marker}}$ is the number of reads that map to markers, and $n_{\text{total}}$ is the total number of reads. Most importantly, this formula considers the probability that a randomly chosen strain from the (unknown) overall population at time $t$ is equal to $i$. In spirit, this ends up being very similar to the taxonomy-bin abundance estimator given in MetaPhlAn.

The goal is to estimate the above *without* performing yet another costly alignment of all reads to the reference collection (e.g. for estimating $n_{\text{DB-chr}}$). We estimate the first ratio by substituting each term with their approximate expected values; letting $r_s$ be the true overall abundances, note its cancellation:

$$\left(\frac{n_{\text{DB-chr}}}{n_{\text{marker}}}\right) \approx \frac{\sum_{s \in \text{DB}} r_s N}{\sum_{s \in \text{DB}} r_s \frac{L_s}{G_s} N} = \frac{1}{\sum_{s \in \text{DB}} (Y_t)_s \times (L_s / G_s)}$$

where $L_s$ is the total marker length of strain $s$ and $G_s$ is its total chromosomal length. For the second ratio, we simply count the number of marker-aligning reads and divide by the overall read depth.

Finally, our ELMC analysis (Figure 5, Supp. Fig S7) shows that summing the above across *E. faecalis* produced species-level abundances which agreed with Bracken within an order of magnitude. This observation makes it feasible to design an estimator that uses Bracken [24] species estimates. More precisely, to estimate the overall relative abundance of strain cluster $i$ of species $s$, one first produces a vector $\widetilde{Y}_t$ which zeroes out all entries of $Y_t$ that are not of species $s$, and re-normalizes it to sum to 1. Then, one scales this vector by the Bracken estimate $\widehat{r}_{t,s}^{\text{Bracken}}$ of species $s$ at timepoint $t$:

$$\mathbb{P}(\text{Strain} = i \mid Y_t) \approx \widehat{r}_{t,s}^{\text{Bracken}} \times (\widetilde{Y}_t)_i \qquad \text{(Estimator \#2)}$$

Algorithmically, this is a very different estimator from the first; the first only uses statistics within the framework of ChronoStrain's model and thus is more convenient to compute, but the second uses a third-party estimate of the re-scaling (ideally using a large, extensive database) which does not share the same biases as our method.

#### B.6 ELMC analysis with in-silico mutations

##### B.6.1 Experimental Setup

Using the ELMC dataset, we wished to probe the behavior of ChronoStrain and mGEMS when faced with a slightly incorrect database but on real (not simulated) reads. In particular, `demix_check` by design conservatively assigns good scores only to clusters which contain genomes both in the database and in the sample, while ChronoStrain's Bayesian model has some flexibility since it is an end-to-end model capturing sequence-level similarity. To test the two methods in a setting where the reads do not *quite* match the database (but are still reasonably close), we re-ran the methods after introducing random mutations into the database.

Note that in the semisynthetic benchmark (where we also simulated mutations), we mutated *reads*. However, we cannot mutate reads in an unbiased way like we did for the semisynthetic: if two reads came from the same strain and from the same position on the genome, both reads must contain the same mutation. Since we do not know what reads belong to what strain, an analogous experiment cannot be done. Instead, for this experiment we mutated the *database*. To do so, we first we picked exactly one isolate positively identified by mGEMS (appearing first in lexicographic order by ID) from each infant in the original run, resulting in 117 isolates total. Then, we mutated these 117 isolates' assemblies using a random nucleotide change with mutation rate $0.002$ (on average, this leaves 500 nucleotides in-between mutated sites). For both methods, we re-built the databases by combining these 117 mutated assemblies with the background 2,026 European isolates. Both methods' clustering schemes — using the same thresholds used to construct the original databases — resulted in all 117 infant isolates broken apart into separate clusters. Inference was run and interpreted using the same parameters as the original run (Methods - ELMC Analysis). In particular, ChronoStrain used $\pi = 0.001$ and interpreted using $\overline{\pi} = 0.95$ with abundance threshold 0.065. mGEMS was run with Themisto and mSWEEP at default settings, and we ran `demix_check` on all bins.

The above specifically excludes the other 204 non-chosen infant isolates, but this was done on purpose to remove a layer of complication. In reality, many of the 321 infant isolates are likely identical strains, especially if they were cultured from the same infant. If two isolates are indeed independent cultures of identical strains, we would need to mutate both of them in an identical manner to obtain comparable results. However, a priori we do not know which isolates are identical: their assemblies have different numbers of contigs, have different regions of completion, and have different mis-assemblies. This leaves room for some arbitrary, and potentially confounding, decision-making process to determine which isolates are identical strains (e.g. using PopPUNK clusters, which often groups isolates across different infants together). In addition, there is some guesswork involved in "combining" multiple contig-level assemblies into a single, unified genome so as to introduce the same set of mutations for genetically equivalent isolates. For instance, it is not clear whether re-doing assemblies using a merged collection of reads is the right approach.

##### B.6.2 Interpretation of Results

Here, we interpret the results produced by the above setup, first alluded to in the main text (ELMC analysis results) and illustrated in Main Figure 6 and Supplemental Figure S9. For mGEMS, this experiment not only tests the abundance estimates (produced by mSWEEP [8]), it also tests strain calls filtered by the QC tool `demix_check`. This tool evaluates *"confidence scores for whether the reads [...] originate from the [strain] they're assigned to"* [25], referring to mGEMS' binned-read outputs. Recall that in the semi-synthetic benchmark, all bins including those containing the ground-truth strains (prior to introducing mutations) were assigned `demix_check` scores of 4[10]. In both the original analysis and our re-run, we only retained strain calls if the `demix_check` score was 2 or below[11]. Using the same hyper-parameters and post-inference thresholds as in the main results (ChronoStrain $\overline{\pi} = 0.95$, abundance ratio $\geq 0.065$; mGEMS `demix_check` $\leq 2$, abundance ratio $\geq 0.01$), mGEMS calls decreased from 117/117 to only 45/117 ELMC isolates. In contrast, ChronoStrain stayed almost identical, calling 108/117 to calling 109/117 isolates with the slightly mutated genomes (Figure 6a and Supp. Figure S9, panel a3/b3/c3).

The drastic reduction in strain calls by mGEMS is expected behavior and is the intended design of `demix_check`. Indeed, the `demix_check` scores decayed in value across the board (Figure 6, panel b); relaxing the `demix_check` thresholds effectively re-includes all strains whose strain ratio is above $0.01$. In particular, when one admits all `demix_check` scores, mGEMS calls 93/117 isolates (Figure S9, panel c3). However, by doing so mGEMS now calls a median of six strains per sample (6:1 ratio of median number of strain calls between mGEMS and CS for mutants in the database, (Figure S9, panel c1). Thus, in this setting filtering by `demix_check` scores naturally becomes less specific for the isolates, even if there is a database genome which is fairly close to the genome present in the sample.

In contrast, ChronoStrain's calls are robust to slight mismatches between the database and the metagenomic sample. Furthermore, even the slight increase in calls (108 to 109) can be explained: due to the mutations introduced into the database, each of the 117 isolates were assigned its own unique cluster (in both methods). Separately, we observed that a more granular database encouraged higher specificity, when we ran ChronoStrain using a database clustering threshold of $0.9999$ similarity (Supp. Fig. S8). Again, we emphasize that all the inference priors and parameters used for thresholding for ChronoStrain were the same in this mutation experiment as in the original run, and the results are essentially unchanged. This further demonstrates the robustness and real world utility of ChronoStrain to call the best strain cluster even when the reference genomes in that cluster are not identical to the genome that generated the reads.

---

[10] A score of 4 means *"The query-ref distances are greater than the threshold, and are closer to the median between-cluster distance than the median within-cluster distance"* [26].

[11] A score of 2 means *"The query-ref distances are greater than the ref-ref distances from the same cluster, but are within the threshold"* [26]

26. Thorpe, H. *demix-check 65dcad6b316724b34b380472abec216789d8a4b3* https://github.com/harry-thorpe/demix_check.
