## Supplemental Data - Trajectories for "Strain tracking with uncertainty quantification"

#### Contents

1 UMB – timeseries comparison per participant 1

2 ELMC – timeseries plots per infant 26

#### 1 UMB – timeseries comparison per participant

Here, we provide time-series analysis results for both ChronoStrain (panels **a**, **b**, **c** in each plot) and StrainGST (panels **d**, **e**, **f**) for all UMB participants. Each $y$-axis in the time-series trajectories is the *overall* relative abundance, and not the *database-normalized* abundance which is method-specific. For instance, $10^{-2}$ refers to the relative abundance equal to a hundredth of the sample's bacterial cell count. All credible intervals were drawn using 5,000 posterior samples using ChronoStrain. The trajectories (panels **c**, **f**) are estimates from stool. (Note: We remind the reader that the phylogroup colorings are purely plot-specific annotations and were not used for inference.)

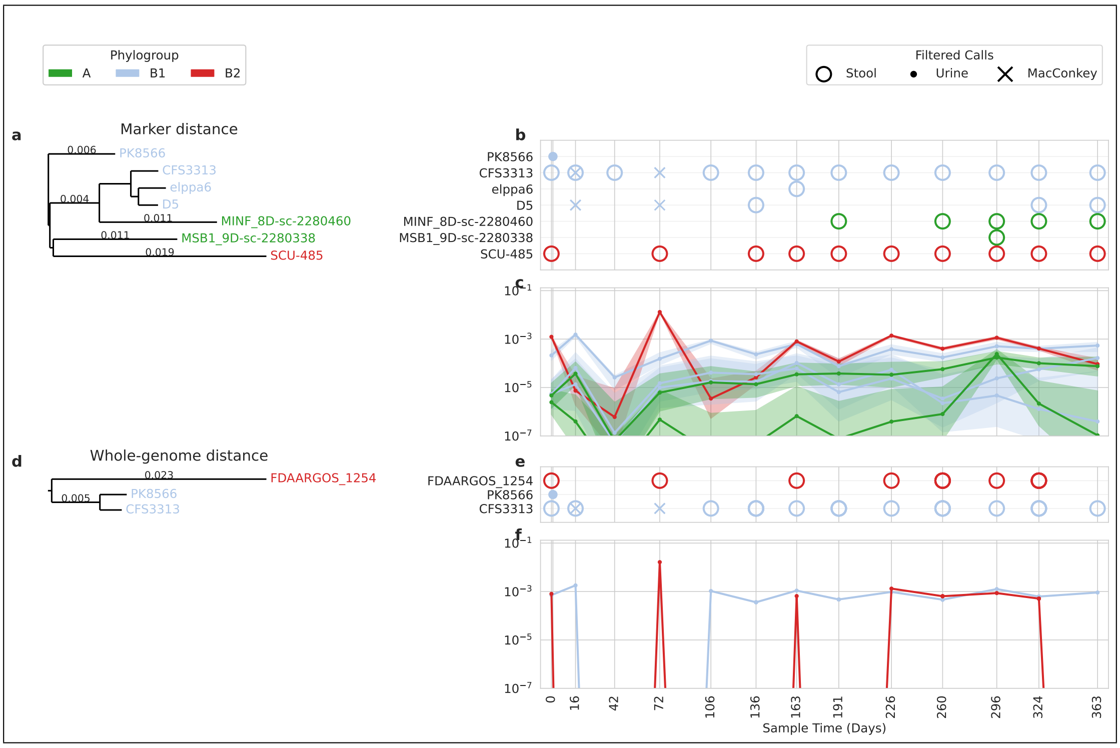


Figure U1: Participant UMB01

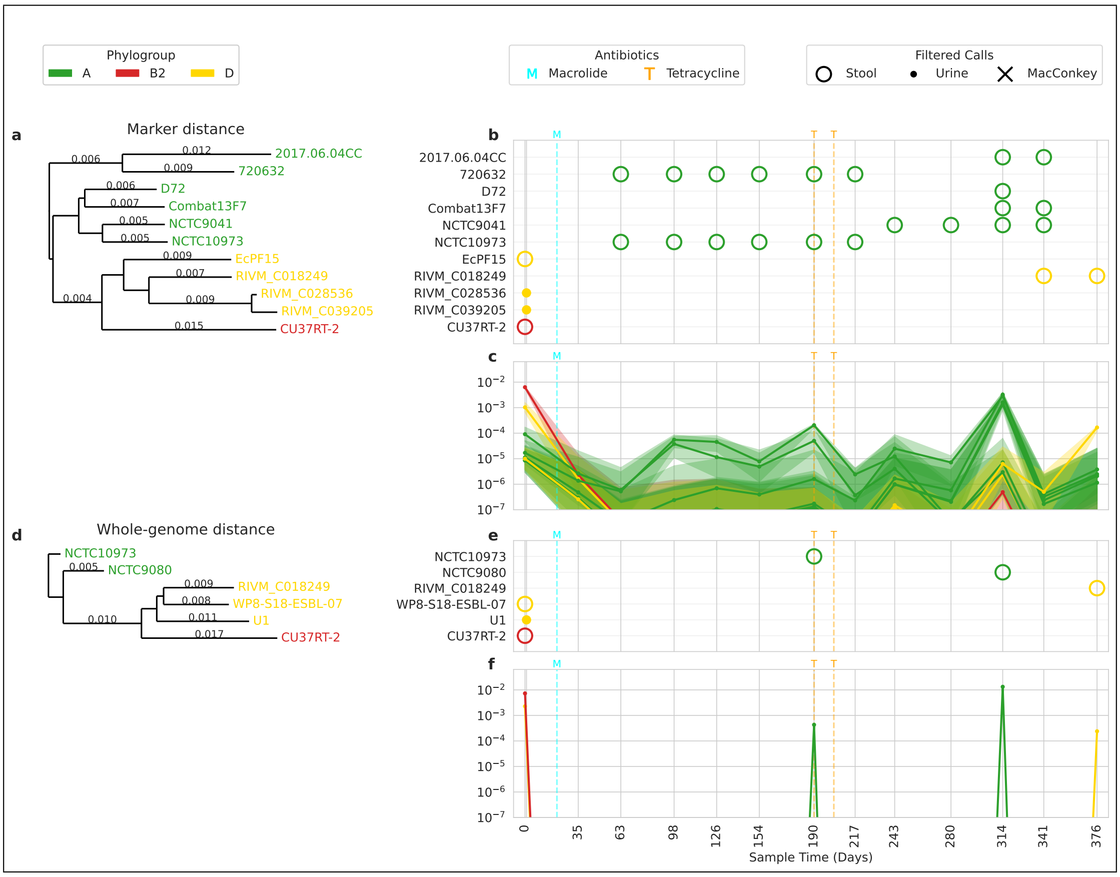


Figure U2: Participant UMB02

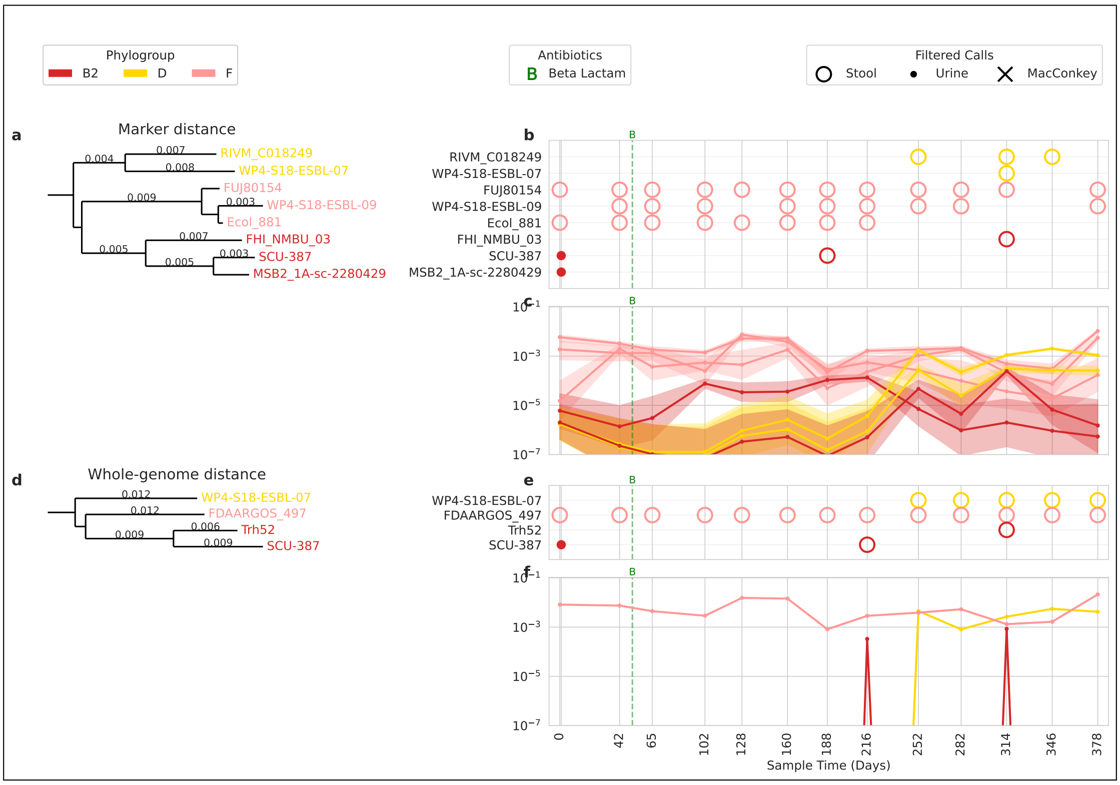


Figure U3: Participant UMB03

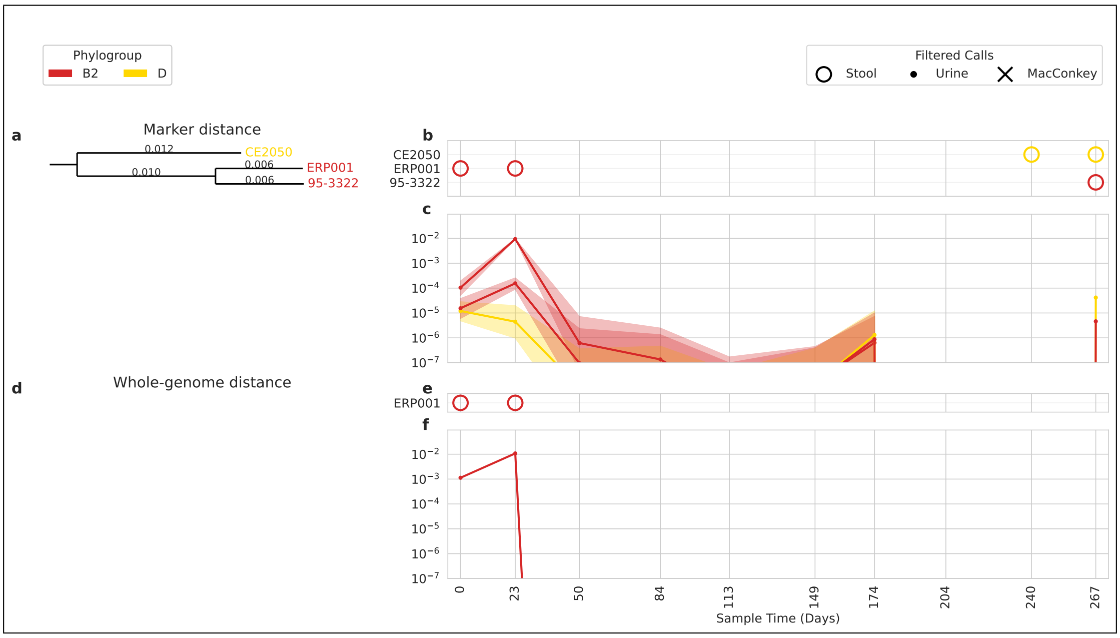


Figure U4: Participant UMB04

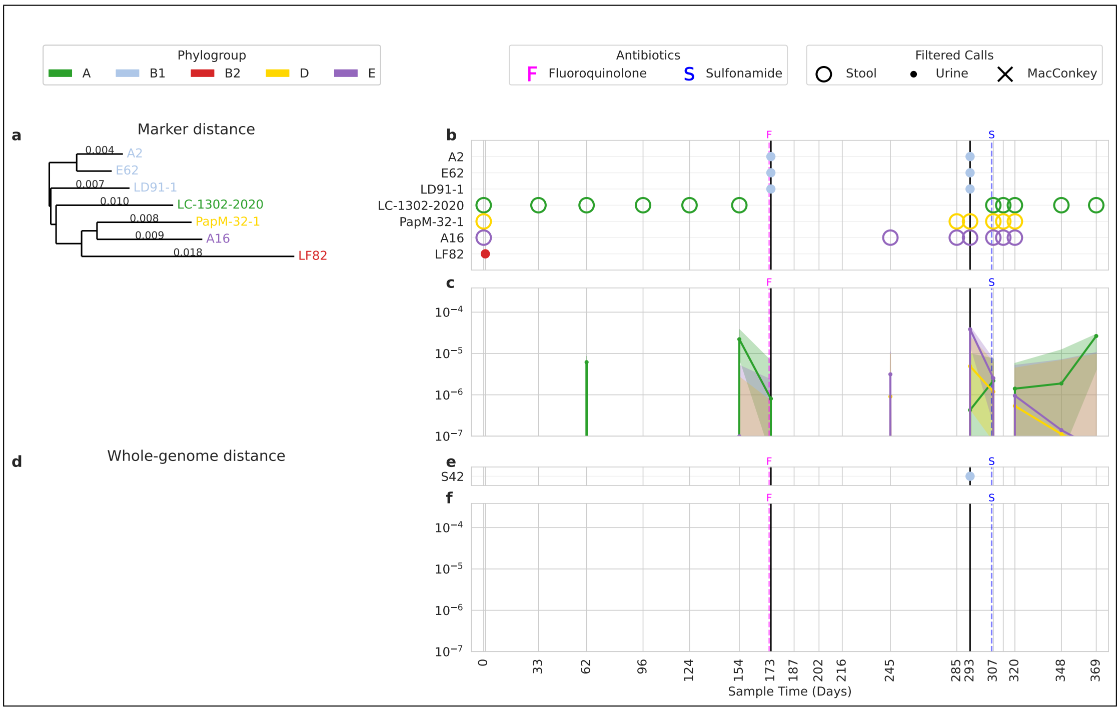


Figure U5: Participant UMB05

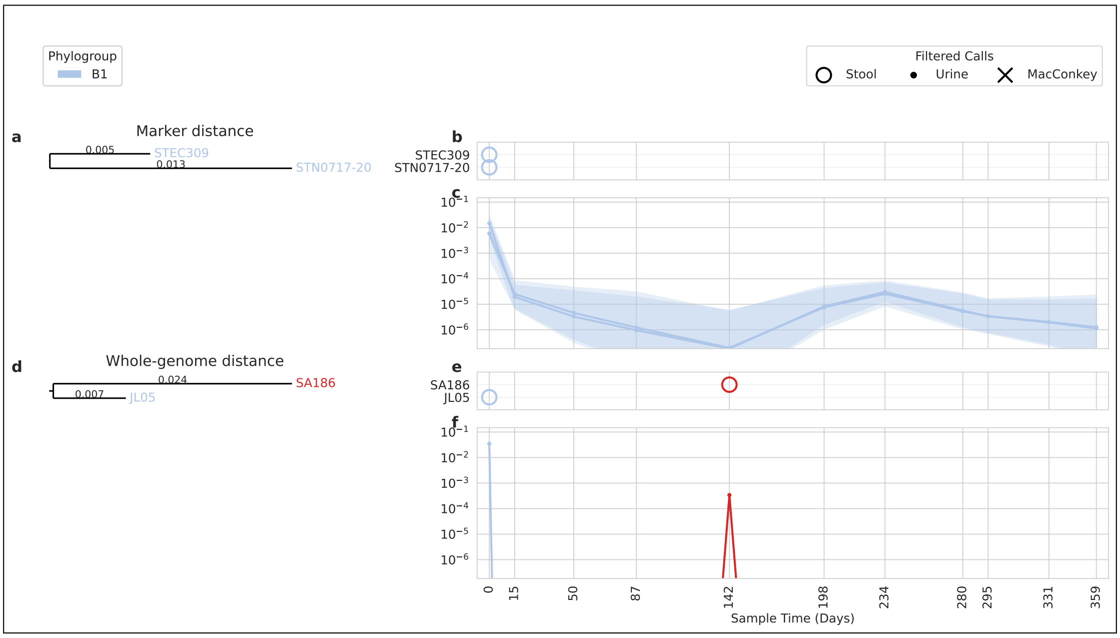


Figure U6: Participant UMB06

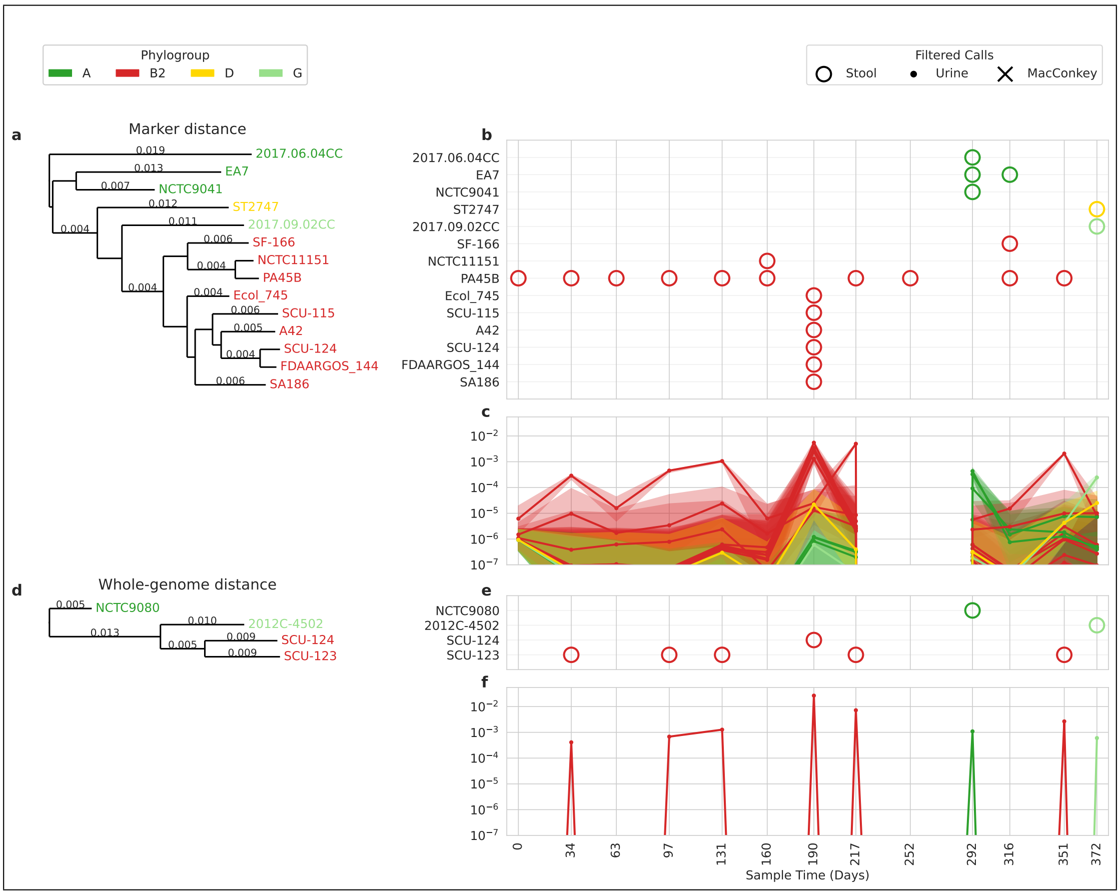


Figure U7: Participant UMB07

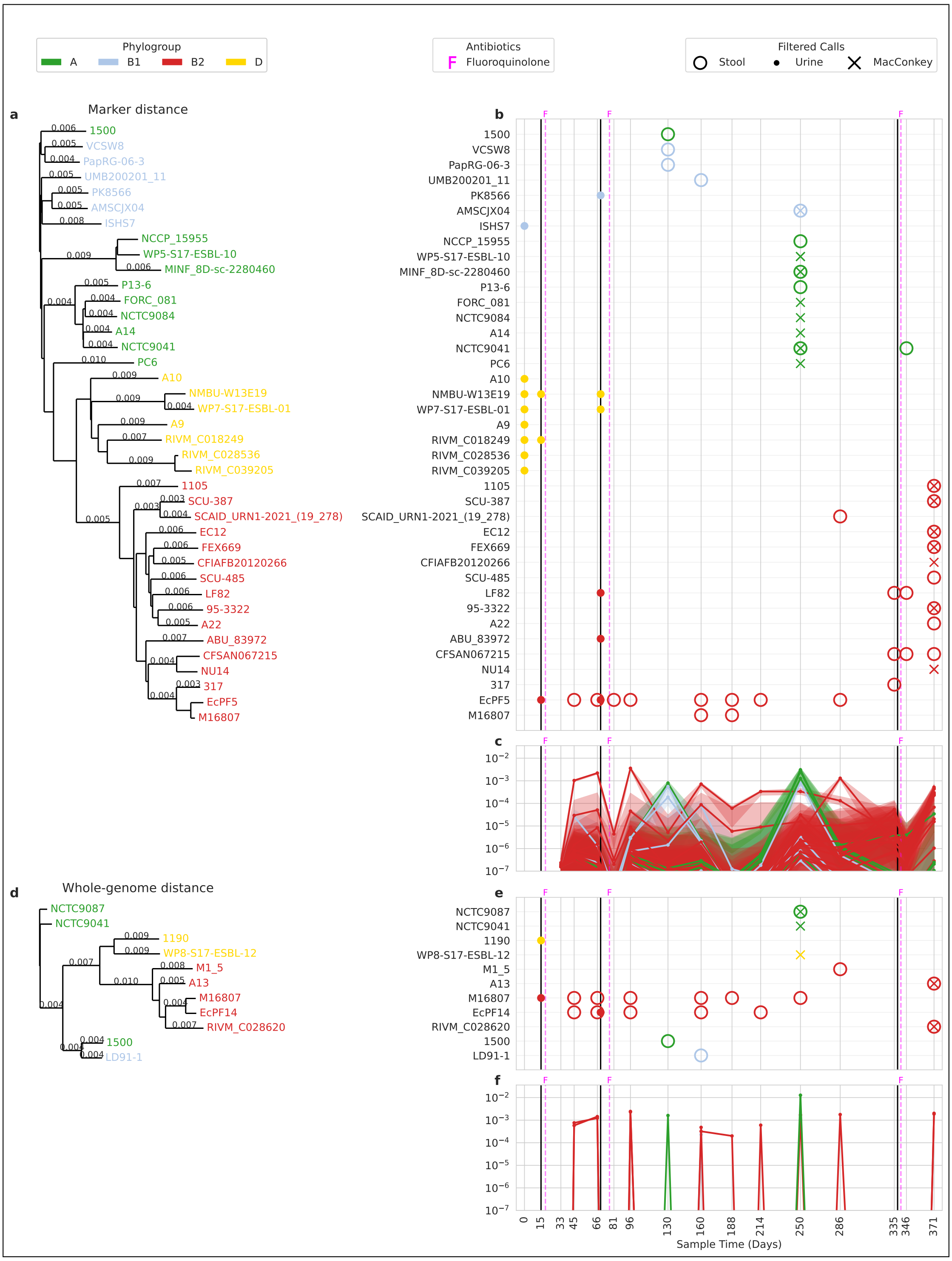


Figure U8: Participant UMB08

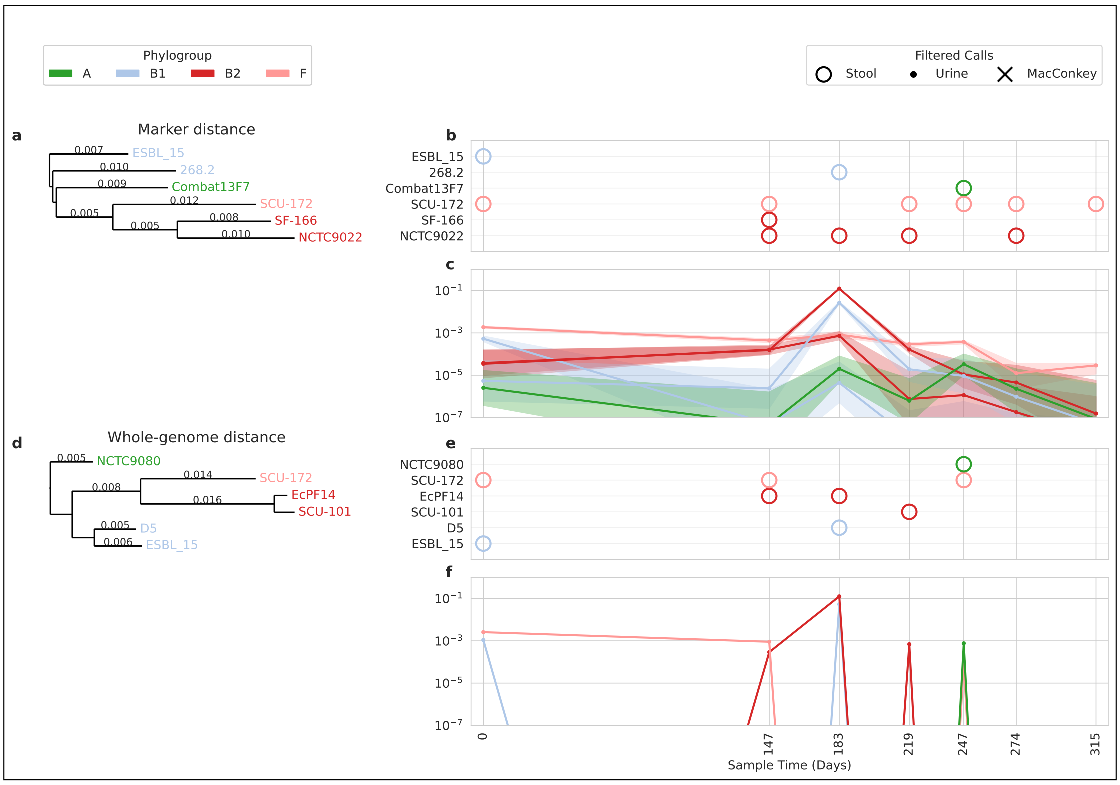


Figure U9: Participant UMB09

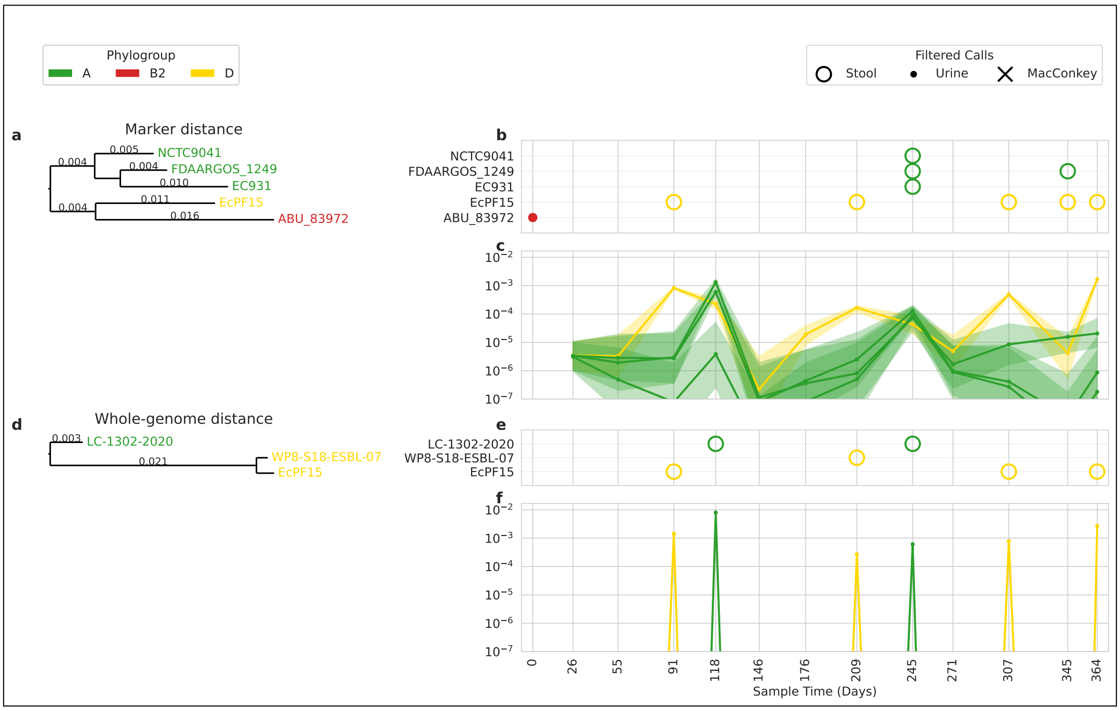


Figure U10: Participant UMB10

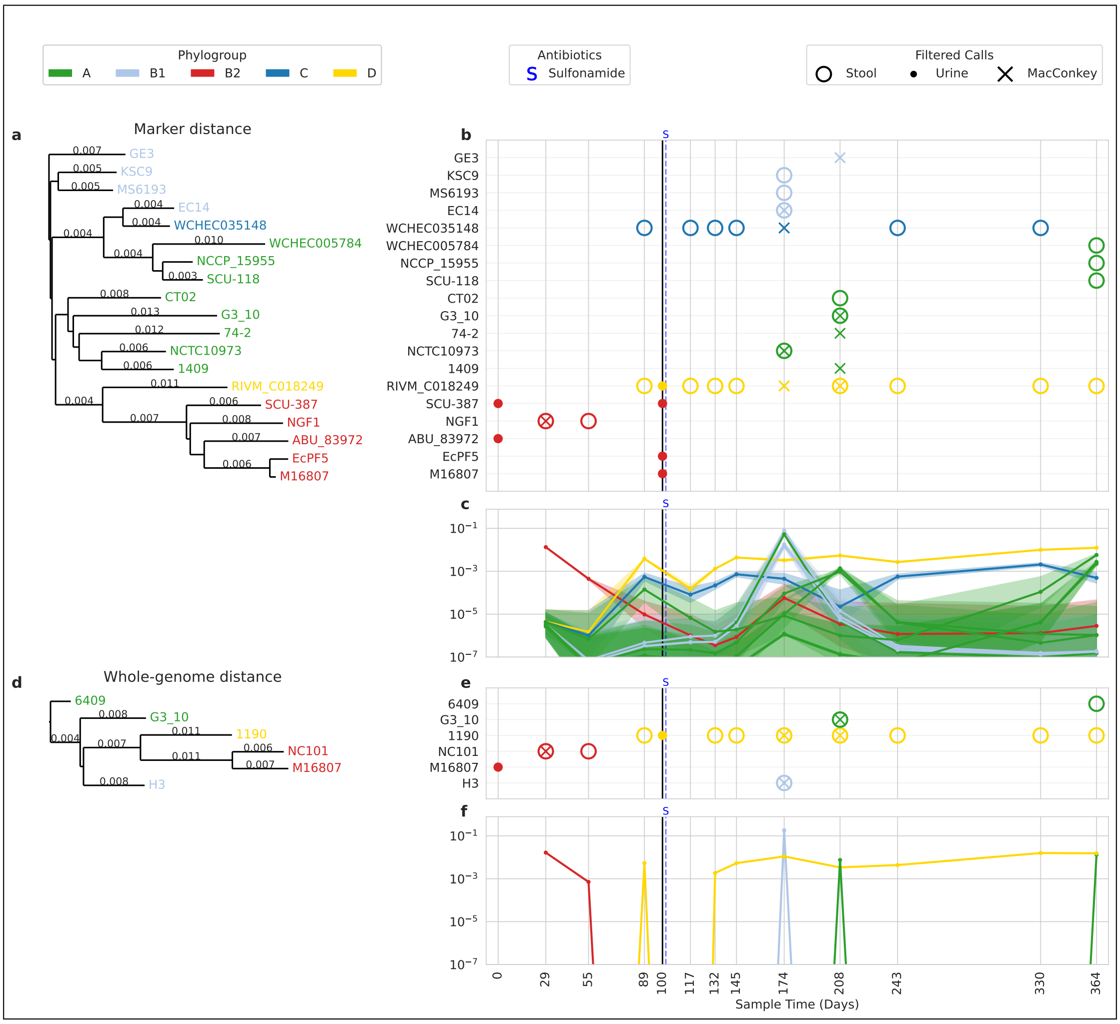


Figure U11: Participant UMB11

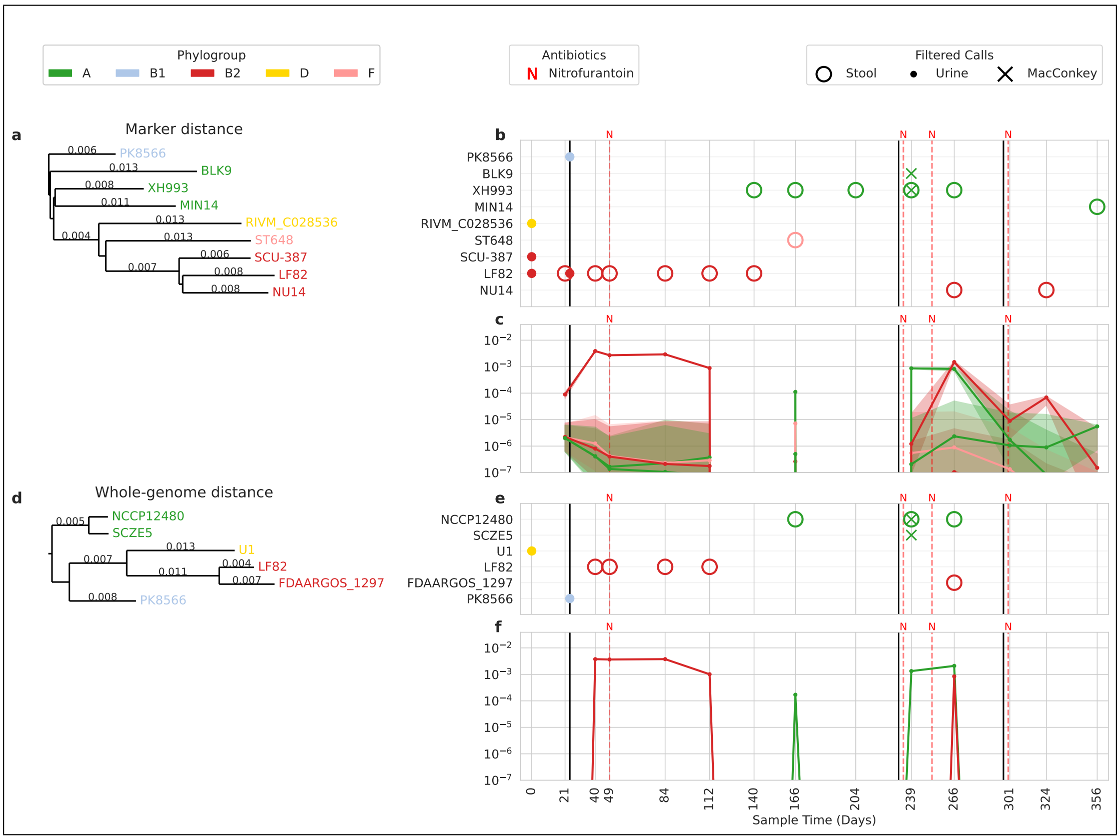


Figure U12: Participant UMB12

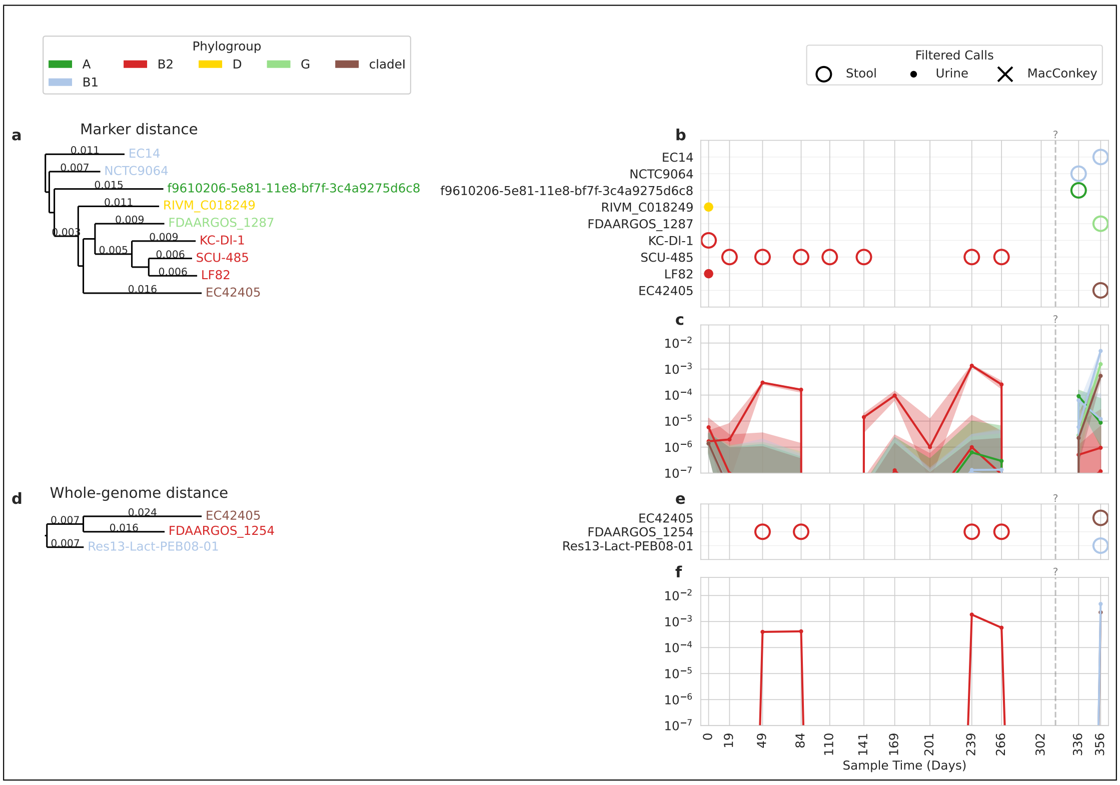


Figure U13: Participant UMB13

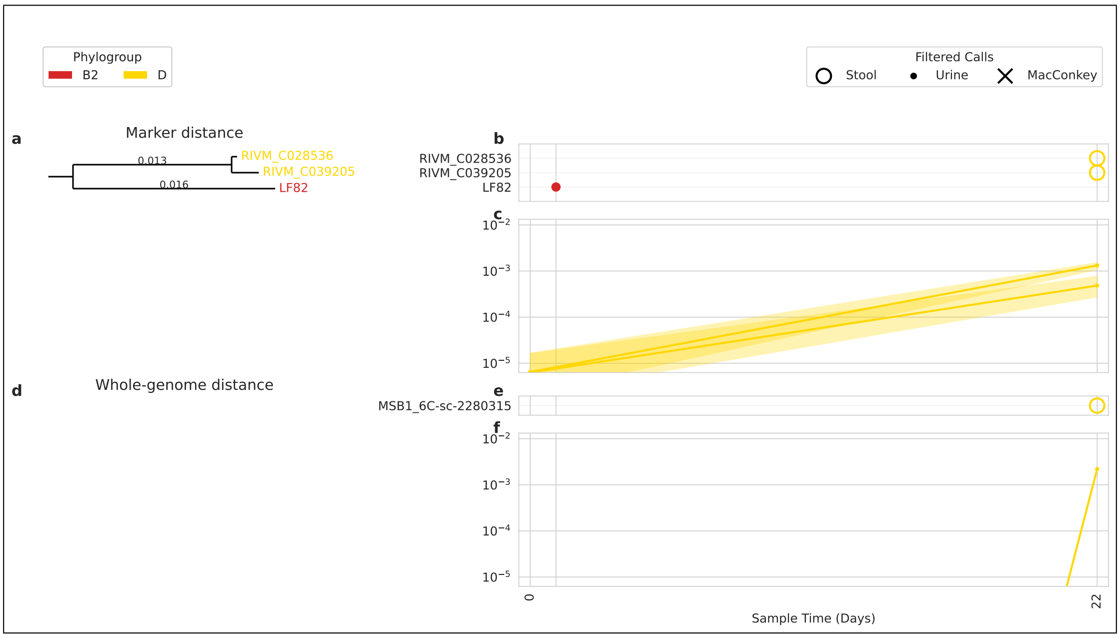


Figure U14: Participant UMB14

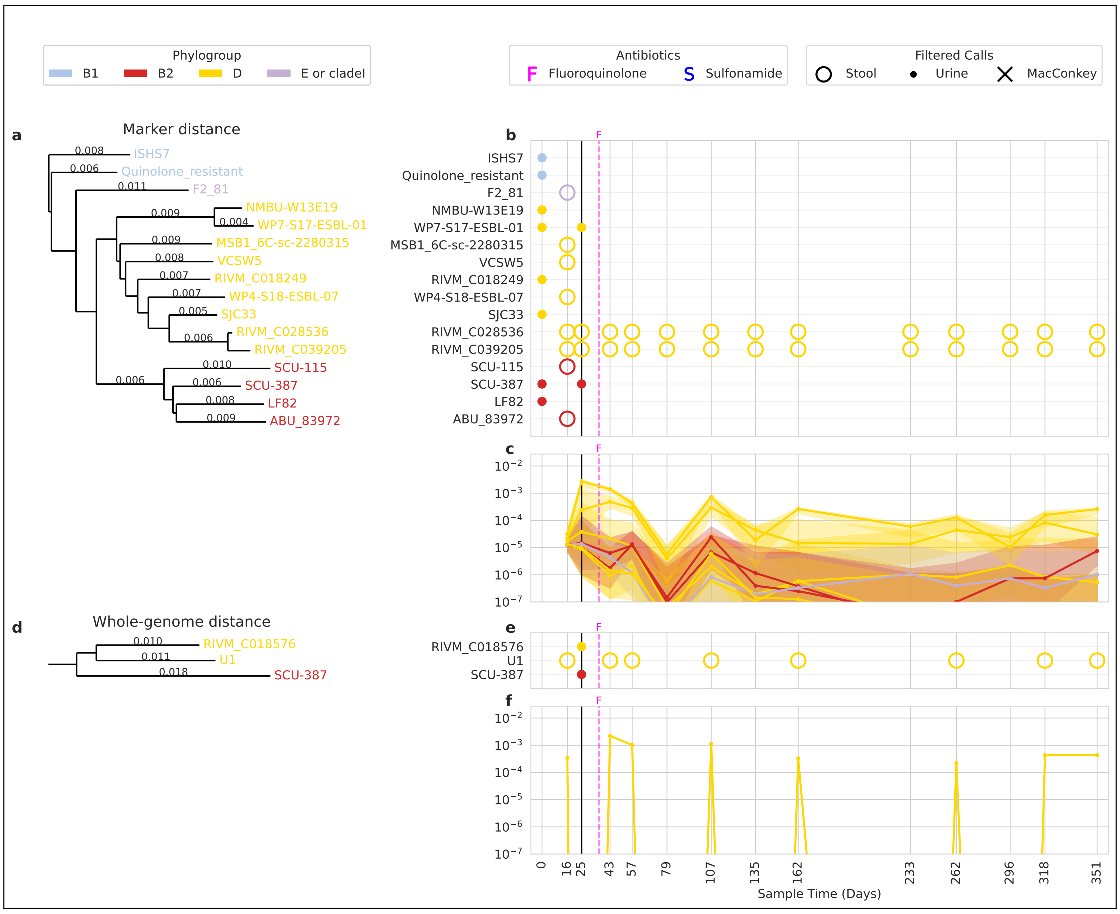


Figure U15: Participant UMB15

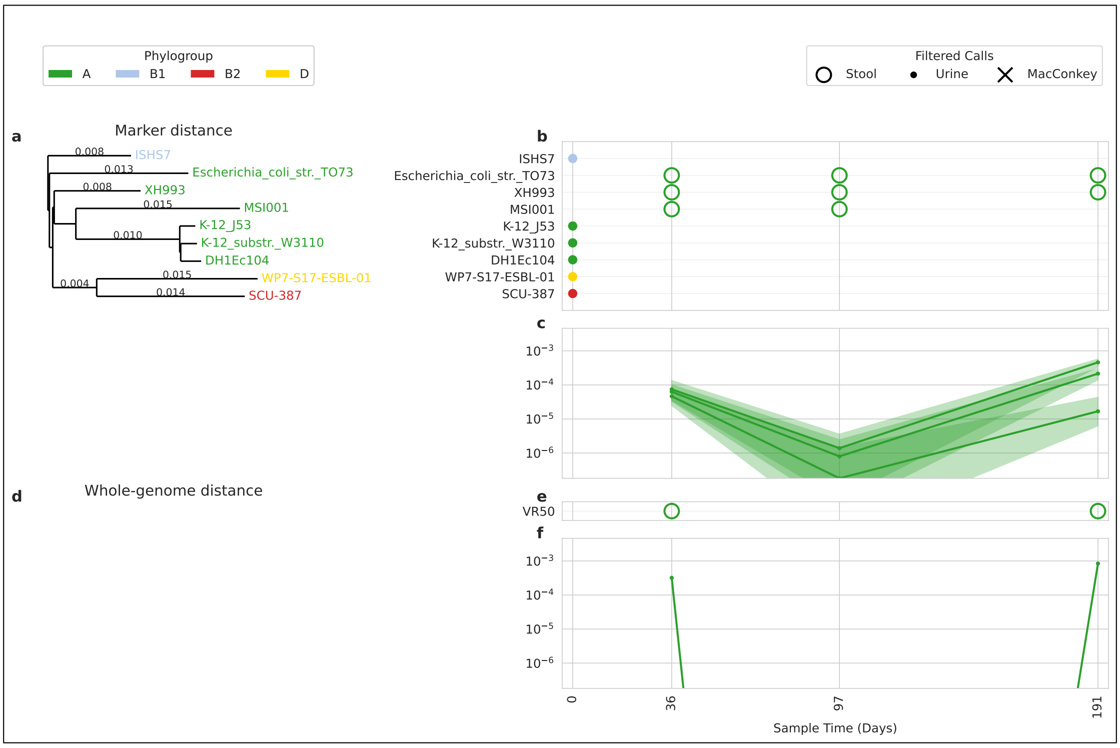


Figure U16: Participant UMB16

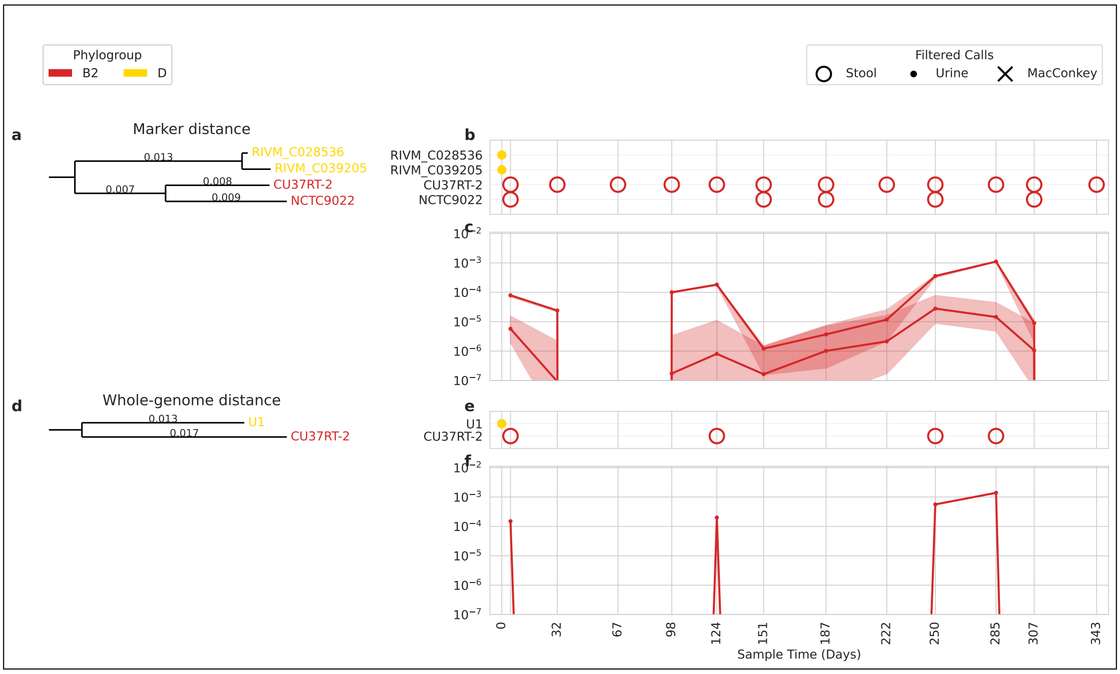


Figure U17: Participant UMB17

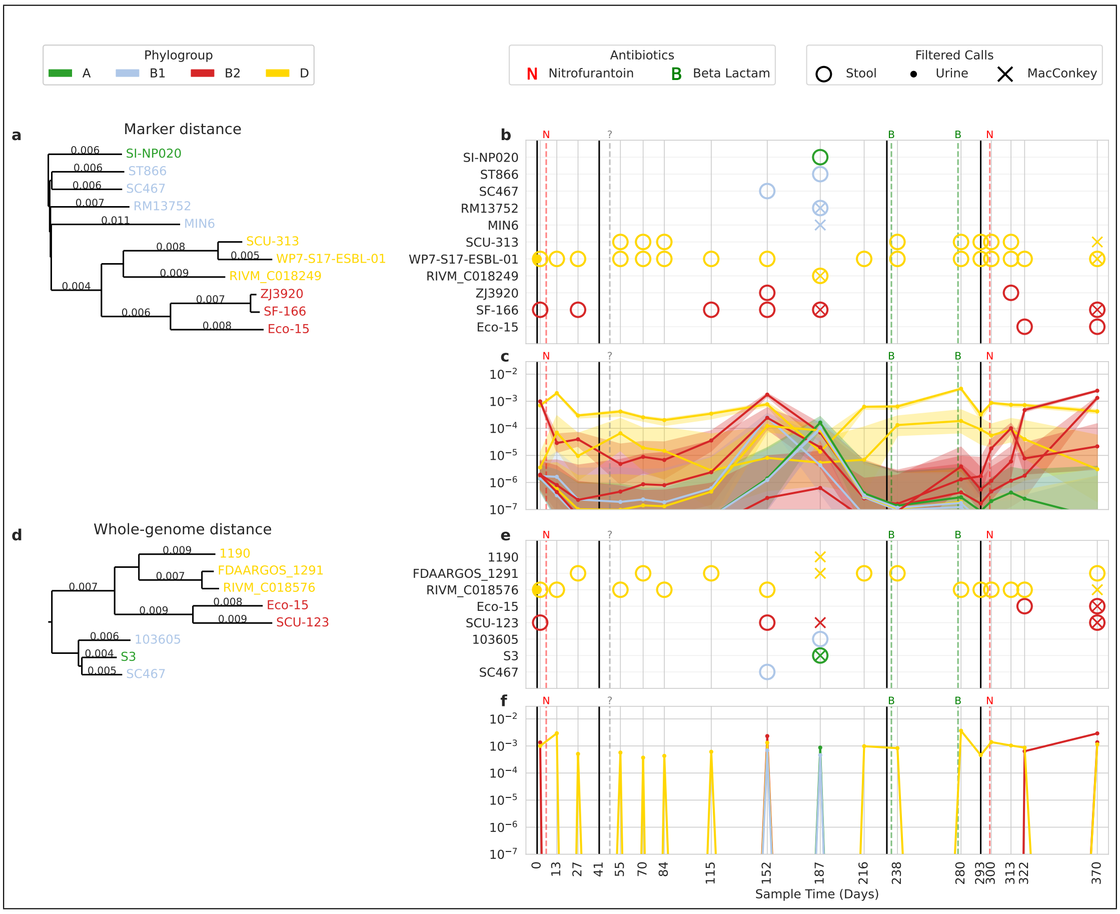


Figure U18: Participant UMB18

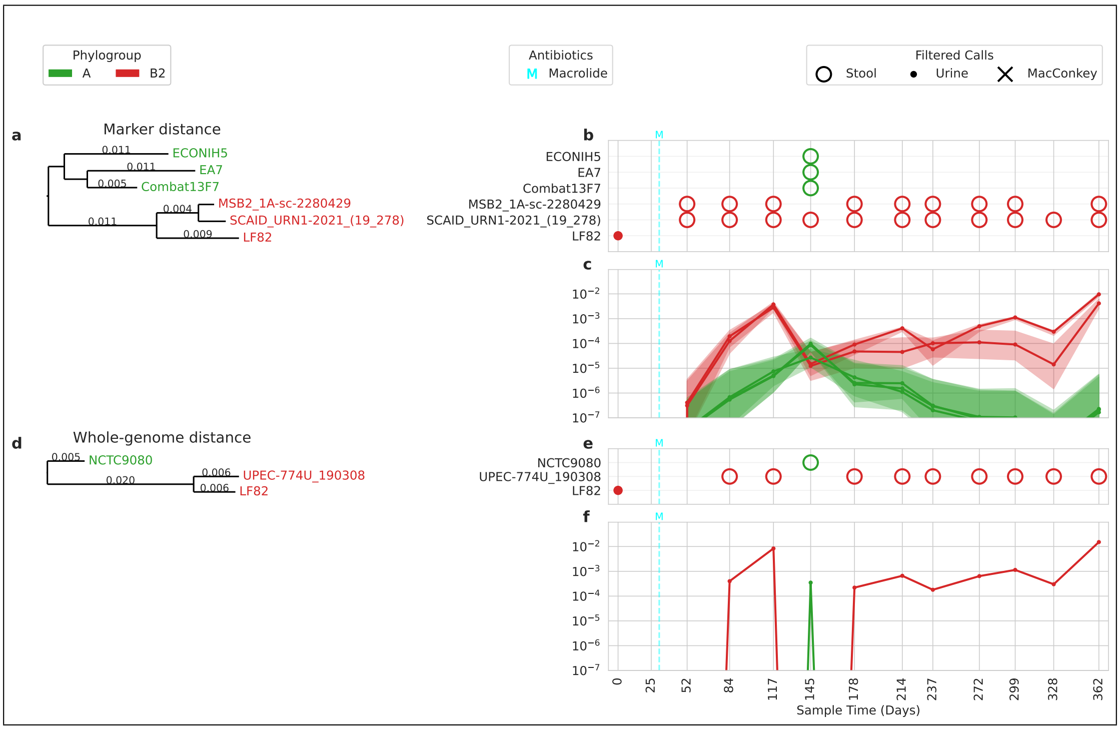


Figure U19: Participant UMB19

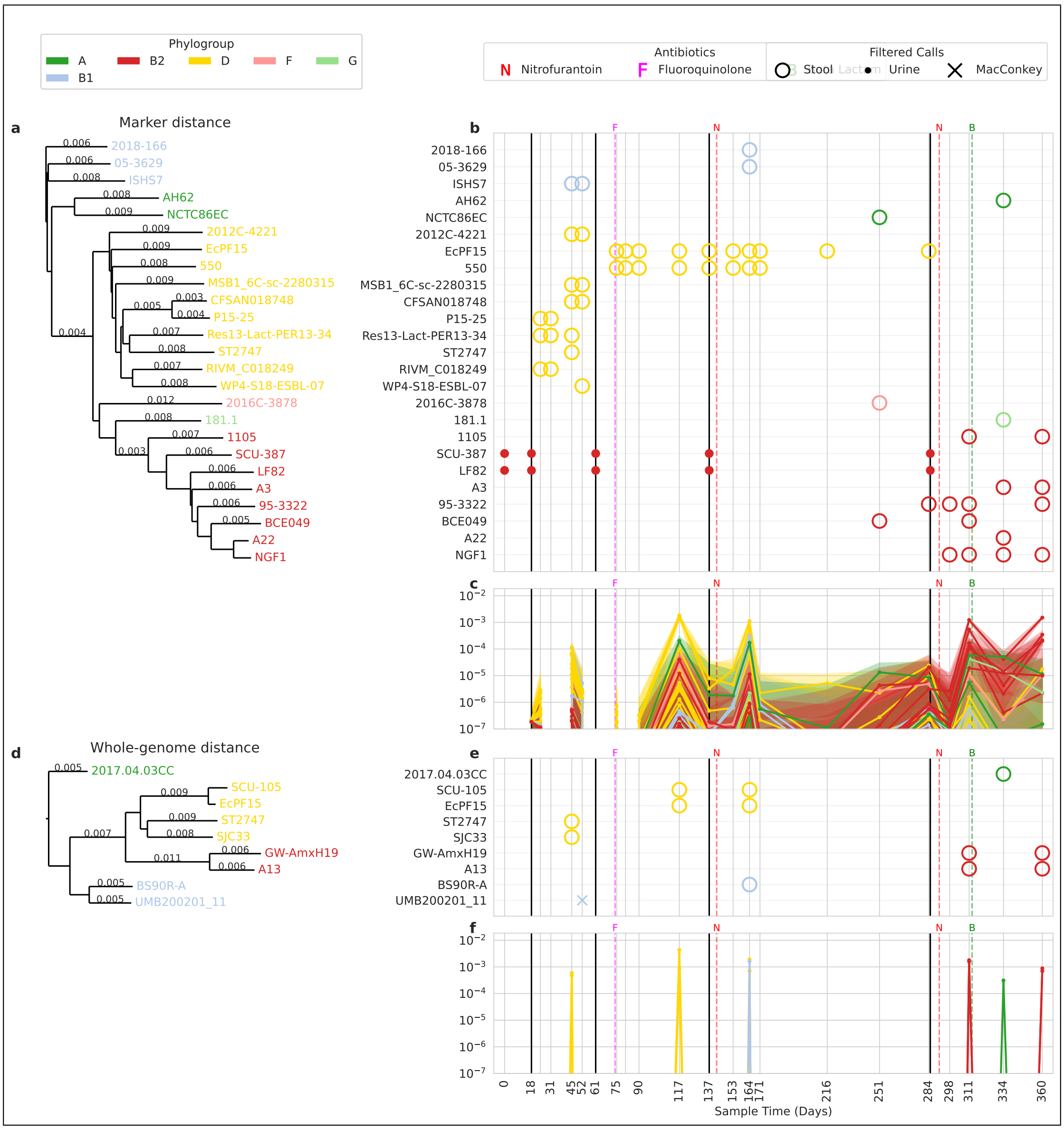


Figure U20: Participant UMB20

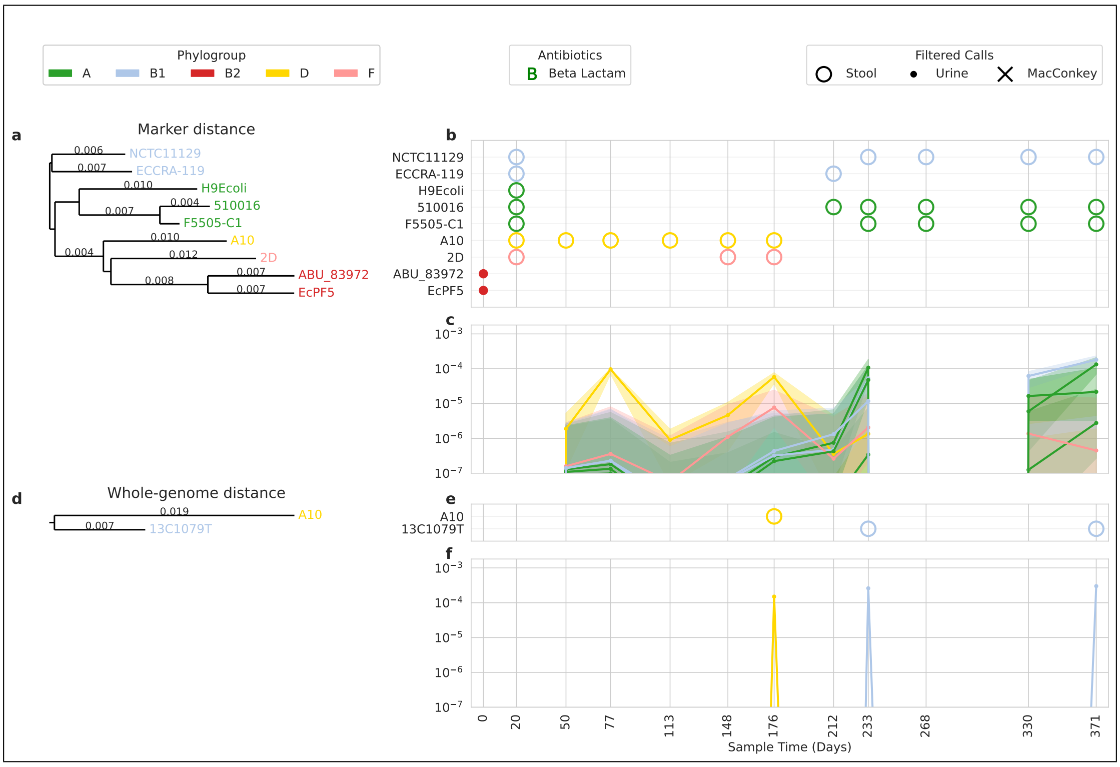


Figure U21: Participant UMB21

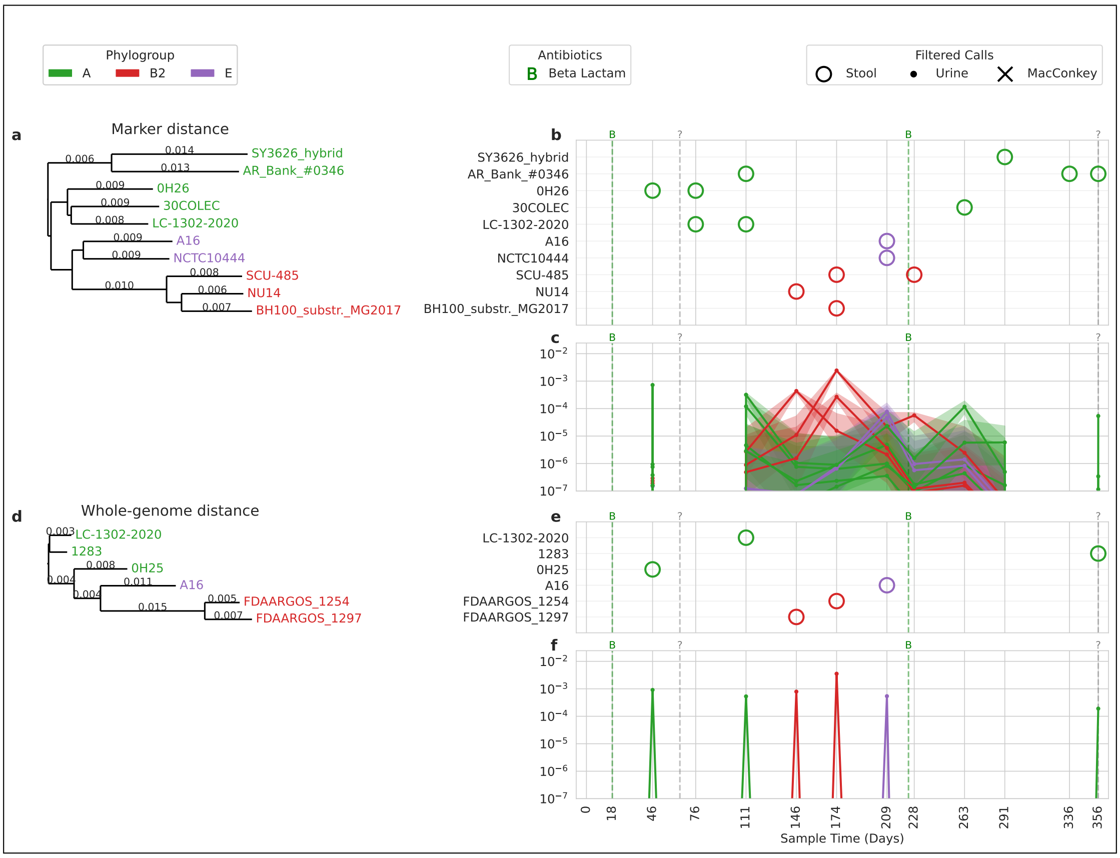


Figure U22: Participant UMB22

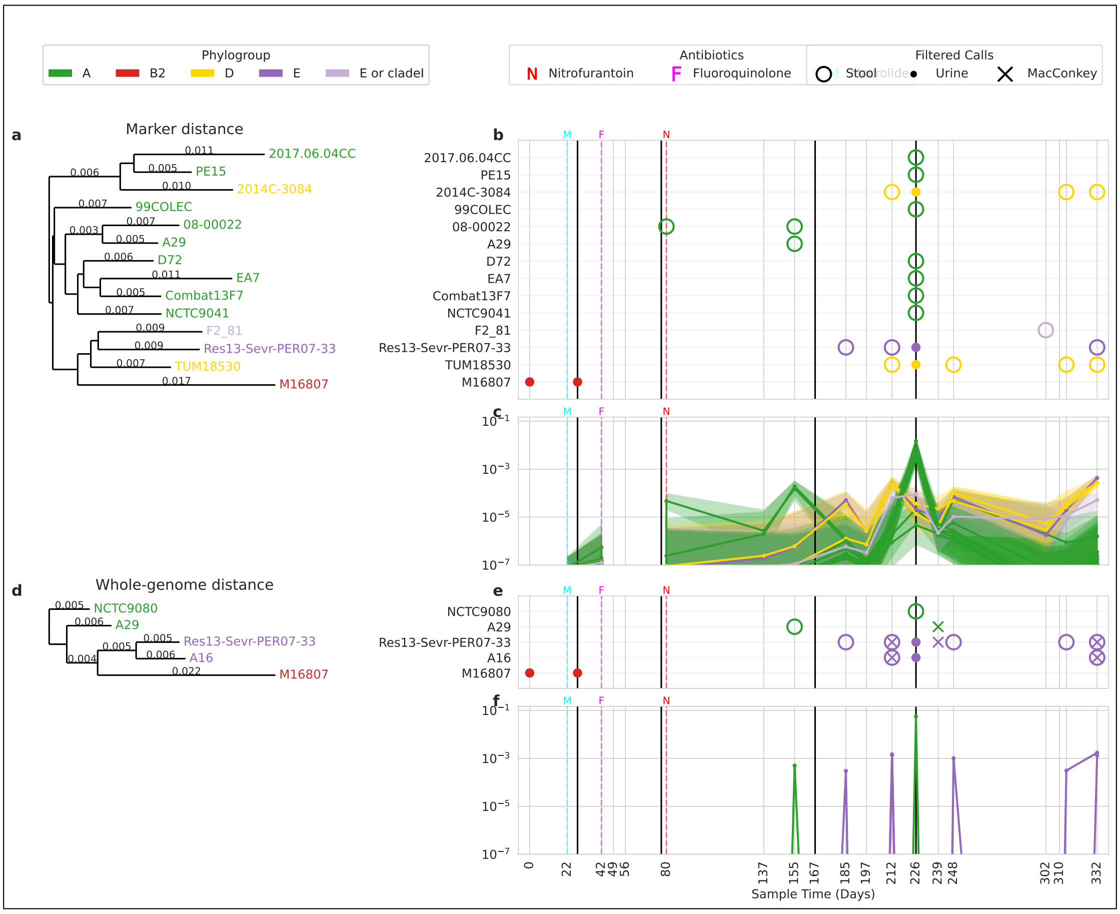


Figure U23: Participant UMB23

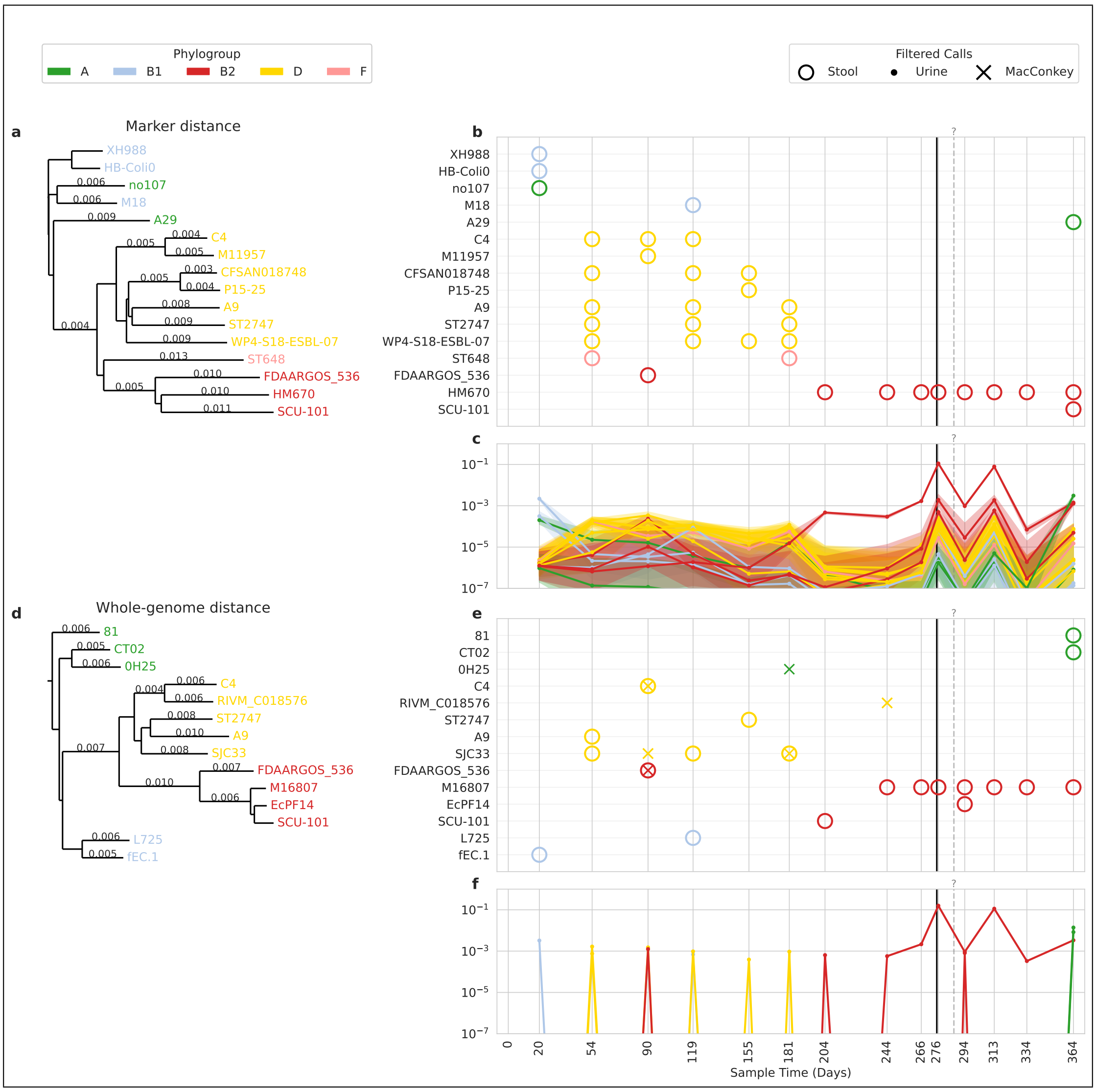


Figure U24: Participant UMB24

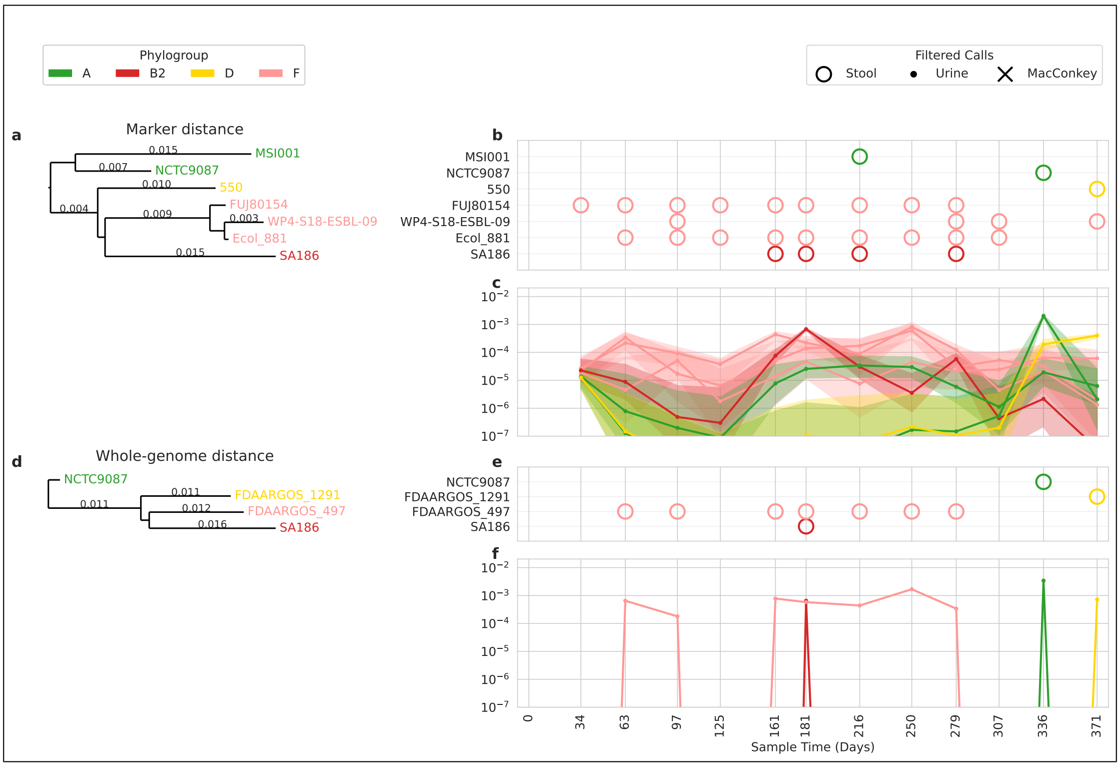


Figure U25: Participant UMB25

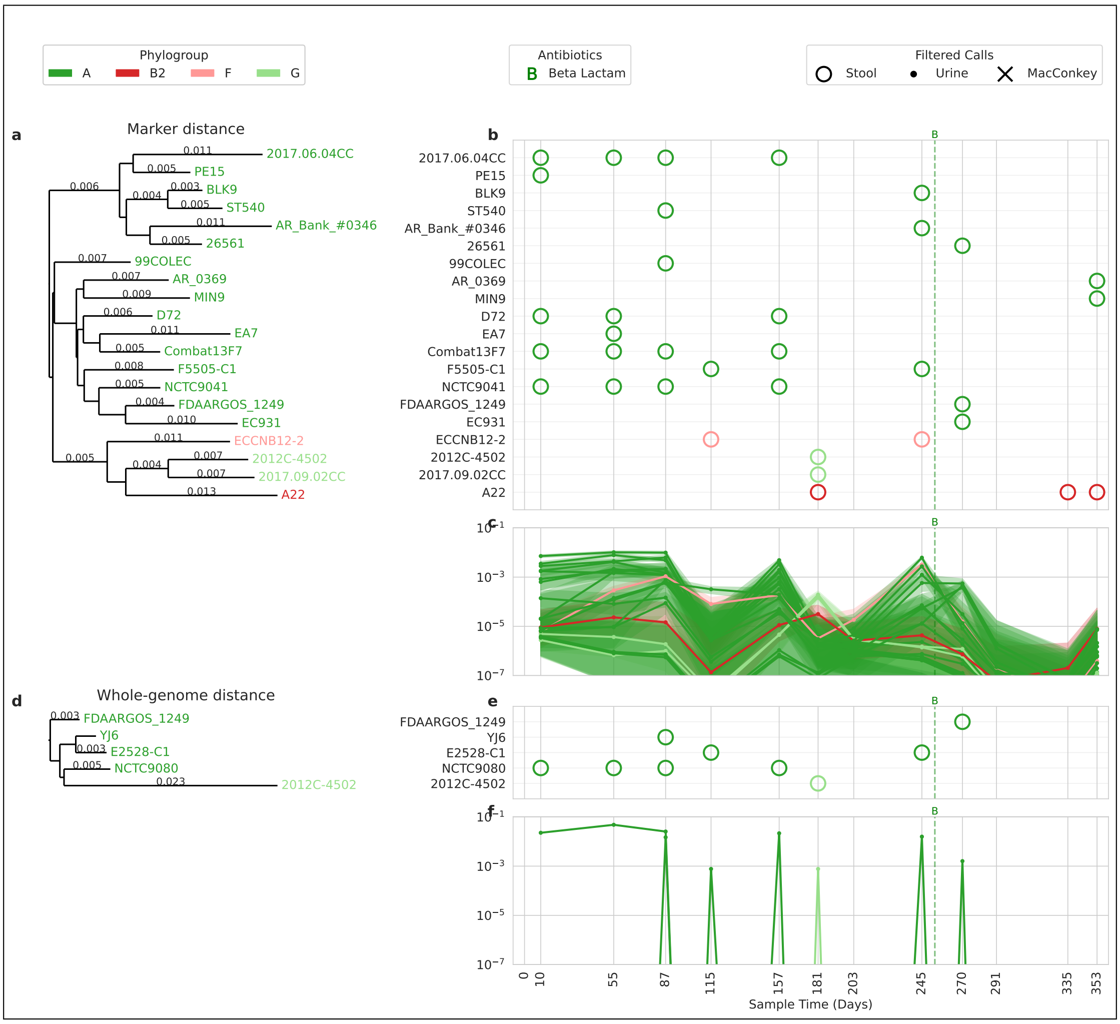


Figure U26: Participant UMB26

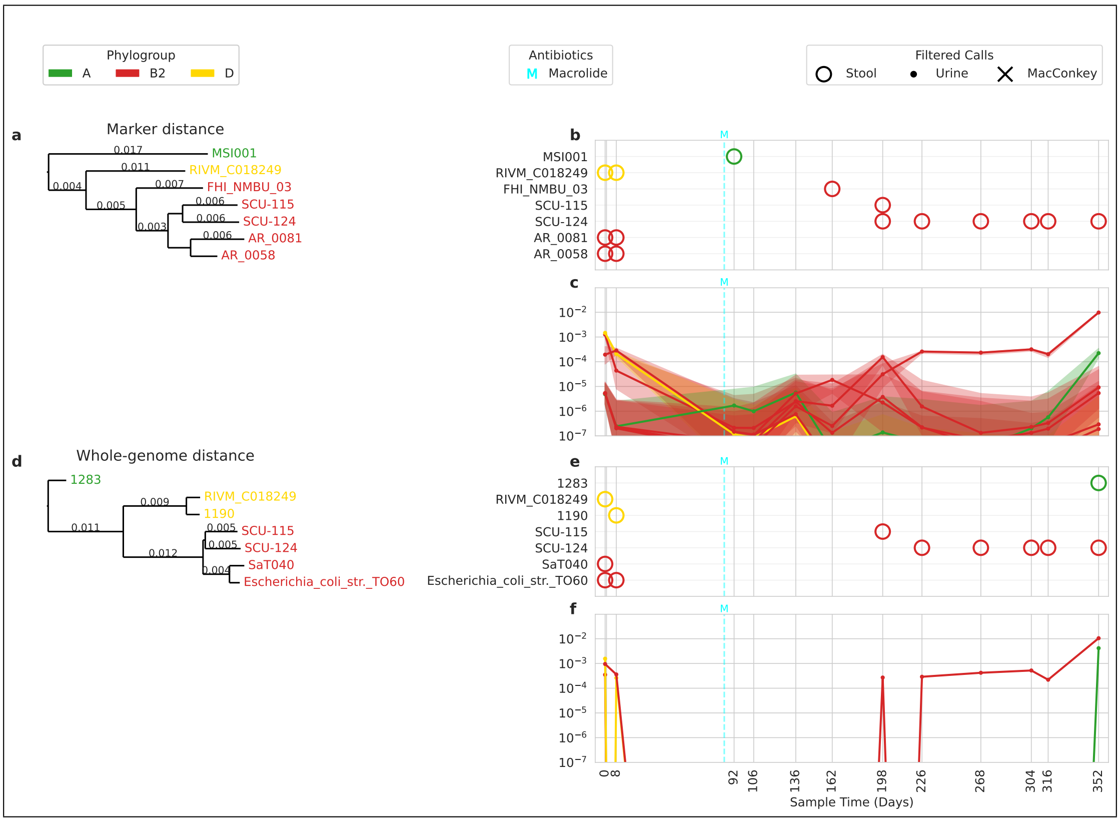


Figure U27: Participant UMB27

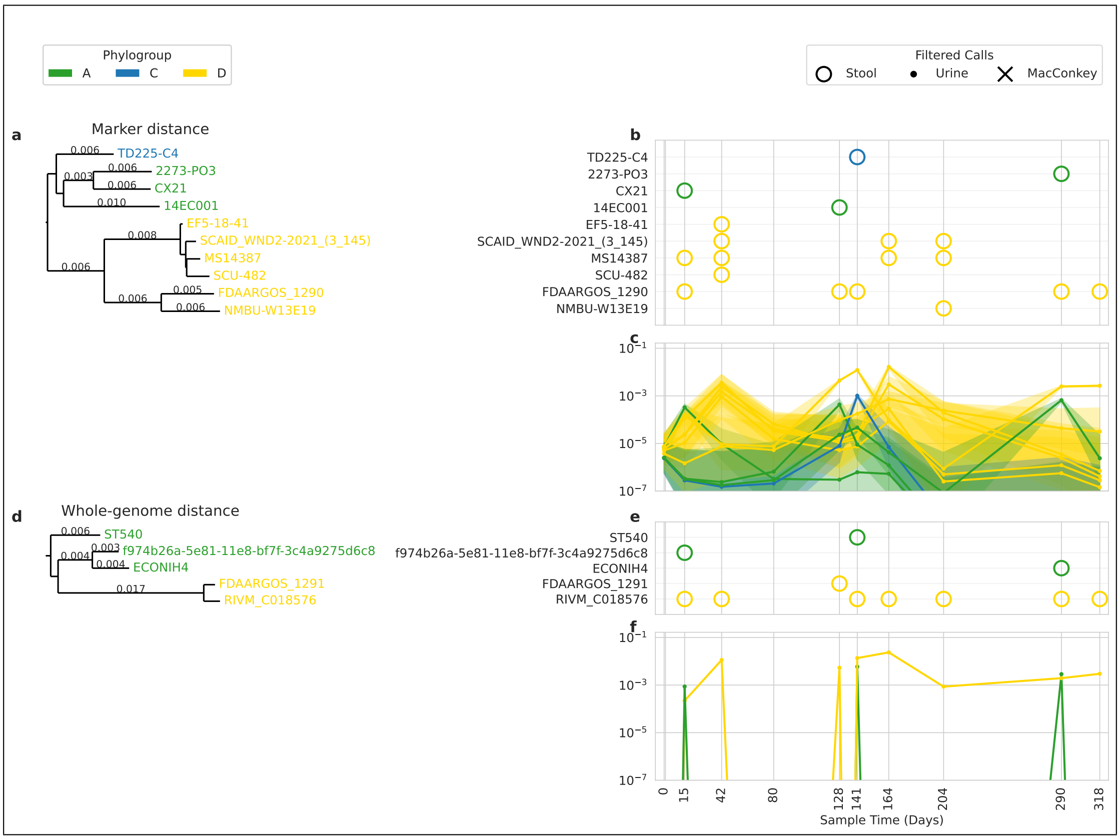


Figure U28: Participant UMB28

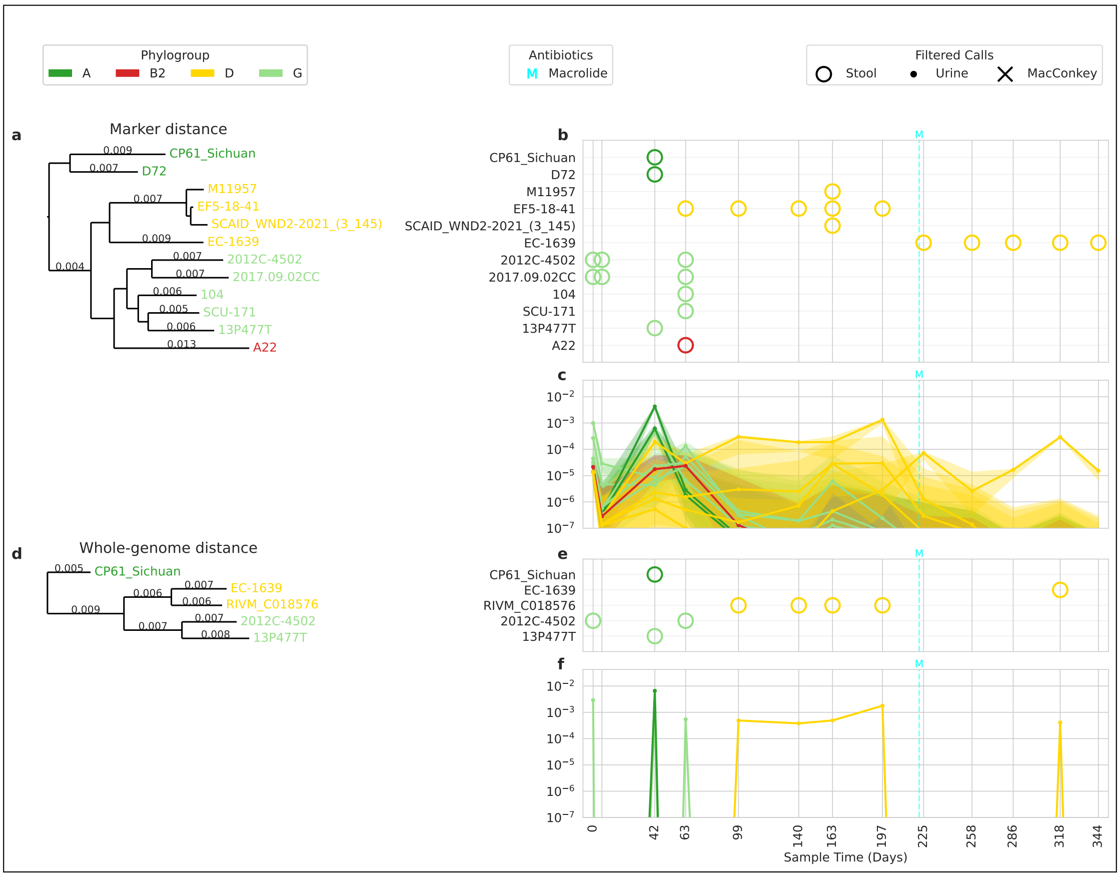


Figure U29: Participant UMB29

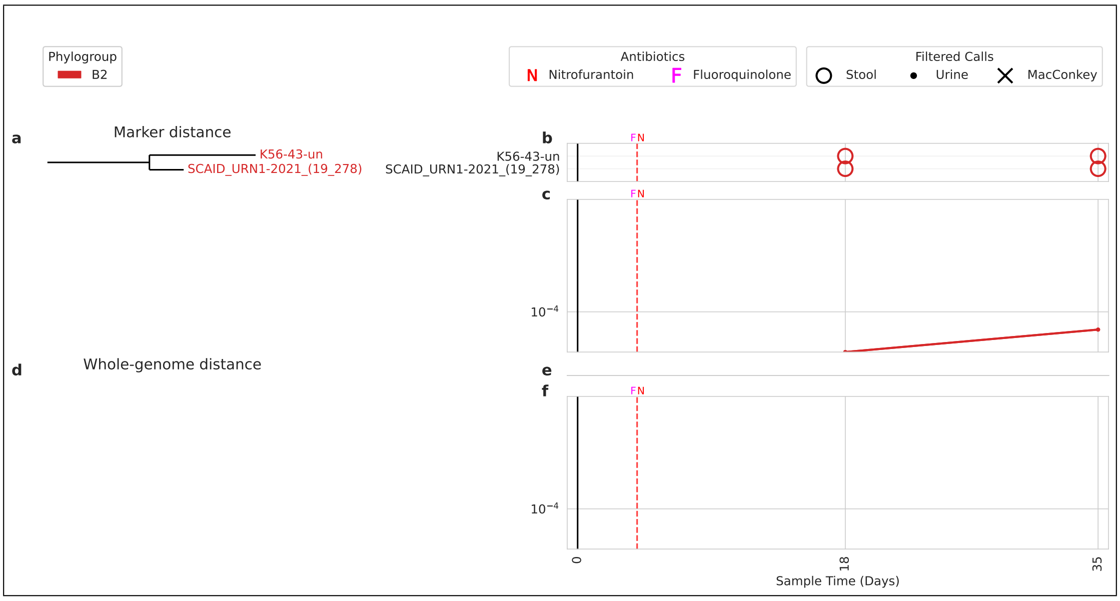


Figure U30: Participant UMB30

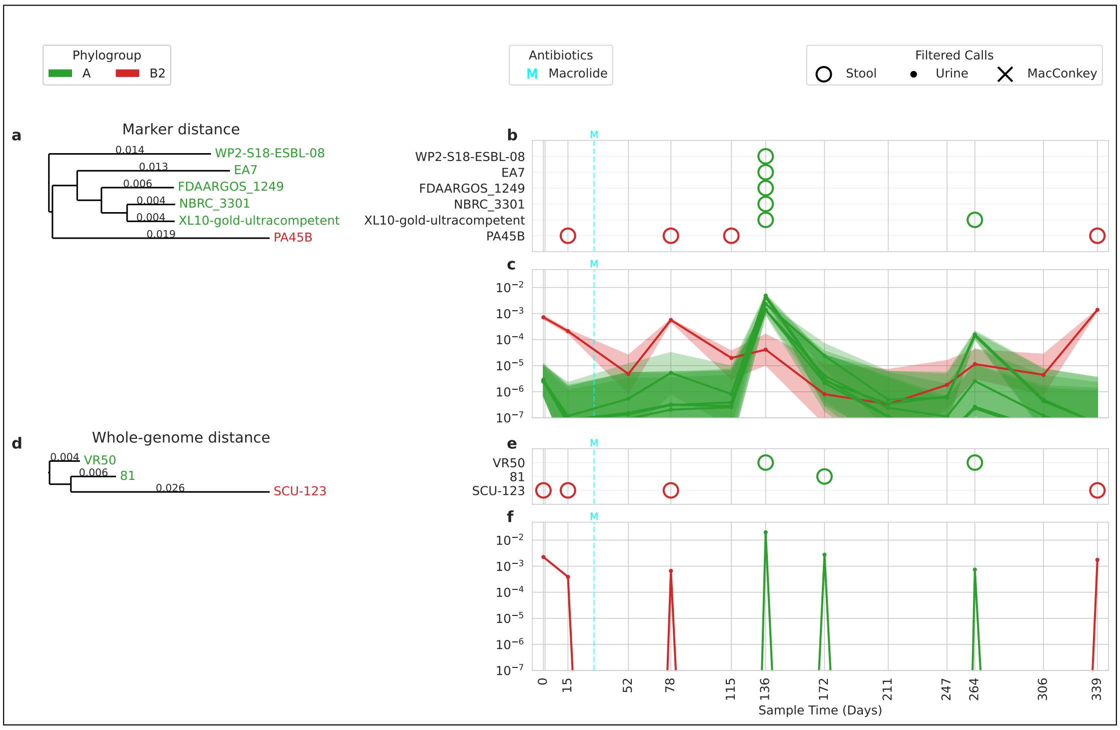


Figure U31: Participant UMB31

### 2 ELMC – timeseries plots per infant

Here, we provide all time-series plots for the ELMC cohort infant analysis. In the style of the main text (Figure 5, panels a1-a3 and b1-b3), we plot the results using ChronoStrain and using mGEMS after each respective method's filtering steps. For details on the methods and the post-hoc filters used, refer to the Methods section.

As a reminder: just like the UMB illustrations, each $y$-axis is the *overall* relative abundance, and not the *database-normalized* abundance which is method-specific. The red lines indicates MetaPhlAn4 *species*-level abundance estimates, while the blue lines indicate Bracken species-level estimates (the triangle pointers only serve to improve visibility). The markers (circles or X) are drawn for a trajectory if the corresponding cluster passed the method's filter at that timepoint. An O indicates that the strain cluster contains an isolate from the same timepoint and from the same infant, whereas an X indicates no isolate (but it still passed the abundance/quality filter).

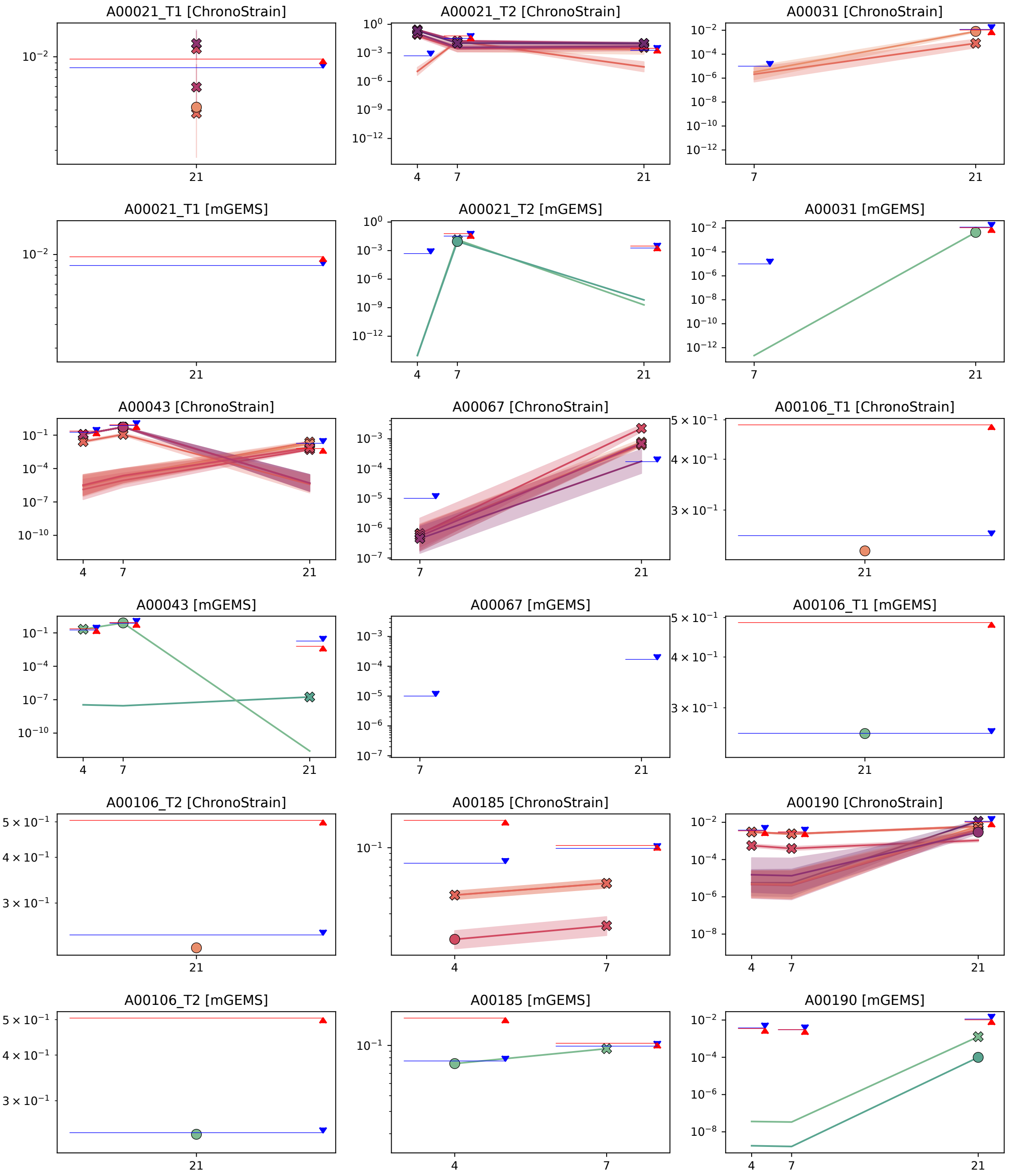


Figure E1: ELMC results (part 1 of 21)

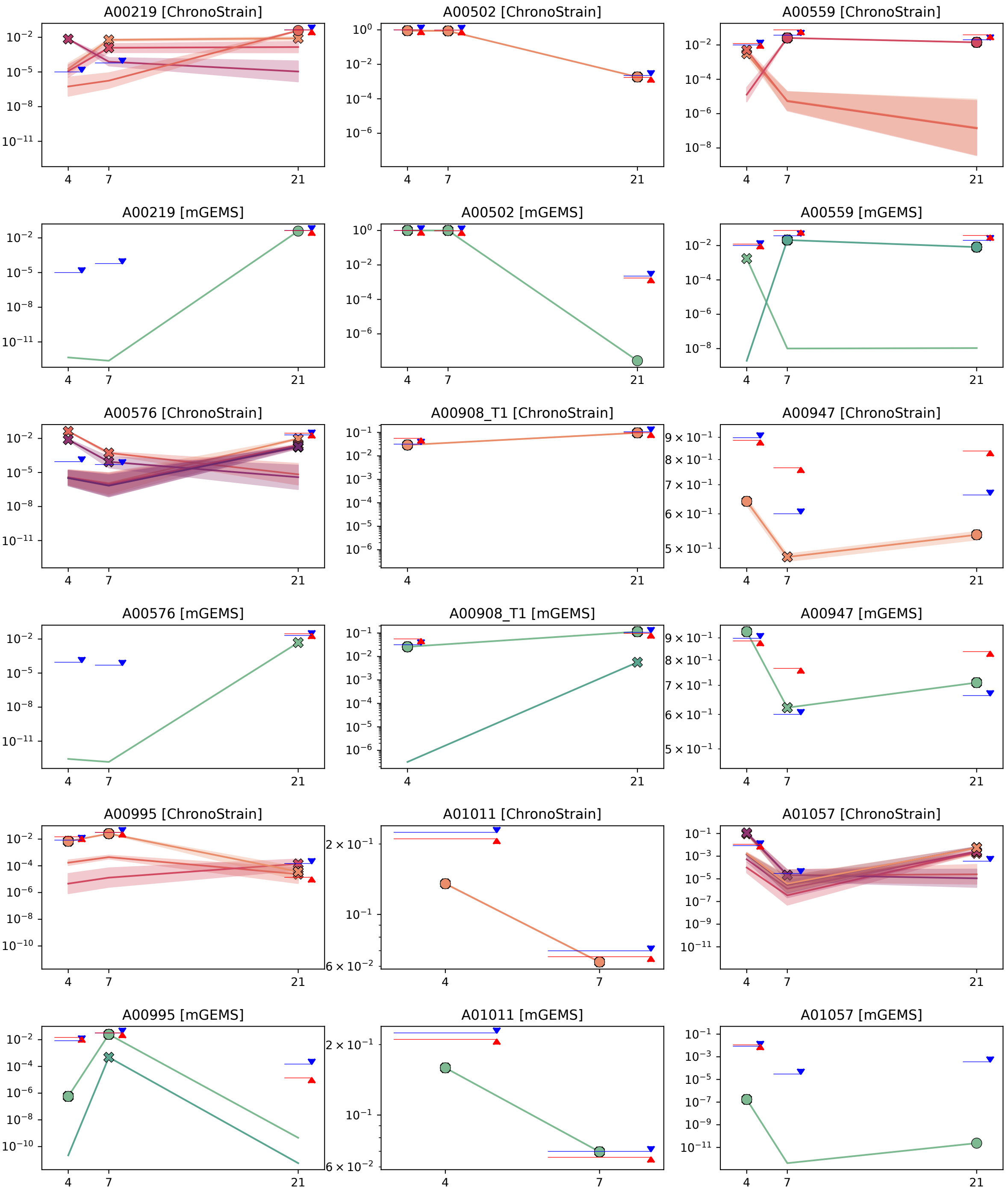


Figure E2: ELMC results (part 2 of 21)

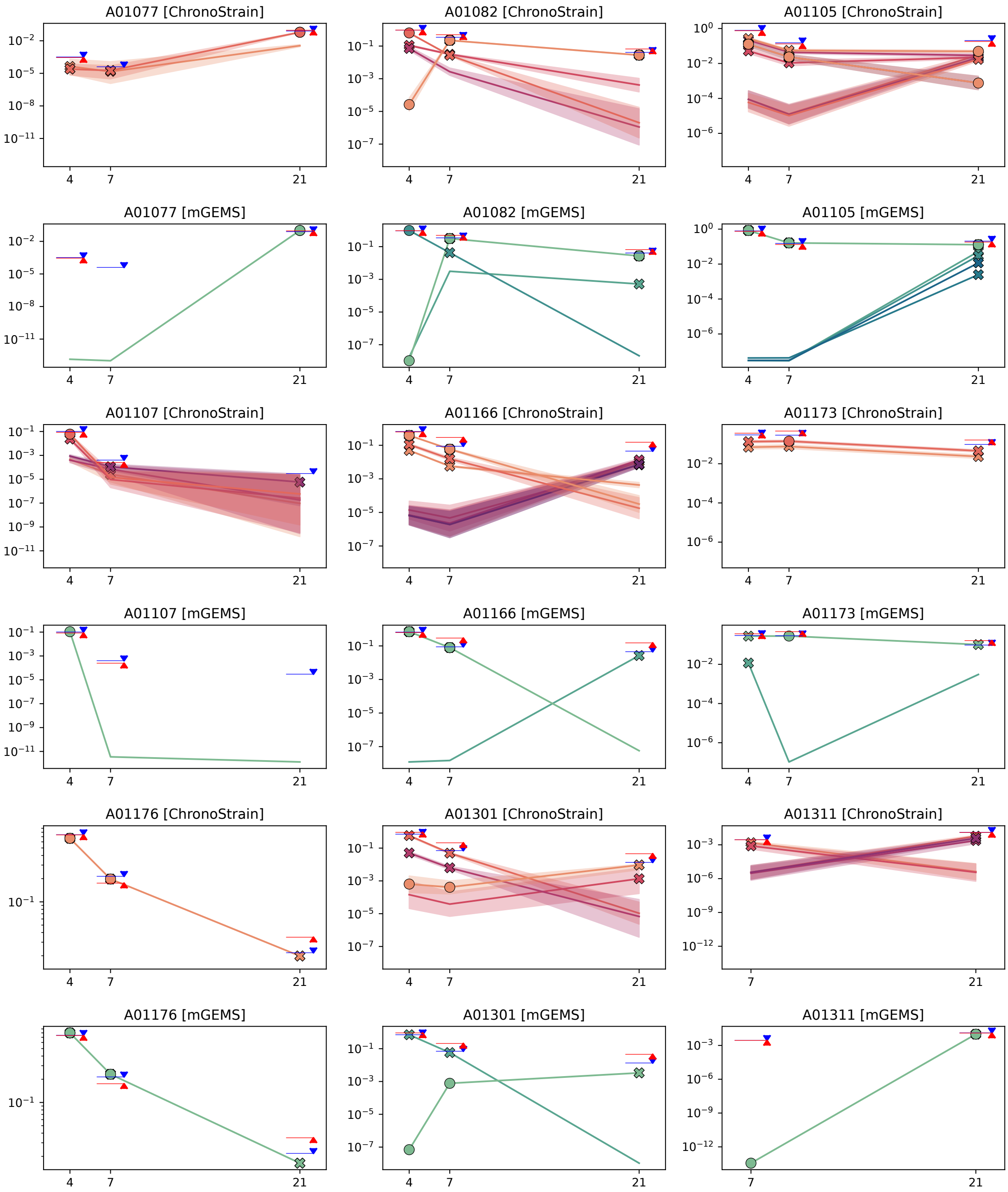


Figure E3: ELMC results (part 3 of 21)

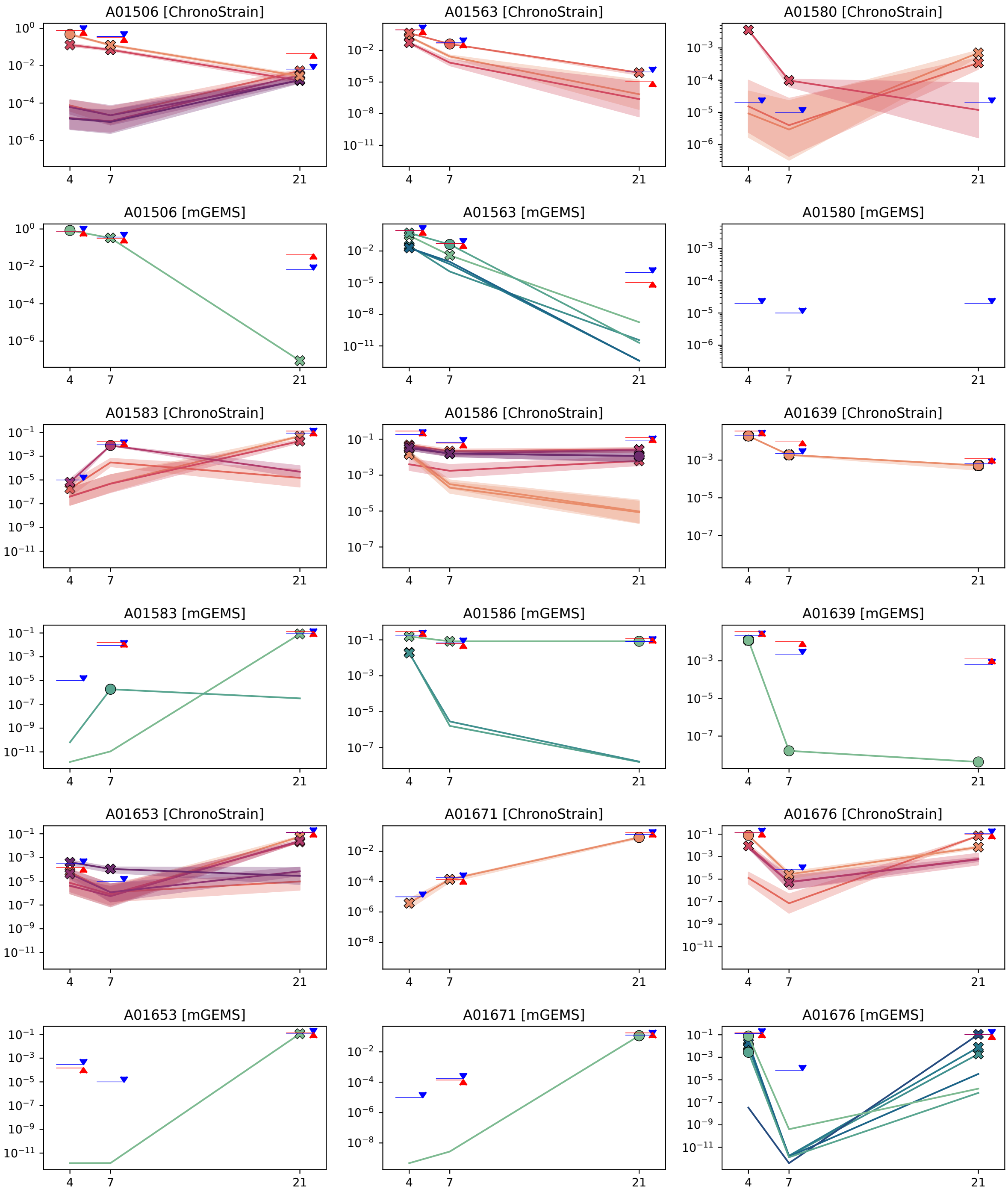


Figure E4: ELMC results (part 4 of 21)

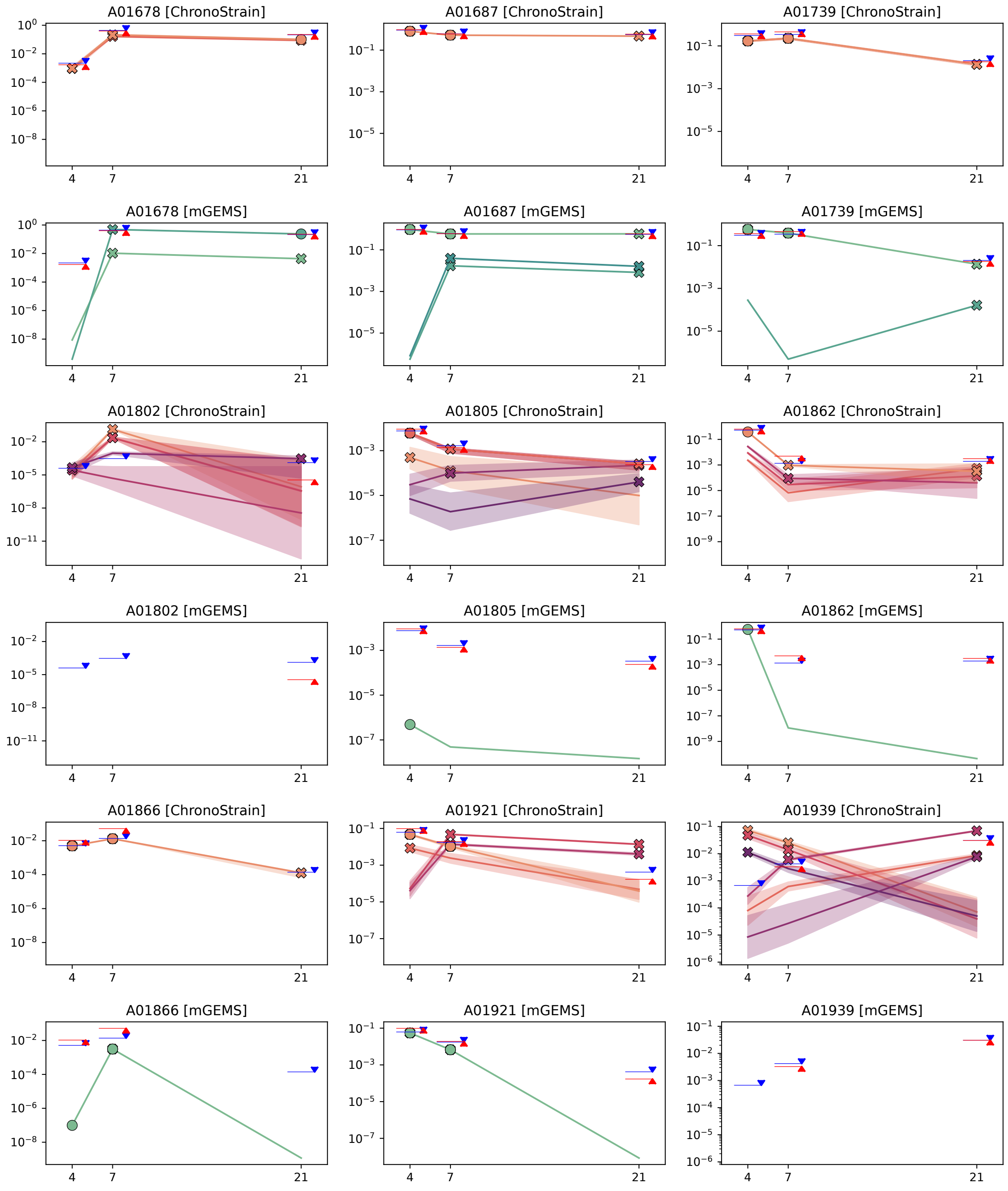


Figure E5: ELMC results (part 5 of 21)

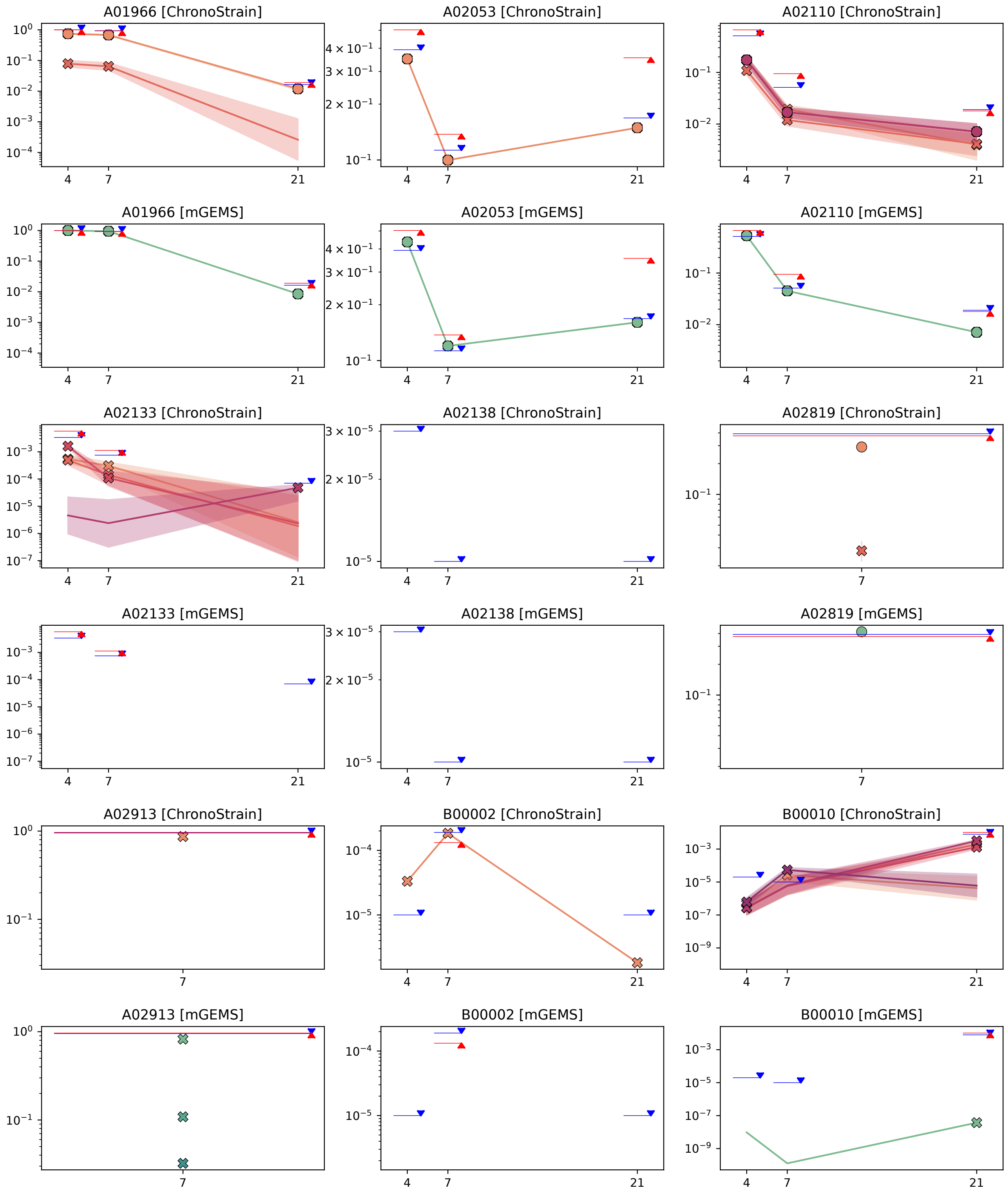


Figure E6: ELMC results (part 6 of 21)

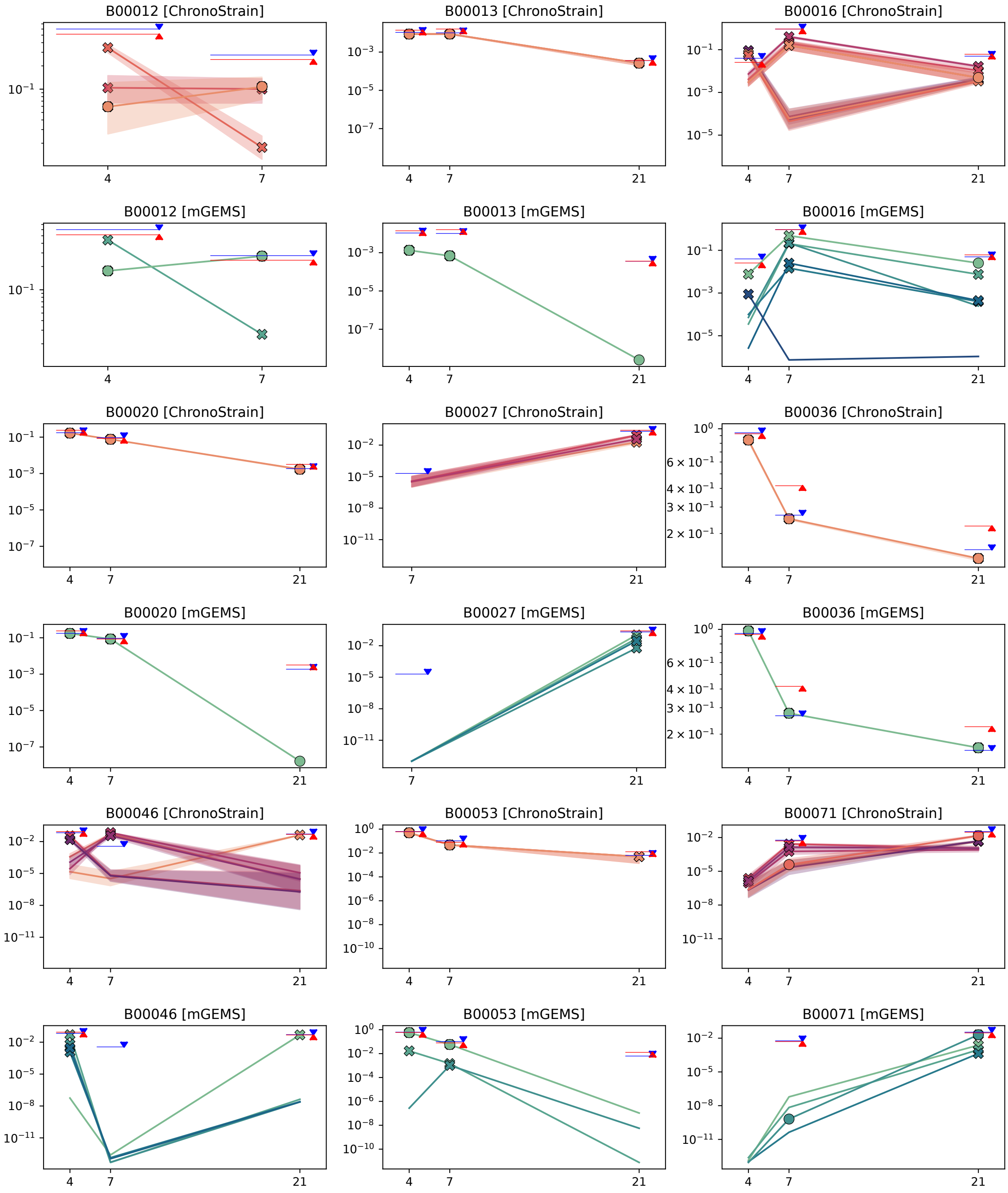


Figure E7: ELMC results (part 7 of 21)

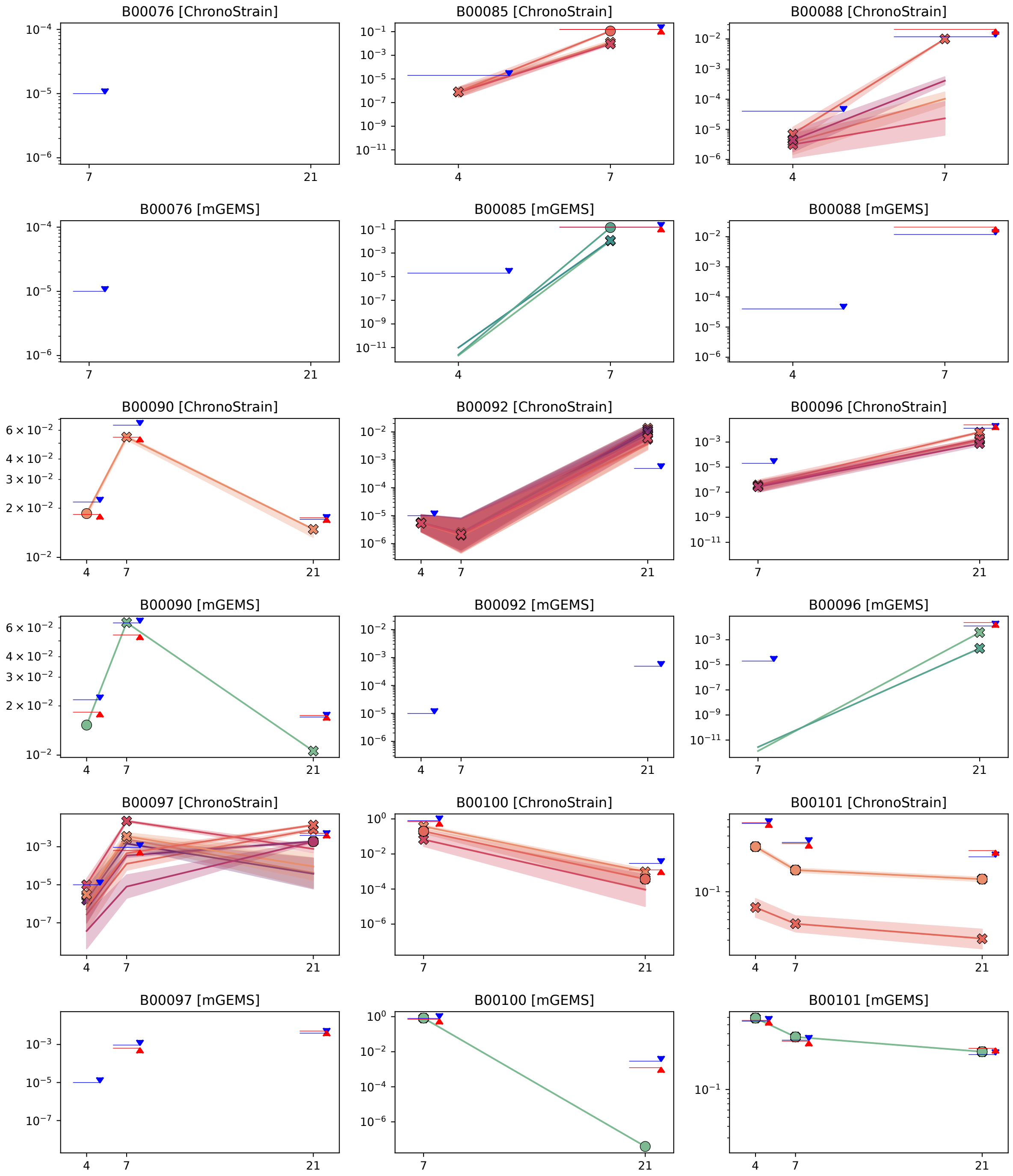


Figure E8: ELMC results (part 8 of 21)

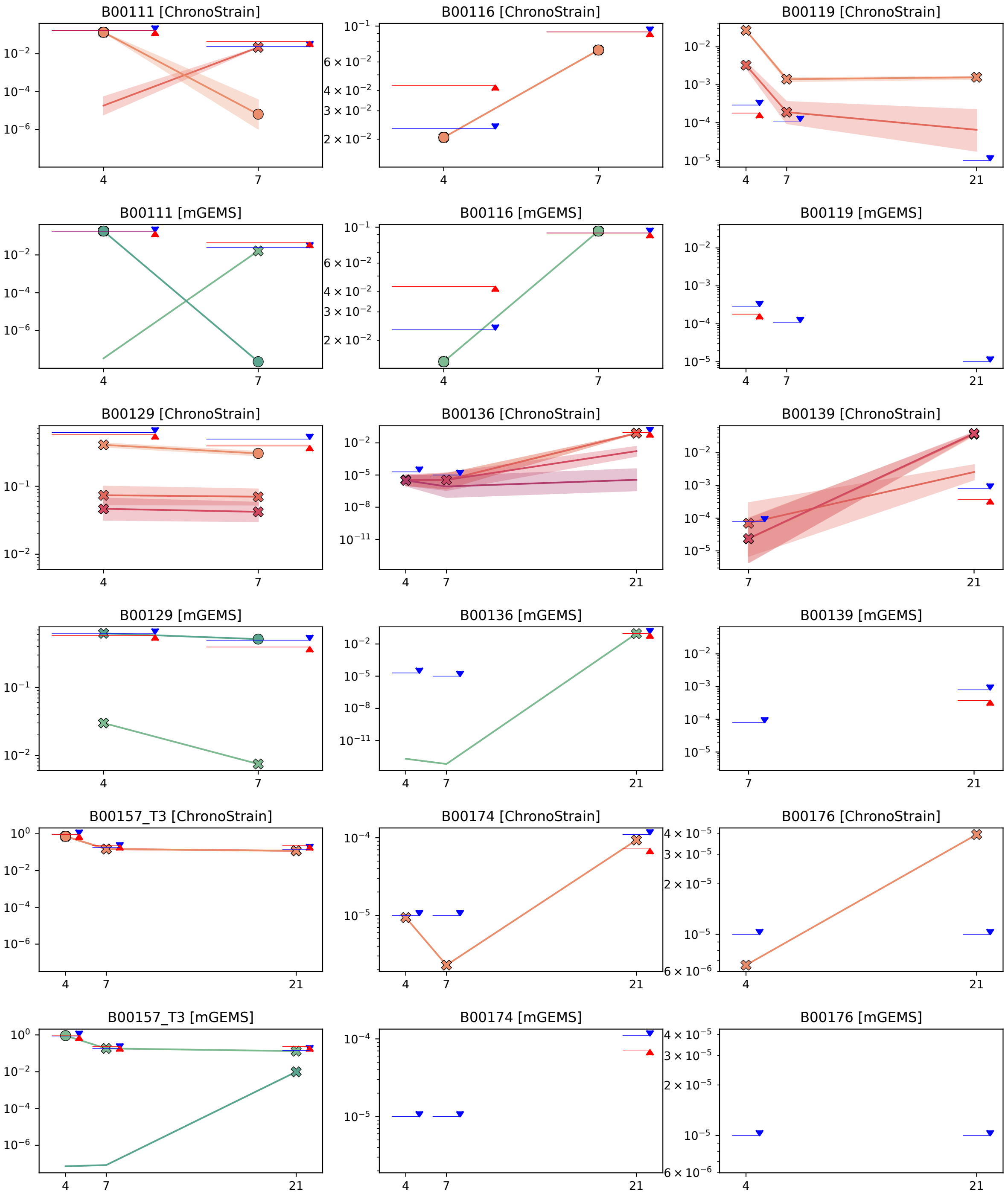


Figure E9: ELMC results (part 9 of 21)

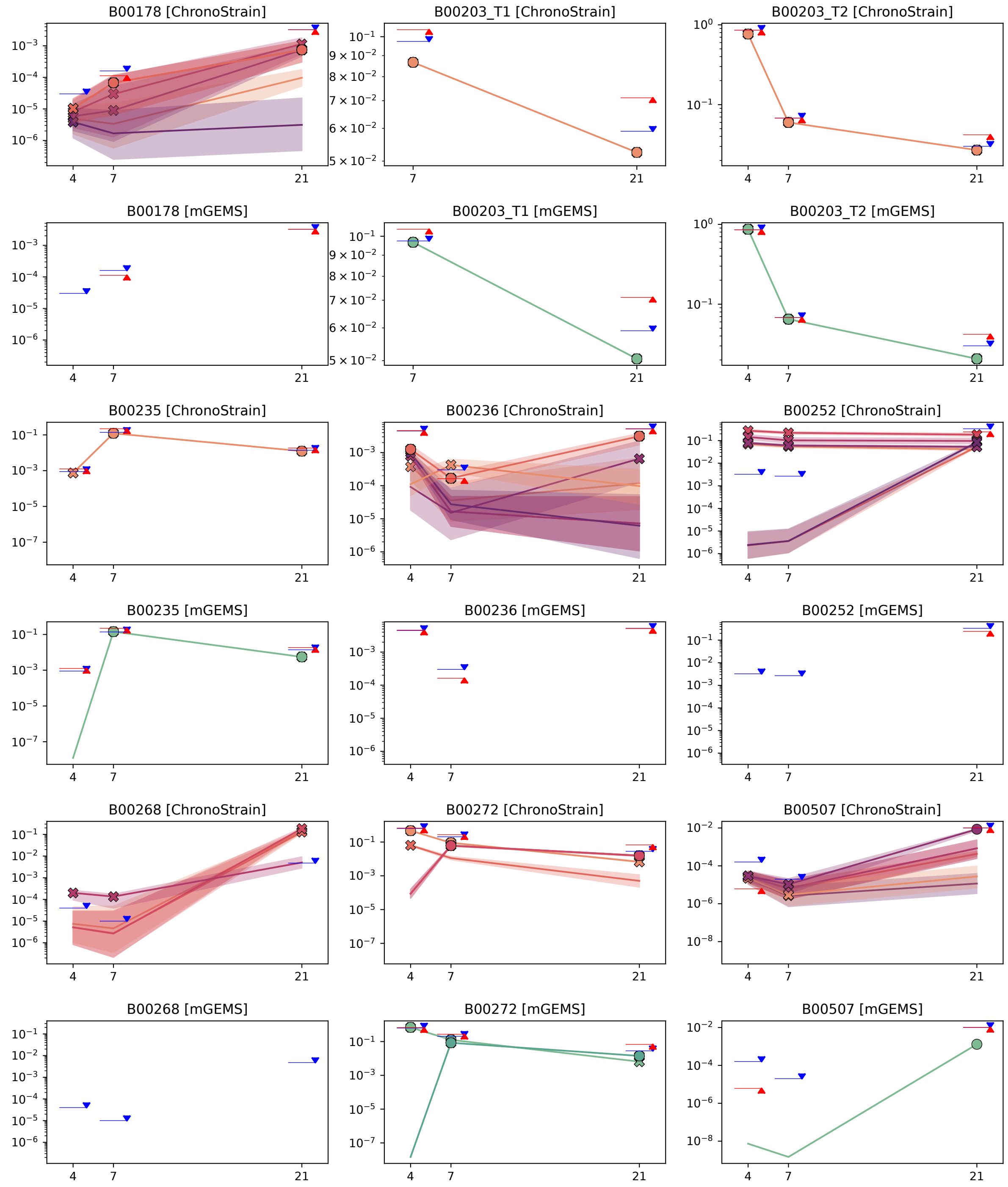


Figure E10: ELMC results (part 10 of 21)

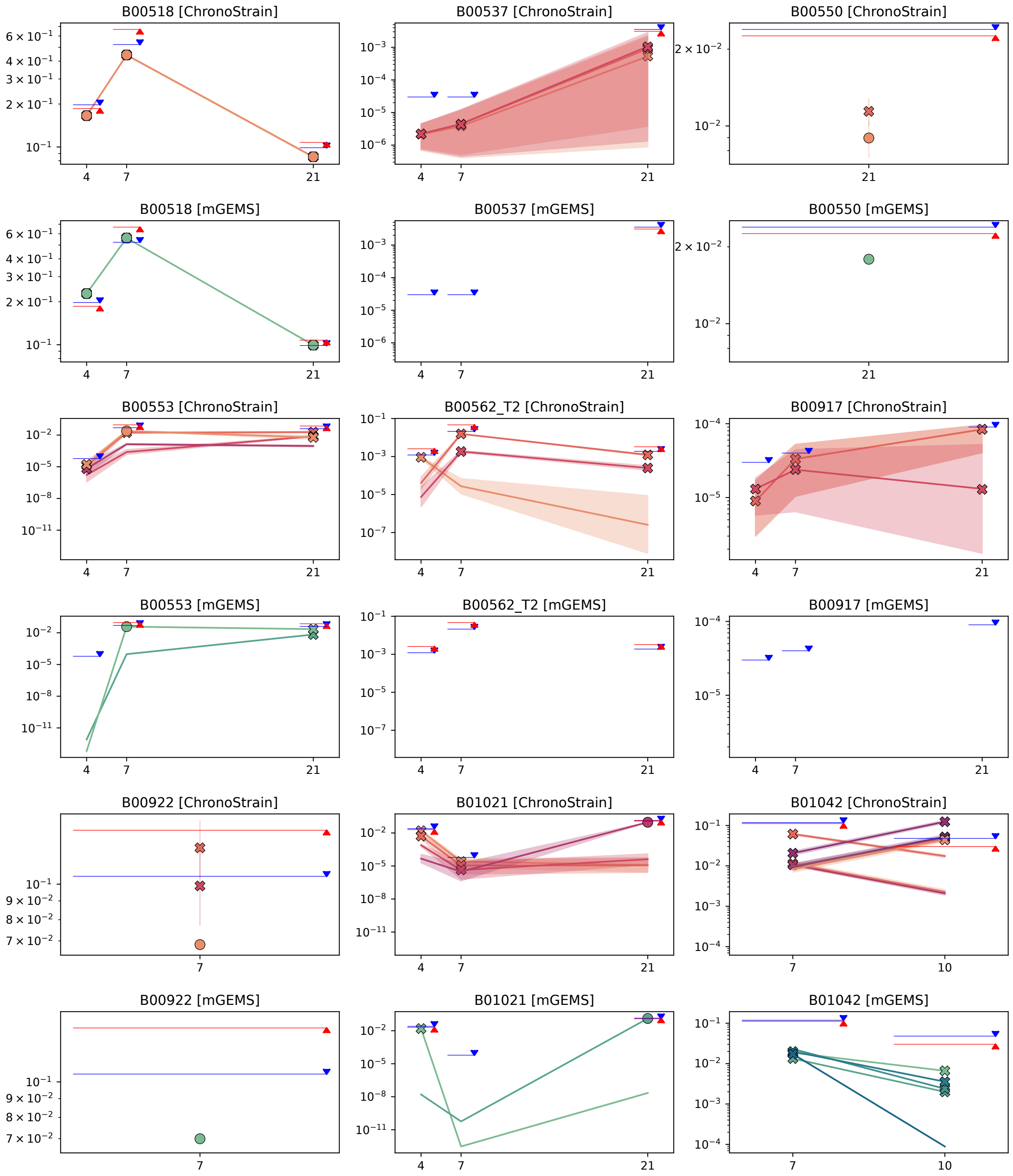


Figure E11: ELMC results (part 11 of 21)

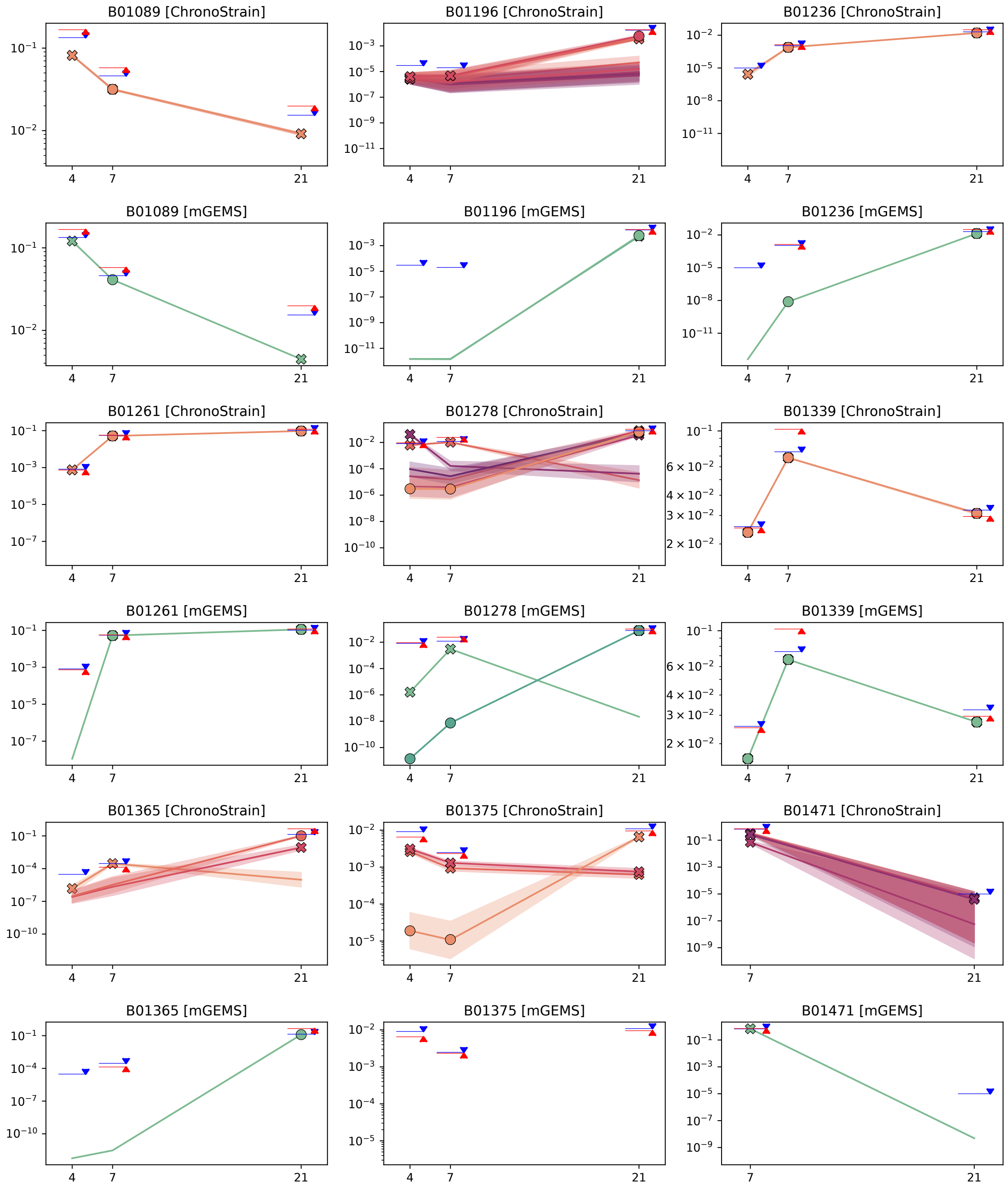


Figure E12: ELMC results (part 12 of 21)

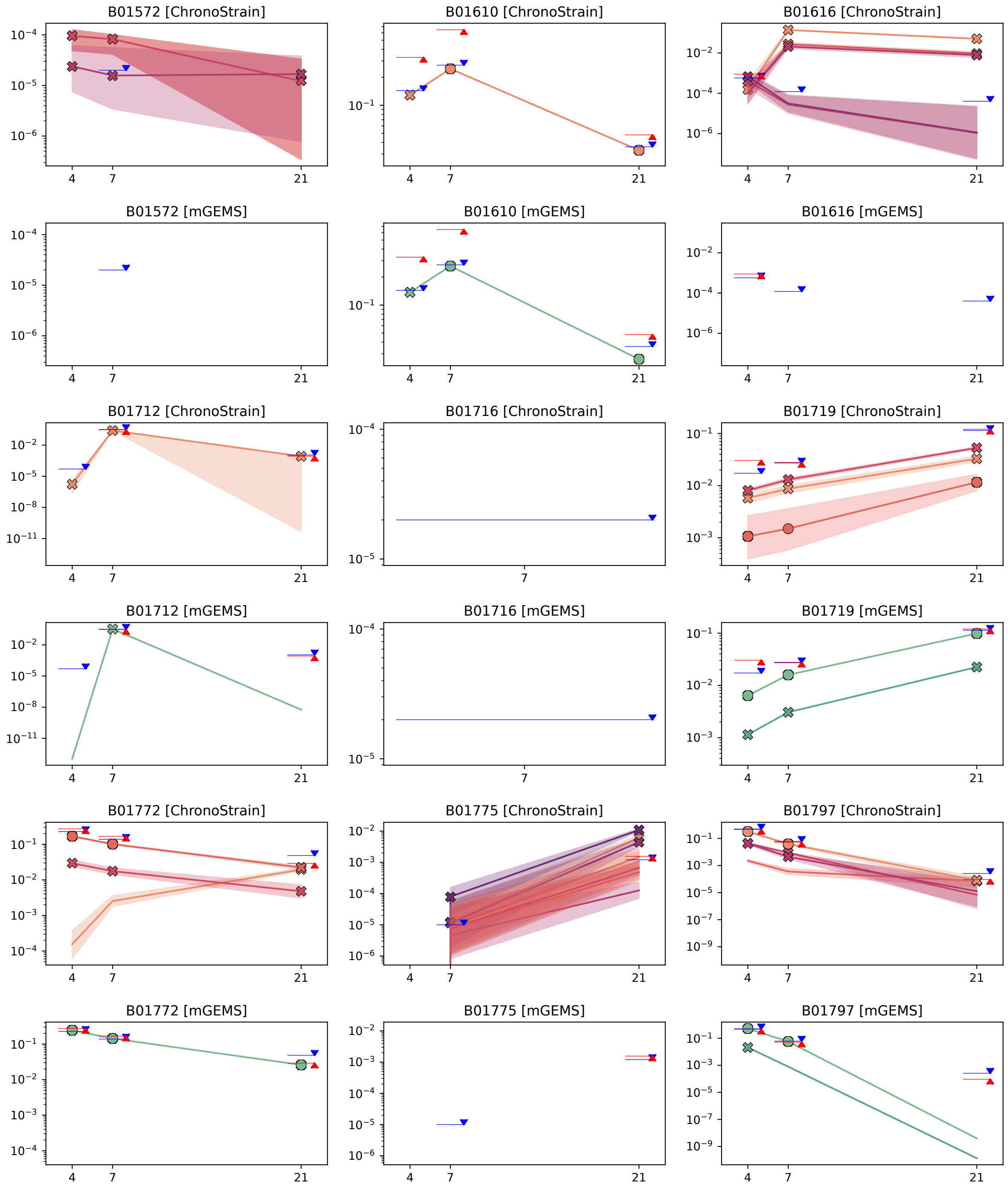


Figure E13: ELMC results (part 13 of 21)

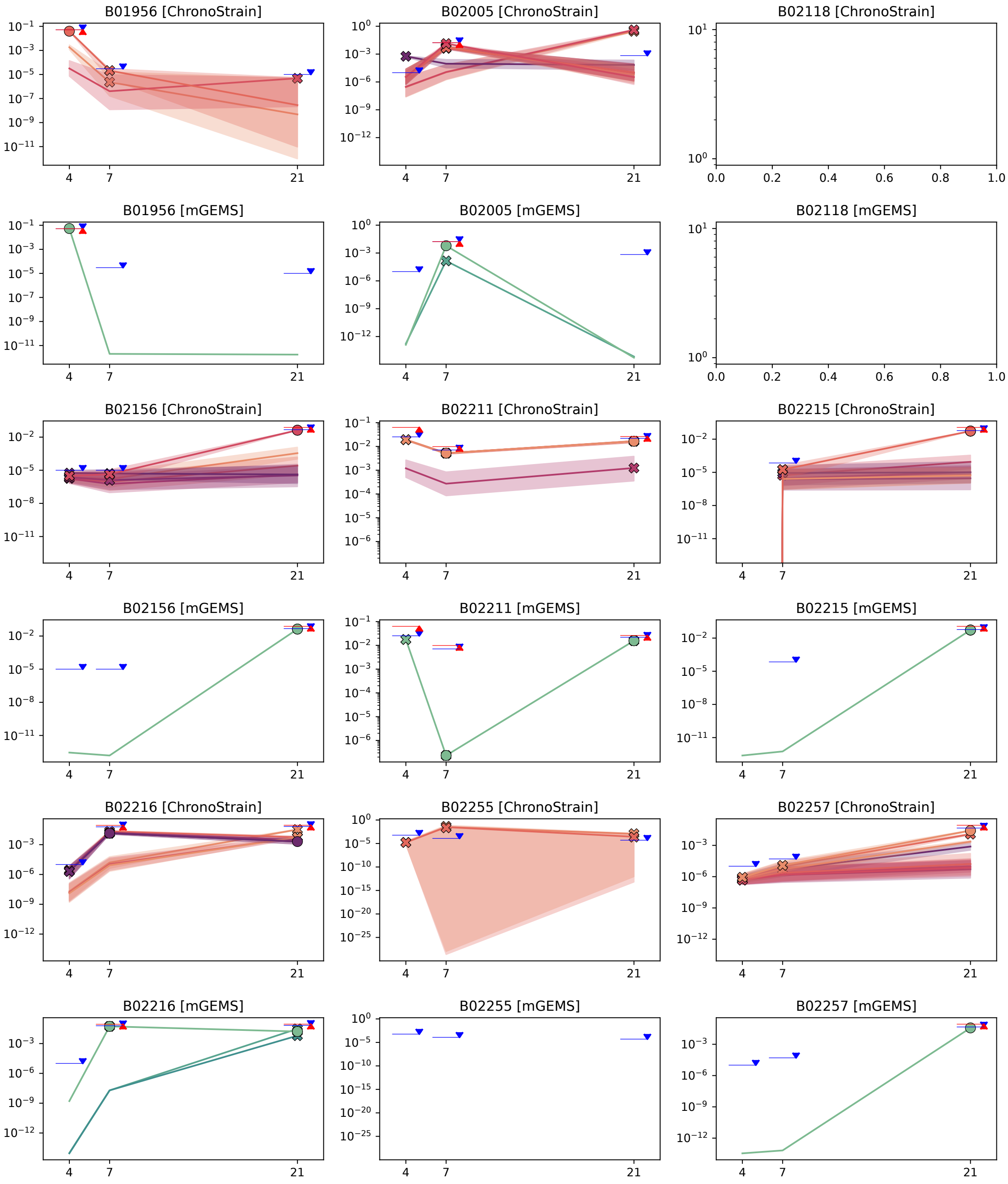


Figure E14: ELMC results (part 14 of 21)

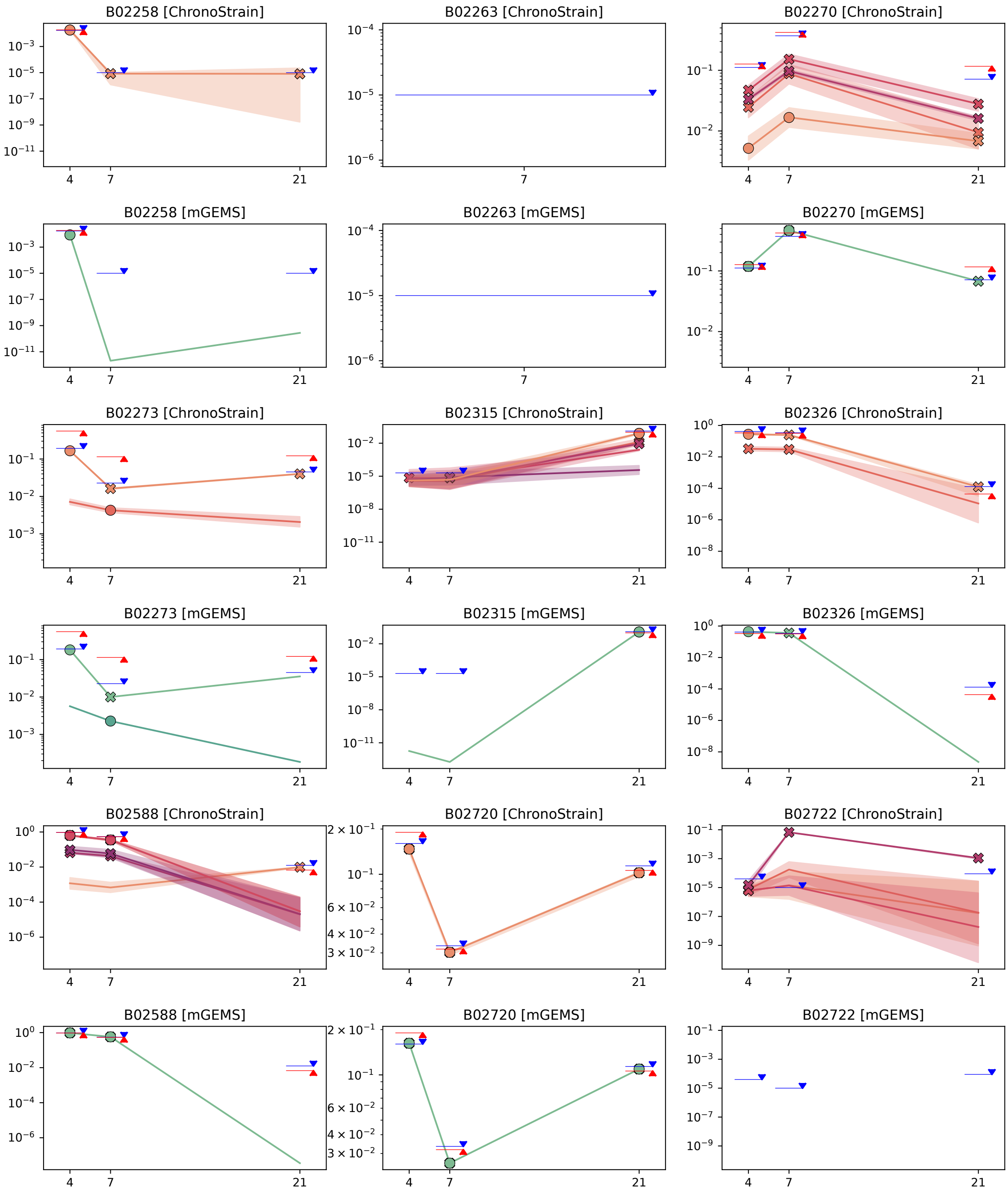


Figure E15: ELMC results (part 15 of 21)

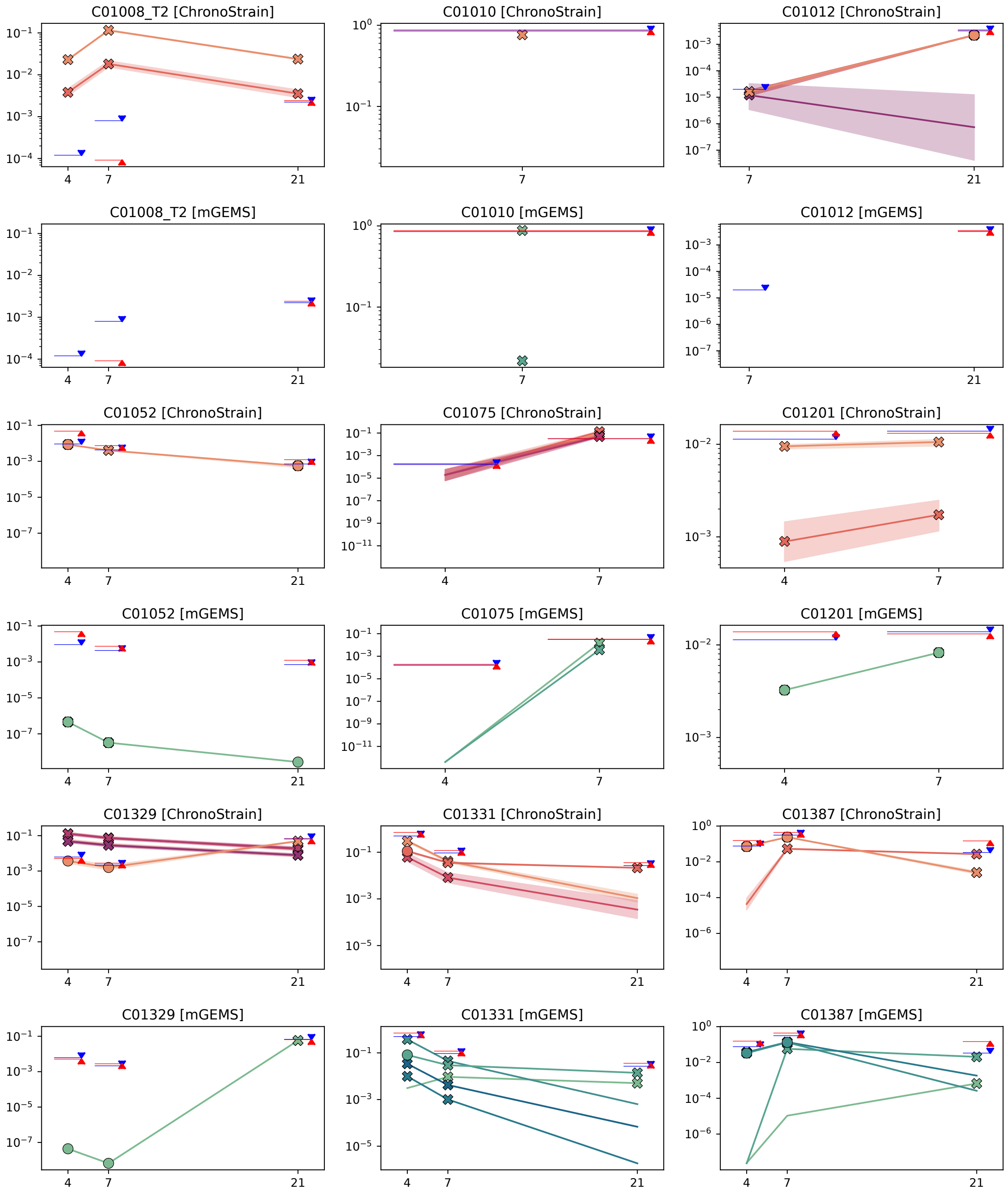


Figure E16: ELMC results (part 16 of 21)

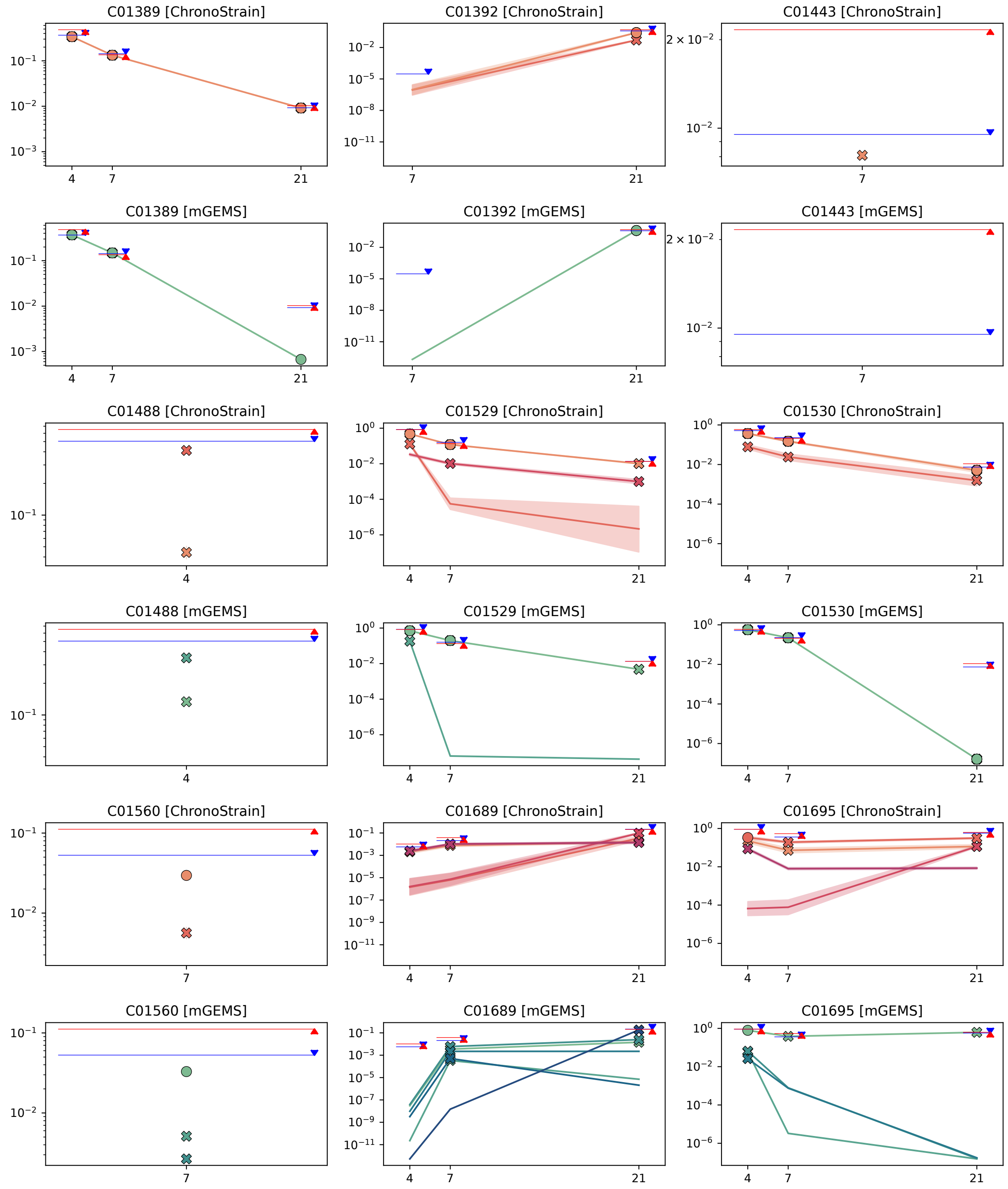


Figure E17: ELMC results (part 17 of 21)

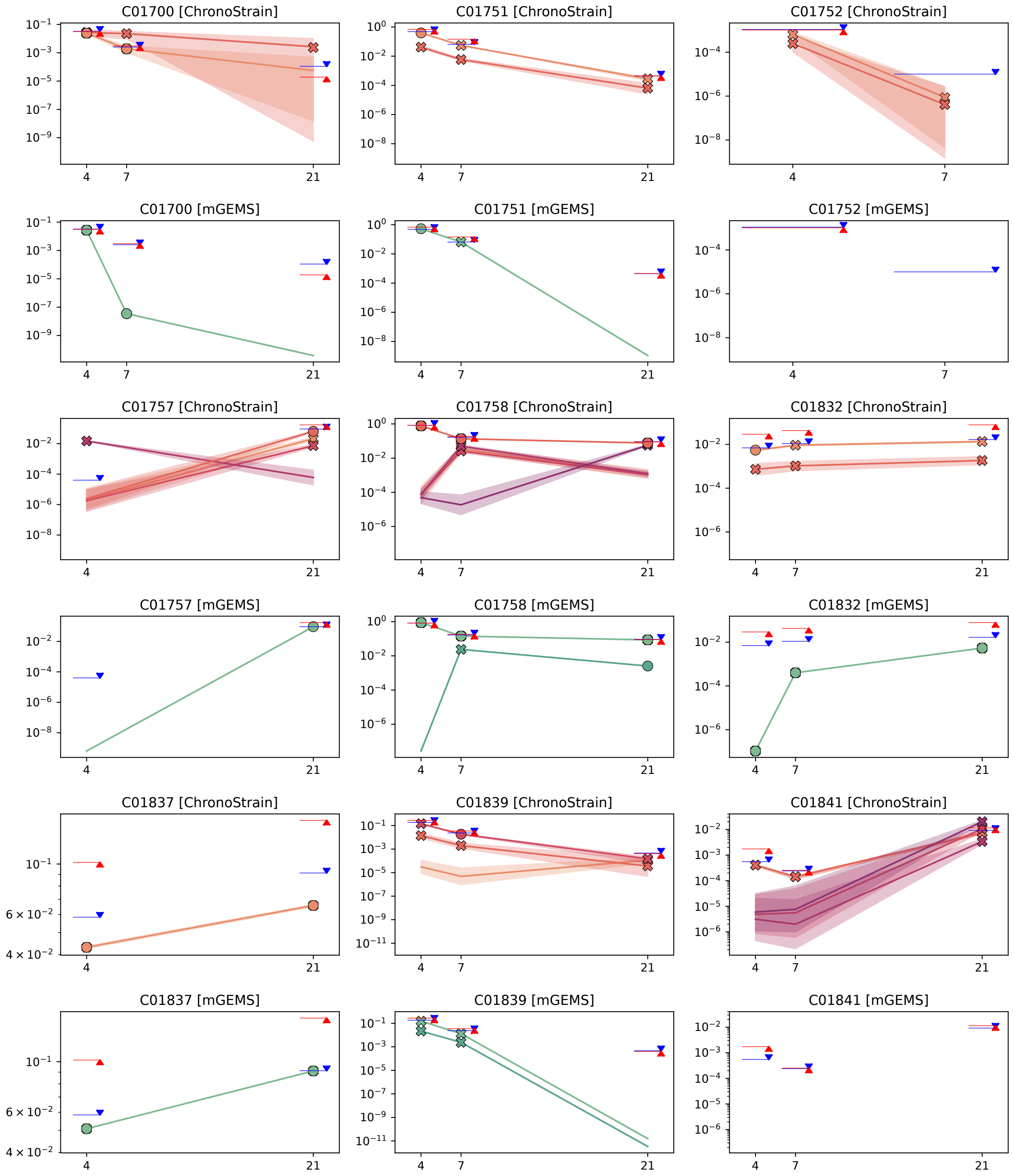


Figure E18: ELMC results (part 18 of 21)

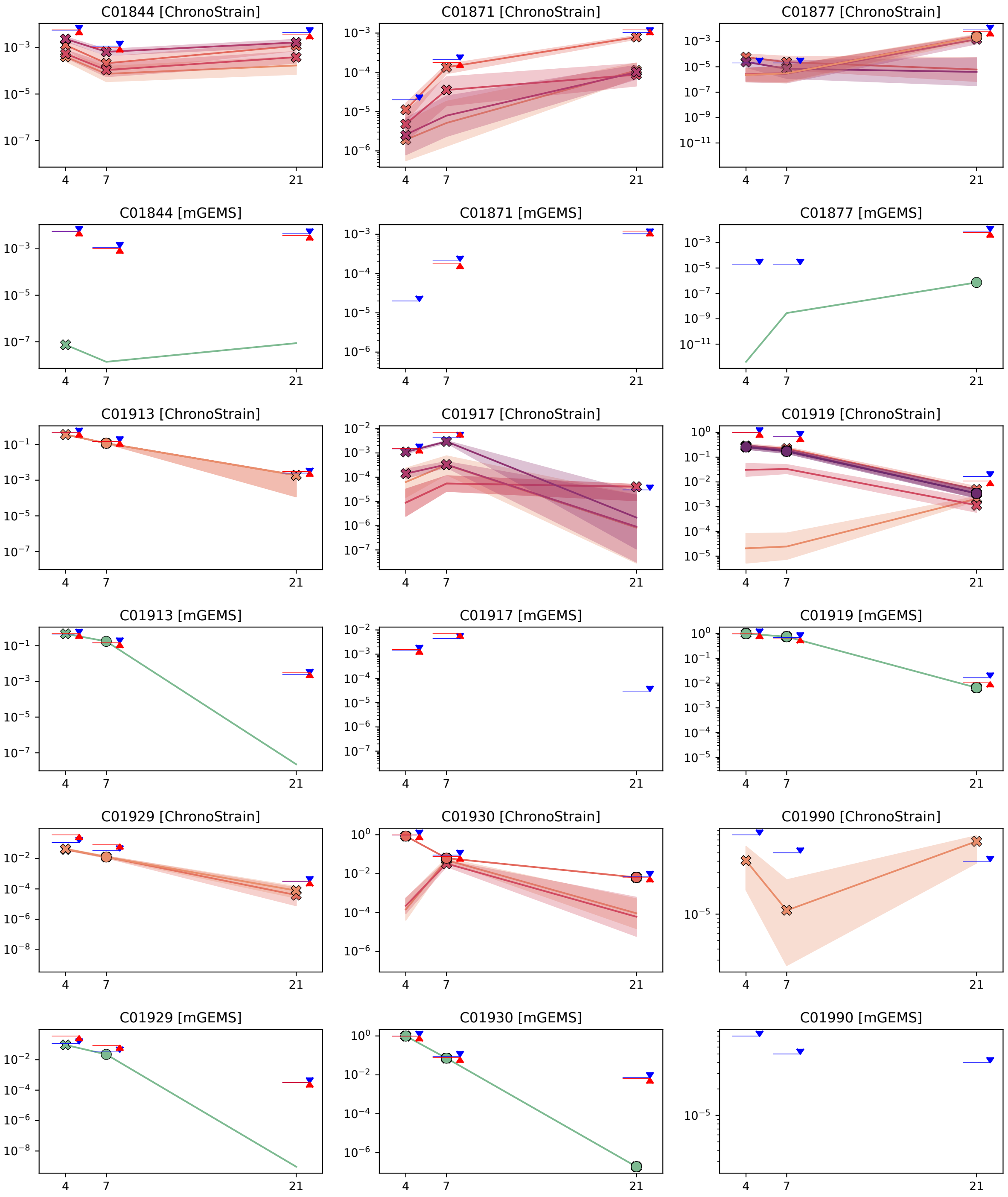


Figure E19: ELMC results (part 19 of 21)

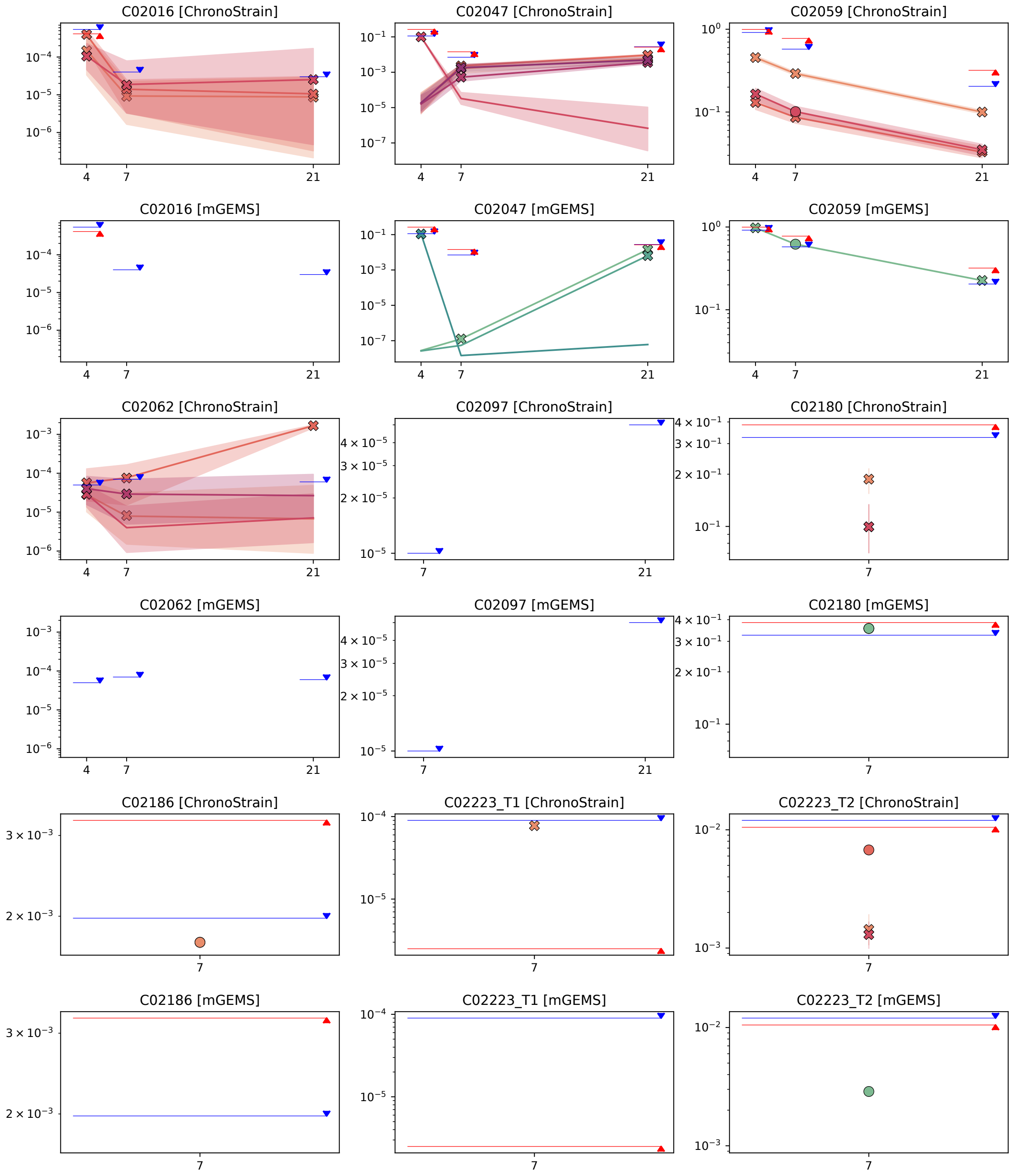


Figure E20: ELMC results (part 20 of 21)

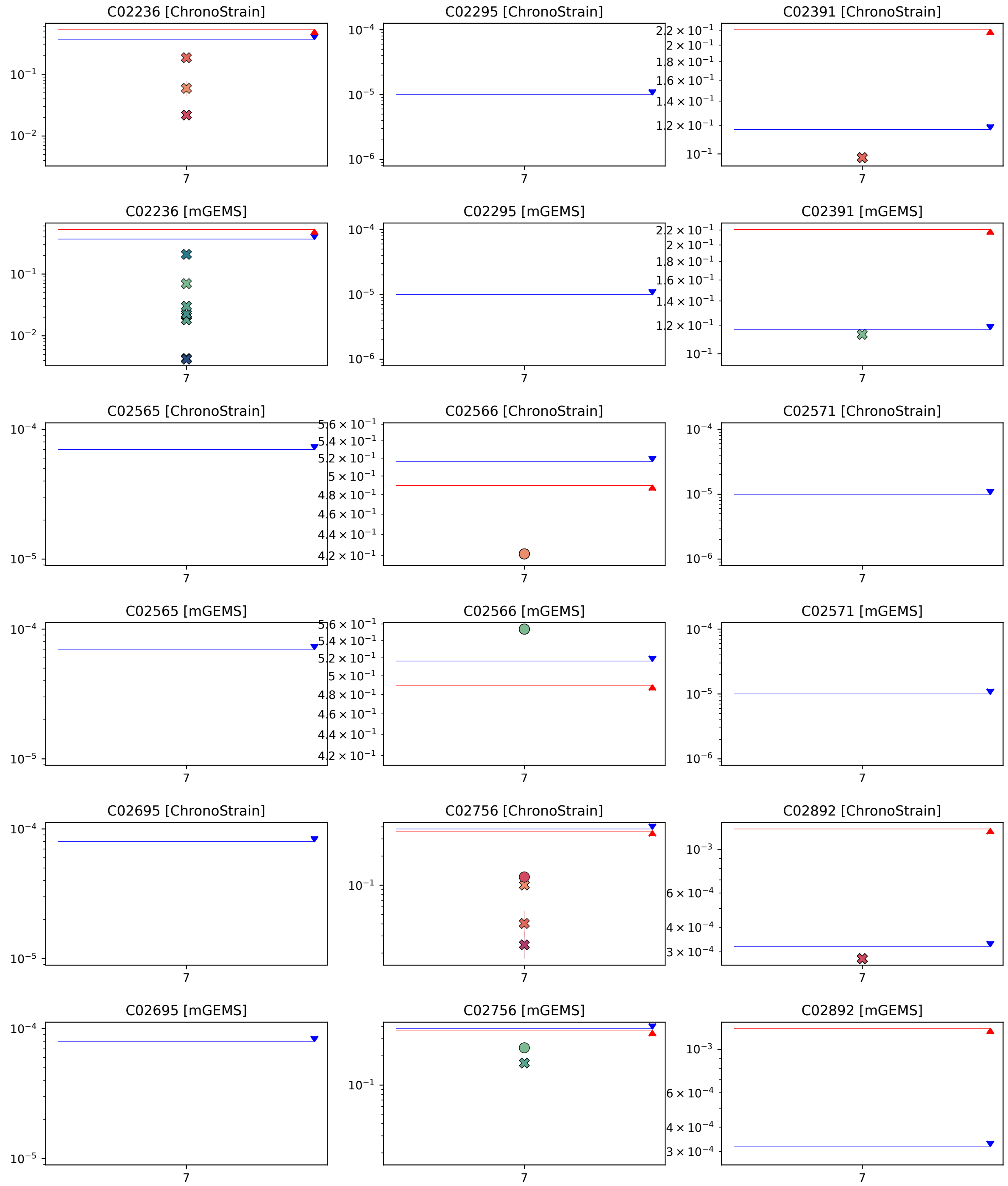


Figure E21: ELMC results (part 21 of 21)
